# Plexin-B2 is a key regulator of cell mechanics during multicellular organization

**DOI:** 10.1101/792077

**Authors:** Chrystian Junqueira Alves, Rafael Dariolli, Theodore Hannah, Robert J. Wiener, Nicolas Daviaud, Rut Tejero, G. Luca Gusella, Nadejda M. Tsankova, Rodrigo Alves Dias, José Paulo R. Furtado de Mendonça, Evren U. Azeloglu, Roland H. Friedel, Hongyan Zou

## Abstract

During multicellular organization, individual cells need to constantly respond to environmental cues and adjust contractile and adhesive forces in order to maintain tissue integrity. The signaling pathways linking biochemical cues and tissue mechanics are unclear. Here, we show that Plexin-B2 regulates mechanochemical integration during multicellular organization. In human embryonic stem cells (hESCs), Plexin-B2 controls cell shape and tissue geometry in both 2D epithelial colony and 3D spheroid aggregates by regulating actomyosin contractility and junctional/cell-matrix adhesive properties. Atomic force microscopy (AFM) directly demonstrates that Plexin-B2 modulates cell stiffness in hESC colonies, which in turn impacts cell proliferation and cell fate specification through β-catenin signaling and YAP mechanosensing. YAP also functions as a mechanoregulator downstream of Plexin-B2, thus forming a mechanochemical integrative loop. In human neuroprogenitor cells (hNPCs), Plexin-B2 similarly controls cell stiffness and tensile forces, as revealed by AFM and FRET tension sensor studies. Strikingly, Plexin-B2-deficient hNPCs display accelerated neuronal differentiation. From an organogenesis perspective, Plexin-B2 maintains cytoarchitectural integrity of neuroepithelium, as modeled in cerebral organoids. On a signaling level, Plexin-B2 engages extracellular as well as intracellular Ras-GAP and RBD domains for mechanoregulation through Rap and Rac GTPases. Our data unveil a fundamental function of Plexin-B2 for mechanochemical integration during multicellular organization, and shed light on the principle of force-mediated regulation of stem cell biology and tissue morphogenesis.

## INTRODUCTION

Multicellular organization relies on force-mediated processes ^1,2^. During tissue morphogenesis, there are constant cellular rearrangements that require rapid cytoskeletal remodeling and mechanical adjustment in response to environmental cues ^3,4^. Cell generated forces are transmitted between cells and with the extracellular matrix (ECM), which provide important signals to inform cell fate decisions ^5^. While mechanotransduction pathways that allow cells to perceive and adapt to physical environment are better understood ^1,6^, how cells translate biochemical cues into physical forces remains unclear.

In neurodevelopment, neural tube closure and ventricular formation exemplify complex mechanomorphogenetic processes. How neuroprogenitor cells (NPCs) orchestrate three key mechanoelements ^7^– i.e. actomyosin contraction, cell-cell junction, and cell-matrix adhesion–to maintain tissue tension and tissue cohesiveness in the developing neuroepithelium is poorly understood.

Plexin-B2 is a member of B-class plexin axon guidance receptors that are activated by class 4 semaphorin ligands ^8^. Plexin-B2 deletion in mice results in neural tube closure defect ^9-11^. Plexin-B2 also regulates cerebellar granule cell migration, corticogenesis, as well as proliferation and migration of neuroblasts in postnatal rostral migratory stream ^9,10,12,13^. Besides neurodevelopment, plexins critically regulate cellular interactions in a wide range of contexts, including vascular development, immune system activation, bone homeostasis, as well as renal epithelial morphogenesis and repair ^14-23^. However, a basic mechanism unifying the multifaceted roles of Plexin-B2 in diverse tissue types is lacking.

In a recent phylogenetic analysis of plexin/semaphorin evolution, we found that plexins most likely emerged in an ancestral unicellular organism more than 600 million years ago, predating the appearance of nervous system ^24^. The domain structures of plexins are highly conserved throughout all clades, including extracellular ring structure ^25,26^ and intracellular Ras-GAP (GTPase activating protein) domain. The Ras-GAP domain of plexins can inactivate small G proteins of the Rap1/2 families ^27^, which have a pleiotropic network of effectors to regulate cytoskeletal dynamics ^28^ and spatial-temporal control of cell adhesion ^29^. Hence we speculated that during the transition from unicellular to multicellular organisms, plexins may have evolved as a key mechanoregulator to counterbalance adhesive forces from other ancient molecules such as cadherins and integrins.

Here, we demonstrated that Plexin-B2 dictates cell shape and tissue geometry in hESC colony by regulating actomyosin contractility while adjusting junctional and cell-matrix adhesive forces. Atomic force microscopy (AFM) revealed cell stiffness and surface topology of hESC as a function of Plexin-B2 activity. This in turn, impacted proliferation and cell fate of hESC through β-catenin signaling and mechanosensing activity of Yes-associated protein (YAP). YAP also functions as a downstream effector of Plexin-B2 to regulate cell mechanics, thus forming a mechanochemical integrative loop. In hESC-derived hNPC, Plexin-B2 similarly controls cell stiffness and cell differentiation. Plexin-B2 is also essential for cytoarchitectural integrity of neuroepithelium in the developing cerebral organoids. Structurally, Plexin-B2 requires its extracellular domain and engages Ras-GAP and RBD domains for mechanoregulation through Rap and Rac small GTPases. In summary, our study deepens the appreciation of force-mediated regulation of stem cell physiology and tissue morphogenesis through Plexin-B2, YAP, β-catenin, and Rap/Rac axis.

## RESULTS

### Plexin-B2 regulates hESC colony geometry and growth kinetics

To study the role of Plexin-B2 in multicellular organization, we generated CRISPR/Cas9-mediated *PLXNB2* knockout (KO) in hESC (Figure 1A). Three clones of wild type (WT) or *PLXNB2* KO hESC were selected, and sequencing showed bi-allelic frameshift or in-frame deletional mutations in KO clones (Figure S1A). Western blotting (WB) confirmed loss of mature Plexin-B2 (170 kDa) in KO cells, while a precursor form (240 kDa) remained detectable (Figure S1B), but presumably nonfunctional due to misfolding induced by in-frame mutations. Normal karyotypes were verified for both WT and KO clones (Figure S1C). Introducing CRISPR-resistant *PLXNB2* restored the mature form in KO cells, while overexpression (OE) was achieved by lentiviral expression of *PLXNB2* in WT cells (Figure S1B). Immunocytochemistry (ICC) revealed wide expression of Plexin-B2 in WT hESCs, but absent or further enhanced in KO or OE cells, respectively (Figure 1B). Sema4C, a putative ligand, was also abundantly expressed in hESCs (Figure S1D).

**Figure 1.**
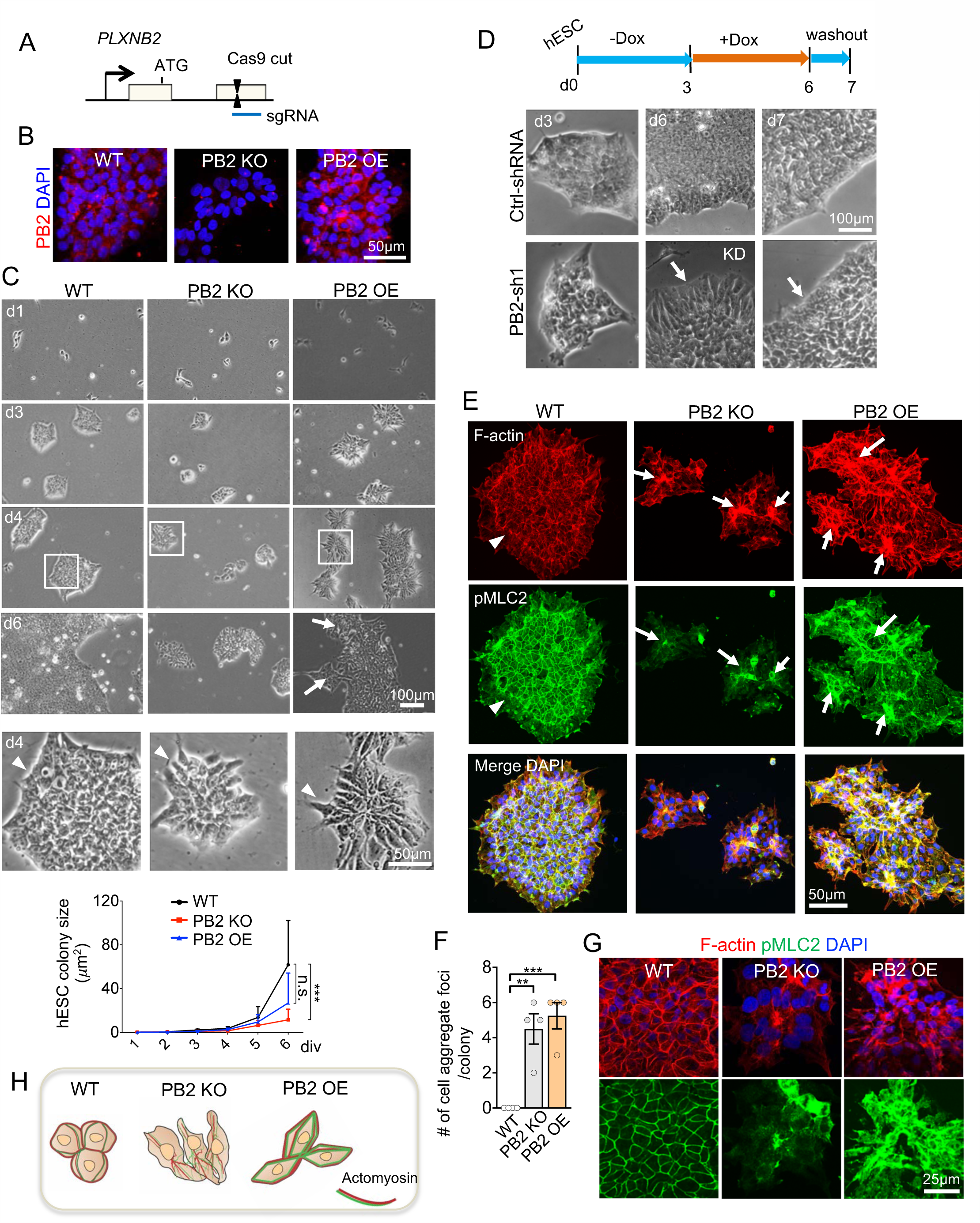
Plexin-B2 regulates colony geometry and growth kinetics of hESCs in adherent culture. (A) Schematic depiction of small guide RNA (sgRNA) targeting *PLXNB2* exon 3 for CRISPR/Cas9-mediated knockout. (B) ICC images show expression of Plexin-B2 (PB2) in wildtype hESCs (clone WT#1), and absence or enhanced expression in *PLXNB2* KO (clone KO#1) or OE hESCs, respectively. (C) Time-course phase contrast images and quantitation of hESC colony formation. After fresh passage, the initial growth kinetics of WT (clone WT#1), *PLXNB2* KO (clone KO #1) or *PLXNB2* OE hESCs were comparable, but by day 4-6, *PLXNB2* KO hESC colonies were significantly smaller. Arrows point to cell aggregation foci within OE colonies. Enlarged images of boxed areas at d4 are shown in bottom panel. Arrowheads point to cell morphology at colony edge. Data are presented as mean ± SEM. Two-way ANOVA followed by Dunnett’s post hoc test. n=8-10 per group. For Genotype: *F*_2, 269_= 4.033; for Time course: *F*_5, 269_= 5.95; for Interaction: *F*_10, 269_= 2.21. * * * p<0.001; n.s., not significant. (D) Top, experimental scheme of doxycycline (Dox)-inducible shRNA knockdown (KD) of *PLXNB2* during hESC colony formation. Bottom, phase contrast images of hESC colonies. Three days after *PLXNB2* KD (d6), cells displayed elongated morphology at colony edge (arrow). 24 hr after Dox washout (d7), cells returned back to rounded morphology as in control (arrow). (E-F) IF images of hESC colonies stained for F-actin (phalloidin) and pMLC2 show differences in cellular organization, colony geometry, and actomyosin network. DAPI for nuclear staining. Arrowheads point to actomyosin band at colony periphery in WT colony, which was disrupted in both mutants. Arrows point to cell aggregation foci within mutant, but not WT colonies. Graphs in (F) show mean ± SEM. One-way ANOVA followed by Dunnett’s multiple comparisons test versus WT. n=4 fields. *F*_2, 9_= 18.43. * * p<0.01; * * * p<0.001. (G) Confocal images highlight cobblestone-like cortical actomyosin network in WT hESC colony, which was disrupted in *PLXNB2* KO or OE colonies. (H) Schematic depiction of cell morphology, actomyosin network, and colony geometry as regulated by Plexin-B2 during self-organization of hESCs. Also see Figures S1 and S2.

We first examined the impact of *PLXNB2* KO or OE on self-organization of hESCs into adherent epithelial colonies. Unlike mouse ESCs, which mirror inner cell mass of blastocysts, hESCs recapitulate the developmentally later epiblast stage and form flat epithelium-like colonies with apico-basolateral polarization, stereotypical E-cadherin-based adherens junctions, ZO-1-containing tight junctions, as well as integrin-based matrix adhesion ^30,31^. After fresh passage at low density, the initial growth kinetics of individual hESCs appeared comparable in all three groups, but by day 4-6, colonies of *PLXNB2* KO cells appeared much smaller than WT; by comparison, *PLXNB2* OE hESCs initially exhibited diminished propensity for self-assembly, but once cell-cell cohesion took place, the colony expanded at similar speed as WT (Figure 1C). Closer inspection revealed that *PLXNB2* KO colonies displayed distinct colony geometry with less smooth contour due to altered cell morphology and cellular alignment at colony periphery (Figure 1C). The same phenotype was observed in all three clonal KO lines (Figure S1E). Conversely, *PLXNB2* OE colonies exhibited spiky contours with individual cells assumed a more angular appearance at both colony edge and interior (Figure 1C).

To gain temporal control, we introduced two independent doxycycline (Dox)-inducible shRNAs for conditional knockdown (KD) of *PLXNB2*, confirmed by WB (Figure S2A). After fresh passaging without Dox, hESCs self-organized into small colonies in a similar fashion for both shRNA-control and -*PLXNB2*, but introduction of Dox resulted in altered colony geometry and cell morphology at colony edge (Figures 1D and S2B), similar to KO phenotypes. Re-expression of *PLXNB2* after Dox washout reversed the morphological changes within 24 hours (Figure 1D), indicating cytoskeletal dynamics as governed by Plexin-B2.

### Plexin-B2 mediates contractile and adhesive properties of hESC colony

Within hESC colony, earlier traction force studies have revealed the presence of different mechanical niches at different spatial locations, with in-plane traction forces primarily localized at colony edge pointing inward toward colony interior ^5^. We thus postulated that Plexin-B2 might a key regulator of the mechanical interaction between hESCs during colony formation. As tissue tension is generated primarily by actomyosin contraction and initiated by phosphorylated myosin light chain 2 (pMLC2) ^31,32^, we first inspected spatial patterns of filamentous actin (F-actin) and pMLC2 by phalloidin staining and ICC, respectively. In WT hESC colonies, we observed a prominent circumferential actomyosin band spanning multiple cells at colony periphery, and a uniform cobblestone-like cortical actomyosin network in colony interior; both were severely disrupted with Plexin-B2 perturbation (Figures 1E-1H and S2C): in the case of *PLXNB2* KO or KD, cells rearranged into small clusters with actomyosin shifted from colony periphery to colony center, signifying lower traction forces at colony edge; in the case of *PLXNB2* OE, actomyosin accumulated at multiple foci within the colony, cells assumed a tensed appearance, and colony geometry displayed spiky contours. When WT hESCs were exposed to increasing concentrations of SEMA4C, colony geometry displayed a hypercontractile appearance with irregular contours (Figure S2D), resembling the tensed appearance of *PLXNB2* OE colonies.

In multicellular organization, in order to maintain tissue cohesion, contractile forces need to match adhesive forces between cell-cell and cell-matrix in a mechanochemical feedback loop ^33^. Indeed, in WT, we observed a uniform cobblestone-like junctional recruitment of E-cadherin and ZO-1; both were severely diminished in *PLXNB2* KO or KD colonies, but further enhanced in *PLXNB2* OE colonies, albeit in spatial disarray (Figures 2A and S3C). We next examined matrix attachment in hESC colonies. In Drosophila sensory neurons, PlexB forms a complex with integrin β subunit to regulate self-avoidance and tiling of dendritic arbors through surface stabilization of integrin ^34^. Analogously, in WT hESCs, confocal microscopy revealed juxtaposition of Plexin-B2 and activated integrin β1 on cell surface; both were markedly reduced or enhanced in *PLXNB2* deficient or OE cells, respectively (Figure S3A). Consistently, phosphorylated focal adhesion kinase (pFAK) and paxillin, two key focal adhesions (FA) components, were evenly distributed in WT colony, but more abundant albeit in disarray in *PLXNB2* OE, and diminished and redistributed to the center of cell clusters in *PLXNB2* KO, similar to actomyosin patterns (Figure 2B). Notably, only the distribution but not the total protein levels of integrin β1 and pFAK were affected (Figure S3B). Hence, Plexin-B2 orchestrates both traction and adhesive properties in hESC colony.

**Figure 2.**
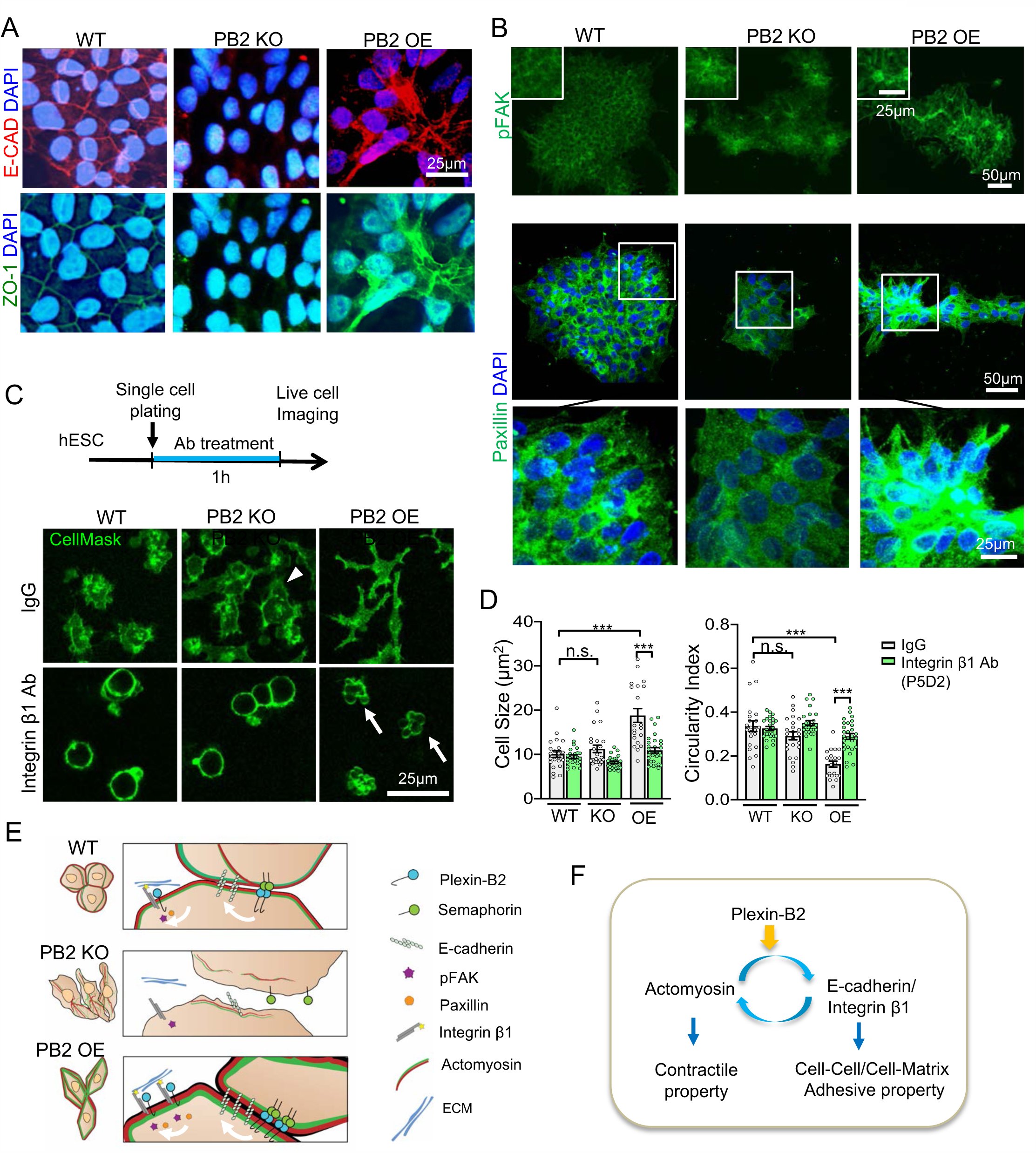
Plexin-B2 regulates tissue mechanics of hESC colony by tuning the mechanochemical feedback loop. (A) Confocal ICC images show alteration of junctional recruitment of E-cadherin and ZO-1 in mutant hESC colonies. (B) Confocal ICC images show altered cellular organization and redistribution of FA components pFAK and paxillin in mutant hESC colonies. (C-D) Schematic of experimental design for live-cell imaging 1 hour after single cell plating (C, top panel). Live-cell images show distinct cell morphology of hESCs revealed by CellMask membrane stain when exposed to function blocking antibody against integrin β1 or control IgG (C, bottom panel). Arrowhead points to large cell protrusion from *PLXNB2* KO hESC. Arrow points to wrinkled cell membrane of *PLXNB2* KO hESCs. (D) Graphs show mean ± SEM for cell size and cell roundness (circularity index). One-way ANOVA followed by Tukey’s multiple comparisons test. Data collected from 4-5 fields per condition. *F*_5, 138_= 20.32. * * * p<0.001; n.s., not significant. (E) Schematic model of the effect of Plexin-B2 in orchestrating three key mechanoelements: actomyosin contraction, E-cadherin-based junctional adhesion, and integrin-mediated cell-matrix attachment. (F) Schematic model of a mechanoregulatory role of Plexin-B2 in fine tuning a mechanochemical feedback loop. Also see Figure S3.

### Plexin-B2 dynamically regulates traction forces during hESC colony expansion

In order to examine each component of the mechanochemical feedback loop separately, we dissociated hESCs into single cells and performed live-cell imaging using CellMask fluorescent dye to label cell membranes. Compared to WT, *PLXNB2* KO cells extended larger and highly dynamic cellular protrusions; whereas *PLXNB2* OE cells assumed a branched morphology with large cell size (Figures 2C-D). To perturb cell-matrix attachment, we treated dissociated cells with a function-blocking antibody against integrin β1 before plating, which resulted in cells rounding up for all three groups, but *PLXNB2* OE cells displayed pronounced membrane blebbing, reflecting a hypercontractile state when decoupled from cell-cell and cell-matrix attachment (Figure 2C). Hence, all three mechanoelements are governed by Plexin-B2 and work together to maintain cell morphology (Figures 2E-2F).

We next introduced GFP-tagged β-actin and performed live-cell videography to track cytoskeletal dynamics during the initiation of colony formation (Figure S3D and Movie S1). After low density passage, control hESCs without Dox appeared highly motile with constant repositioning, and cells coalesced into round colonies with even distribution of actin. *PLXNB2* KD cells were even more motile, and colony contours appeared less rounded with actin accumulated more towards the center of cell clusters (Figure S3D). *PLXNB2* OE cells after low density passage exhibited diminished propensity for self-assembly, leading to cell death probably as an indicator of high tension (Movie S1).

The maintenance of colony integrity and cellular association is widely accepted as an indicator of stem cell state for hESCs ^30^. We therefore investigated if loss of stemness might be responsible for Plexin-B2 phenotypes. We detected no overt changes in the expression of stem cell markers such as SOX2, OCT4, and NANOG in either *PLXNB2* KO or OE colonies, even at colony edge where cell morphologies were most affected (Figure S3E).

### Plexin-B2 regulates cell stiffness in hESC colony

To provide direct measurement of cell stiffness, we applied AFM indentation ^35^ in hESC epithelial colonies, which indeed revealed lower cell stiffness in *PLXNB2* KO than WT (2.2 vs. 2.5 kPa), but increased cell stiffness in *PLXNB2* OE colonies (4.0 kPa) (Figures 3A-3B).

**Figure 3.**
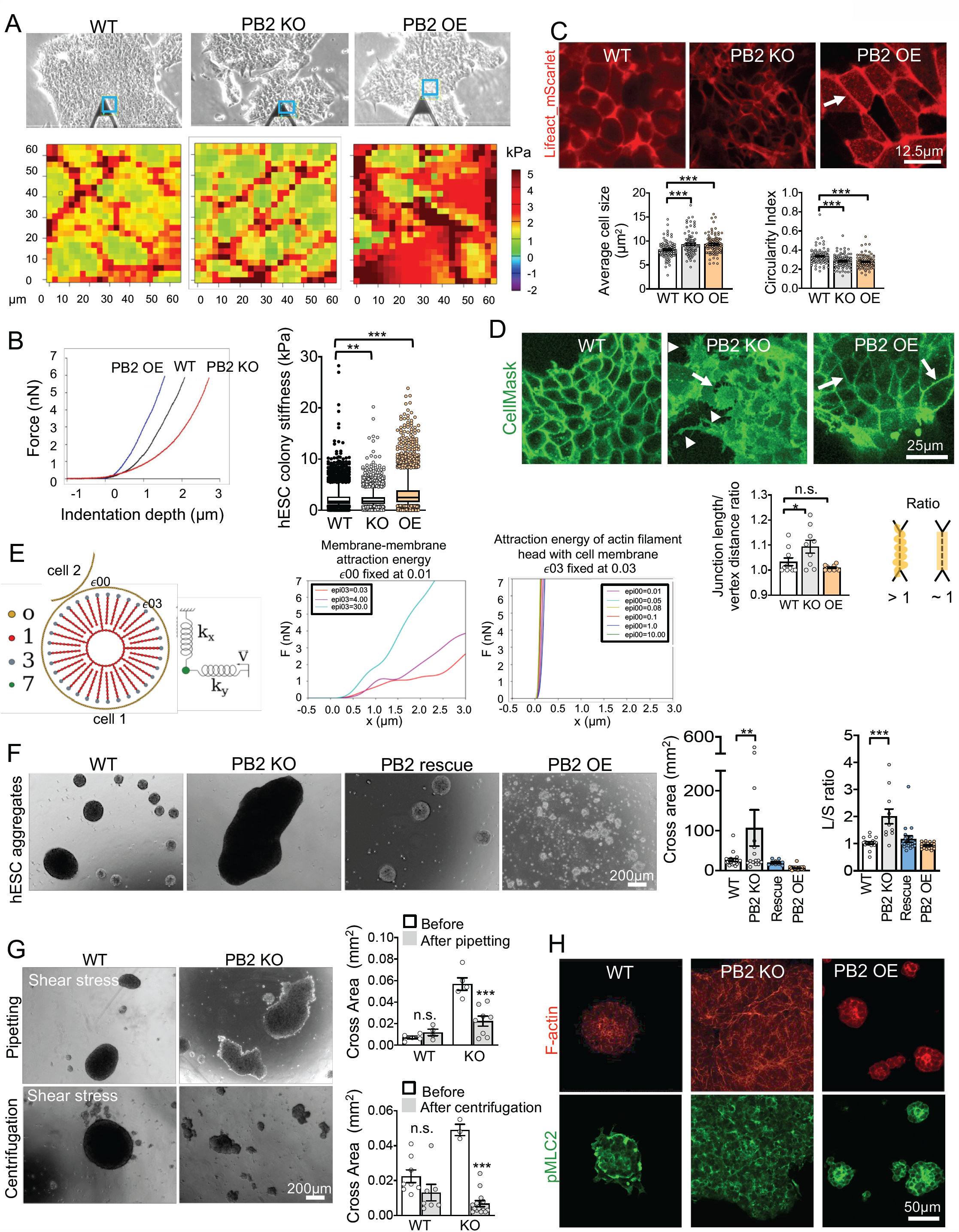
Plexin-B2 regulates cell stiffness and 3D spheroid aggregation of hESCs. (A) Top panel, phase contrast images show areas in hESC colonies (blue squares) measured by AFM. Bottom panel, heat maps of cell stiffness of indicated area. (B) Quantification of AFM indentation measurements (left) and box plots of hESC colony stiffness (right). Box plots show median, 25-75% quantiles, and top and bottom 5% data points. Kruskal-Wallis test followed by Dunn’s multiple comparisons test versus WT. Data collected from 6-7 plates per group. * * p<0.01; * * * p<0.001. (C) Top, live-cell imaging show cortical F-actin in hESC colonies visualized by LifeAct_mScarlet. Arrow points to tensed appearance of cortical F-actin and stretched junction borders in *PLXNB2* OE colony. In contrast, *PLXNB2* KO cells exhibited reduced cortical F-actin and irregular junction borders. Bottom, graphs show mean ± SEM of cell size and cell roundness (circularity index) of indicated hESCs. One-way ANOVA followed by Tukey’s multiple comparisons versus WT. Data collected from 3-5 fields per group. For cell size: *F*_2, 297_= 11.06; for circularity: *F*_2, 297_= 20.15, * * * p<0.001. (D) Top, live-cell imaging shows distinct cell morphology and cell borders in hESC colonies, revealed by CellMask membrane stain. *PLXNB2* KO colony exhibited wavy cell borders (arrow) and prominent cell protrusions (arrowheads), whereas *PLXNB2* OE hESC colony exhibited stretched junction borders (arrow). Bottom, quantification of the ratio of junction length over vertex distance at cell borders (diagram to the right). Data are presented as mean ± SEM. One-way ANOVA followed by Tukey’s multiple comparisons test versus WT. Data collected from 3-5 fields per group. *F*_2, 26_= 6.46. * p<0.05, n.s., not significant. (E) Molecular dynamics mathematical simulation. Left, schematic of structural components of a cell in simulated system (bead-spring model). Beads that make up cell membrane are marked as 0, actin filaments and nuclear membrane as 1, and ends of actin filaments near cell membrane as 3. Tip of AFM probe is depicted as green sphere and marked as 7, with two elastic constants *K*_x_ and *K*_y_ in the x and y directions, respectively. Graphs on right show the force as a function of the distance traveled by the AFM tip, keeping membrane-membrane attraction energy (referred to as *ϵ*00) fixed, while varying the attraction energy of the actin filament head with the cell membrane (*ϵ*03) (left), or vice versa, fixing *ϵ*03, while varying *ϵ*00 (right). (F) Left, images of 3D aggregates of hESC after 48 hr culture in low adherent condition. Right, graphs show mean ± SEM of aggregate size (measured by cross area) and shape (L/S ratio: longest vs. shortest diameter). One-way ANOVA followed by Dunnett’s multiple comparisons test versus WT. Data collected from 3-5 fields per group. For cross area: *F*_3, 77_= 6.53; for L/S ratio: *F*_3, 73_= 16.18. * * p<0.01,* * * p<0.001. (G) Images and quantifications show reduced capability of *PLXNB2* KO hESC aggregates to withstand shear forces of pipetting or low-speed centrifugation as compared to WT. Data represent mean ± SEM. Two-tailed unpaired *t*-test. n=3-5 fields per group. * * * *P*<0.001; n.s., not significant. (H) IHC images of cross sections of 3D hESC aggregates show that *PLXNB2* KO displayed cellular disarray and disorganized F-actin (phalloidin) and pMLC2 networks. *PLXNB2* OE hESC aggregates exhibited more compact actomyosin. Also see Figures S4 and S5.

To better visualize cortical actin filament network in live cells within established hESC colonies, we introduced Lifeact_mScarlet, a fluorescent protein fused with a peptide that specifically binds to F-actin, which importantly does not interfere with actin dynamics ^36^. In *PLXNB2* OE colony, time-lapse live-cell imaging revealed striking angular cell morphology with tensed cortical F-actin, in contrast to the more relaxed appearance of WT cells (Figure 3C and Movie S2). Conversely, *PLXNB2* KO cells accumulated less cortical F-actin, formed ill-defined junctional borders, and frequently extended filopodia-like protrusions at colony edge, reflecting perhaps reduced matrix adhesion (Movie S2). Quantification showed increased cell size and reduced circularity index for *PLXNB2* KO or OE cells relative to WT in epithelial colonies (Figure 3C).

We further examined junctional borders in live hESC colonies at higher resolution using CellMask labeling and live-cell videography (Figure 3D and Movie S3). Strikingly, in *PLXNB2* KO colony, cells displayed wavy junctional borders, and extended frequent filipodia-like protrusions at colony edge; whereas *PLXNB2* OE colony cells exhibited angular shape with straight junctional borders (Figure 3D). We also applied AFM 3D contact imaging to measure cell surface topology in hESC colony, which revealed increased surface roughness for *PLXNB2* KO cells, but smoother surface topology for *PLXNB2* OE cells relative to WT (Figure S4A).

To investigate how Plexin-B2 influences cell shape and colony geometry, we studied the effects of actomyosin inhibitors. When WT hESC colonies were treated with low concentrations of blebbistatin (10µM), which inhibits myosin II, or Y-27632 (20µM), a Rho-associated coiled-coil-containing kinase (ROCK) inhibitor of myosin contractility and actin polymerization ^37^, colony contours became less organized and cells aggregated into small clusters, with increased cellular protrusions at colony periphery, resembling the *PLXNB2* KO phenotypes (Figures S4B-D and Movie S4). Similar findings applied to both *PLXNB2* KO or OE colonies when treated with blebbistatin or ROCK inhibitor, but the effects were more pronounced for KO colonies. Treatment with latrunculin (5µM), which prevents F-actin assembly, resulted in rounding-up of cells, disassembly of cortical F-actin, and cell dissociation in all three groups.

So far, our data demonstrated that Plexin-B2 controls tissue mechanics by tuning actomyosin network in a mechanochemical feedback loop (see Figure 2D). The next question is whether Plexin-B2 primarily regulates actomyosin contractile forces or adhesive forces exerted on cell membrane to control cell stiffness. We performed mathematical simulation of mechanical rigidity of cell membrane using a molecular dynamics (MD) scheme ^38,39^. We first developed a membrane model that could reproduce the main features of the mechanical structures of a cell, based on the coarse-grained bead-spring model for polymers, where cell membrane, nuclear membrane, and actin filaments were modeled as beads connected by springs, while the AFM tip was modeled as a sphere anchored by two springs in x and y directions, respectively (Figure 3E). The proposed model enabled us to probe how cell stiffness might behave if we: i) increase the attraction energy of the actin filament head with the cell membrane (referred to as *ϵ*03), thus simulating actomyosin contractile forces, or ii) increase the interaction energy (adhesion) between neighboring cell membranes (referred to as membrane energy, *ϵ*00), thus simulating adhesive force between cells. We simulated the two scenarios: fixing *ϵ*00 while varying *ϵ*03, or vice versa, fixing *ϵ*03 while varying *ϵ*00 (Figure 3E; Movies S5-6). Results indicated that the first scenario fitted better with our experimental data in that AFM tip needed to make a much smaller force to indent cell membrane, thus supporting the model that Plexin-B2 primarily controls actomyosin contractility to dictate cell stiffness.

### Plexin-B2 regulates tissue mechanics in 3D hESC aggregates

We next examined the role of Plexin-B2 in 3D self-aggregation of hESCs (Figure 3F). In low-attachment culture dishes, WT hESCs spontaneously aggregated into compact spheroids of ∼200-500μm diameter by 48 hours; strikingly, *PLXNB2* KO cells formed larger but irregularly shaped aggregates, measuring up to 1.5mm in the longest axis. Expression of CRISPR-resistant *PLXNB2* rescued the phenotype, thus verifying specificity of the KO phenotype. Conversely, *PLXNB2* OE resulted in more compact aggregates (∼18-60μm in diameter), signifying enhanced contractile state.

Despite their large size, *PLXNB2* KO aggregates exhibited less resistance to external shear forces applied by either gentle pipetting or centrifugation, and easily broke up into smaller, irregularly shaped aggregates with uneven edges (Figure 3G), consistent with reduced tissue tension ^40^. In contrast, WT aggregates were capable of maintaining the compact spheroid shape under shear force stress. As tissue tension is generated primarily by actomyosin contraction ^31^, we observed organized F-actin network at spheroid interior surrounded by a rim of pMLC2 in WT aggregates, which were more compact in *PLXNB2* OE, and highly disorganized in *PLXNB2* KO (Figure 3H). In multicellular organization, actomyosin contraction affects assembly of fibronectin (FN) fibrils in ECM ^41,42^, accordingly, we found organized FN fibrils in a gradient from core to periphery of WT spheroids, which became more compact in *PLXNB2* OE, but in disarray in *PLXNB2* KO (Figure S5A). As an alternative approach to gauge cell-cell cohesion, we placed dissociated hESCs in a hanging drop at high density ^43^, which showed reduced compaction for KO, but more compaction for OE relative to WT cells (Figure S5B).

### Plexin-B2-mediated cell mechanics of hESCs impacts β-catenin and YAP signaling

We next investigated the role of Plexin-B2 on signaling pathways and stem cell behaviors. In hESCs, β-catenin can bind to the intracellular domain of E-cadherin to regulate its function ^44^, and the complex interacts with actin cytoskeleton to mediate cellular adhesion and pluripotency ^30^. We therefore examined subcellular localization of β-catenin. In WT hESCs, like E-cadherin, β-catenin was predominantly membrane bound, in a cobblestone-like pattern, and next to Plexin-B2 (Figures 4A and S5C); however, *PLXNB2* KO or KD led to reduction of junctional β-catenin and redistribution to the center of cell clusters, mirroring the pattern of actomyosin (Figures 4A and S5E). Expression of CRISPR-resistant *PLXNB2* restored the uniform membrane localization of β-catenin, while *PLXNB2* OE led to increased nuclear fraction of β-catenin (Figure 4A). WB confirmed the shift of subcellular localization of β-catenin with Plexin-B2 perturbation (Figure 4B), even though total β-catenin levels were comparable in all three groups (Figure 4C). Subcellular fractionation was verified using histone H3K4me2 for nuclear, GAPDH for cytoplasmic, and Plexin-B2 for membrane fraction (Figure S5D).

**Figure 4.**
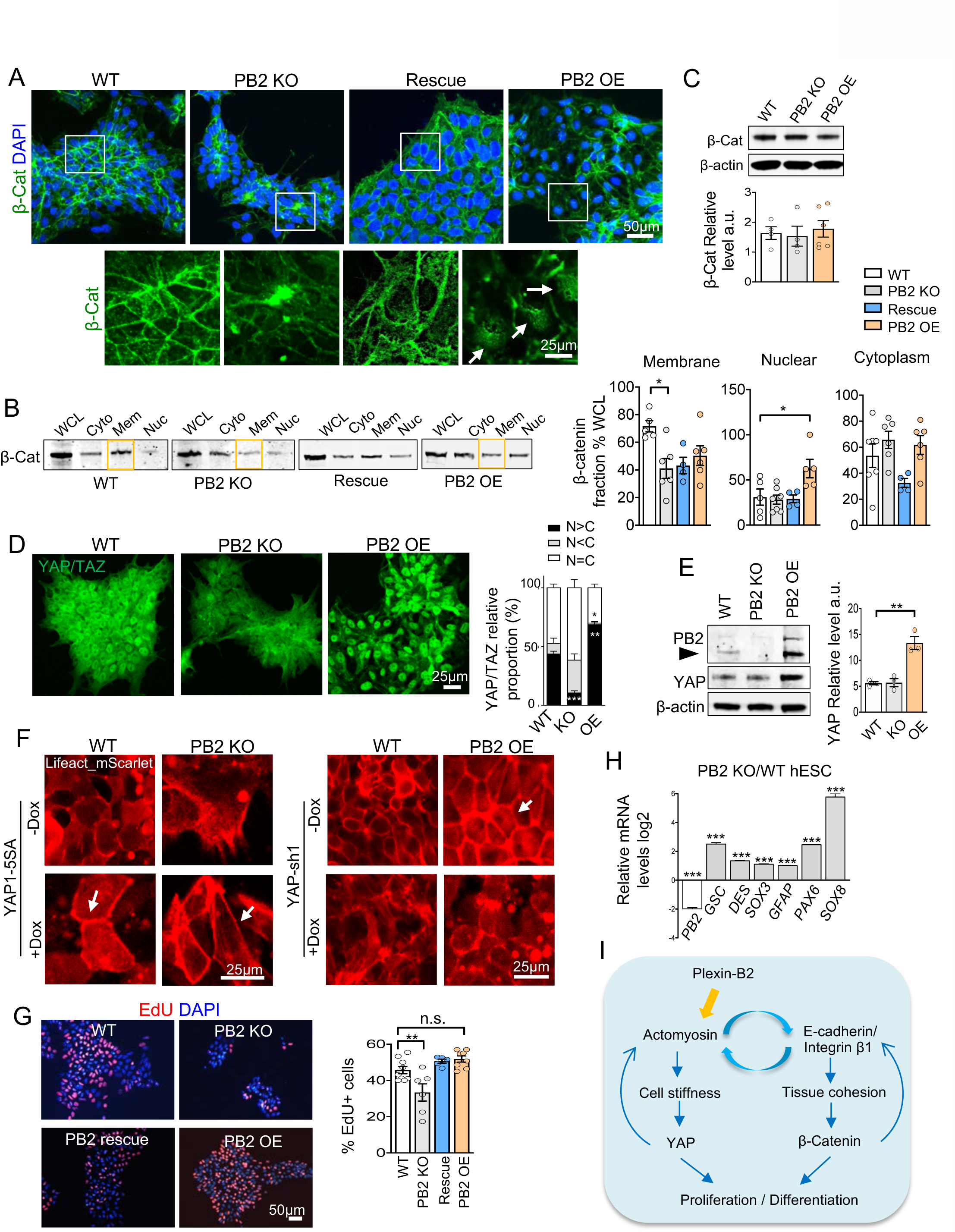
Plexin-B2-mediated regulation of tissue mechanics impacts β-catenin and YAP signaling in hESC. (A) Top, ICC images reveal redistribution of β-catenin in mutant hESC colonies. Bottom, enlarged images of boxed areas highlight membrane localization of β-catenin in WT cells, which was shifted to center of cell cluster foci in *PLXNB2* KO. Arrows point to increased nuclear β-catenin in *PLXNB2* OE cells. (B) WB of cell fractionation show a shift of membrane-bound (Mem) β-catenin (β-cat) to cytoplasm (Cyto) in *PLXNB2* KO hESCs, which was rescued by CRISPR-resistant *PLXNB2. PLXNB2* OE resulted in increased nuclear (Nuc) β-catenin. WCL, whole cell lysate. One-way ANOVA, followed by Tukey’s multiple comparisons test. n=4-7. For Membrane: *F*_3, 17_=4.33; for Nuclear: *F*_3, 17_= 4.85; for Cytoplasm: *F*_3, 19_= 2.93. * p<0.05. (C) WB and quantification show comparable levels of total β-catenin among WT and *PLXNB2* mutant hESC groups. β-actin served as loading control. One-way ANOVA followed by Tukey’s multiple comparisons test versus WT. n=4-6. *F*_2, 11_= 0.19. (D) Confocal IF images show reduced nuclear YAP/TAZ in *PLXNB2* KO hESCs as compared to WT, but a cytoplasmic-to-nuclear shift in *PLXNB2* OE. Quantification shows relative fractions of cells with nuclear YAP/TAZ immunosignals greater, equal, or smaller than cytoplasmic signals (N>C, N=C, N<C). Data presented as mean ± SEM. Two-way ANOVA followed by Dunnett’s post hoc test versus WT. n=3 fields. For Genotype: *F*_2, 6_= 0.47; for Localization: *F*_1.327, 7.964_= 44.32; for Interaction: *F*_4, 12_= 29.07. * p<0.05; * * p<0.01 and * * * p<0.001. (E) WB and quantification show increased level of total YAP in *PLXNB2* OE hESCs as compared to WT or *PLXNB2* KO hESCs. β-actin served as loading control. One-way ANOVA followed by Tukey’s multiple comparisons test versus WT. n=3. *F*_2, 6_= 26. * * p<0.01. (F) Live-cell imaging show cortical F-actin in hESCs visualized by LifeAct_mScarlet, with Dox-inducible expression of YAP1-5SA or shRNA-YAP. Arrows point to tensed cortical F-actin and stretched cellular borders in hESC colonies. Expression of YAP1-5SA in WT or *PLXNB2* KO hESCs phenocopied OE cells with tensed cortical F-actin accumulation and stretched junctional borders. YAP knockdown attenuated the OE phenotypes. (G) IF images and quantification of EdU pulsed cells (30 min) reveal reduced proliferation of *PLXNB2* KO hESCs. One-way ANOVA followed by Dunnett’s multiple comparisons test versus WT, n=5-9 fields. *F*_3, 24_= 9.30. * p<0.05, n.s., not significant. (H) qRT-PCR results show relative transcripts levels of marker genes in *PLXNB2* KO hESCs relative to WT. Two-tailed unpaired *t*-test. n=8. * * * *p*<0.001. (I) Schematic diagram depicting model of Plexin-B2 mediating cellular functions of hESCs by fine-tuning the mechanochemical feedback loop, which impacts β-catenin signaling and YAP mechanosensing. Also see Figures S6-S7.

Individual cells respond to surrounding physical environment through mechanosensors such as YAP, a main nuclear executor of the Hippo pathway ^45^. We next asked whether Plexin-B2-mediated cell intrinsic stiffness also impacts YAP mechanosensor activity by examining the subcellular localization of YAP, a widely used primary read-out of YAP activity: in its active form, YAP is unphosphorylated and translocated into nucleus to activate transcription program to promote cell growth, while phosphorylation of YAP results in its inactivation and retention in cytoplasm for degradation ^46,47^. ICC revealed a nucleus-to-cytoplasm shift of YAP/TAZ (transcriptional coactivator with PDZ-binding motif) in *PLXNB2* KO or KD cells relative to WT, and conversely, a cytoplasm-to-nucleus shift in OE cells (Figures 4D and S5E). Consistently, WB showed increased YAP levels in *PLXNB2* OE hESCs as compared to WT (Figure 4E). Hence, similar to the influence of substrate rigidity on YAP subcellular localization in hESCs ^48^, cell intrinsic stiffness also impacts YAP mechanosensor activity, thus providing another important evidence supporting a mechanoregulatory role of Plexin-B2.

To further interrogate the link between Plexin-B2 and YAP signaling, we introduced Dox-inducible expression of YAP1-5SA, a constitutively active form of YAP (Figures 4F and S6A-B). Live-cell imaging showed that in both WT and KO cells, expression of YAP1-5SA resulted in angular morphology, tensed cortical F-actin, and altered colony geometry, reminiscent of *PLXNB2* OE phenotypes (Figures 4F and S6D). Conversely, we introduced into *PLXNB2* OE hESCs two independent shRNAs targeting YAP, which resulted in attenuation of the OE phenotypes (Figures 4F and S6E). Moreover, in WT hESCs, YAP shRNA KD resulted in larger cell size, reduced cortical F-actin, and altered cell morphology and colony geometry, signifying reduced tensile forces. These data indicated that YAP not only functions as a mechanosensor, but also a mechanoregulator downstream of Plexin-B2, thus forming an integrative loop (Figure 4I).

### Plexin-B2-mediated cellular mechanics impacts proliferation and differentiation of hESC

Tissue tension generated in a cohort of cells can profoundly influence individual cell behavior. Indeed, 30 minute pulsing with 5-ethynyl-2’-deoxyuridine (EdU) labeled far fewer cells in S-phase in *PLXNB2* KO than in WT colonies by both ICC and FACS, a phenotype rescued by CRISPR-resistant *PLXNB2* (Figures 4G and S7A). Of note, *PLXNB2* OE did not further elevate the already high proliferative state of hESCs. Cell survival was not overtly affected by Plexin-B2 ablation based on apoptosis marker cleaved caspase 3 (Figure S7B).

Altered epithelial mechanics can ultimately affect stem cell differentiation status, as small groups of cells in *PLXNB2* KO or KD hESC colonies displayed reduced activity of stem cell marker alkaline phosphatase (AP), and induction of a neural stem cell (NSC) marker, PAX6, at the expense of pluripotency factor NANOG (Figure S7C), indicating premature fate commitment to neural lineage. On the other hand, *PLXNB2* OE caused no premature neuronal differentiation (Figure S7C). We performed qRT-PCR array analysis to survey the expression of a larger set of stem cell and differentiation markers in hESCs, and found no significant transcriptional changes between *PLXNB2* KO and WT cells for ∼90% of the markers (78 out of 89), including 26 ESC markers (out of 28), e.g., *OCT4, NANOG*, and *SOX2* (Figure S7D). The remaining 12% of the markers (11 out of 89) that displayed transcriptional changes were all upregulated in *PLXNB2* KO relative to WT cells, including markers for all three germ layers: e.g., *GSC* (endoderm), *DES* (mesoderm), and *SOX3* (neuroectoderm), as well as neural markers, e.g. *GFAP, PAX6* and *SOX8* (Figures 4H and S7D).

### Plexin-B2 controls actomyosin contractibility and cell stiffness in hNPCs

Given the critical role of Plexin-B2 in neurodevelopment, we next asked whether Plexin-B2 also controls biomechanics of NPCs. To this end, we derived hNPCs from WT, *PLXNB2* KO or OE hESCs and analyzed actomyosin network (Figure 5A). As in hESCs, both F-actin and pMLC2, as well as FA components Paxillin and Vinculin, were markedly reduced in hNPCs with *PLXNB2* KO, but increased with *PLXNB2* OE, even though pFAK levels were not significantly changed in all three groups (Figures 5B and S8A). Confocal microscopy revealed juxtaposition of Plexin-B2 and activated integrin β1 on cell surface of WT hNPCs, both diminished in *PLXNB2* KO hNPCs (Figure S8B).

**Figure 5.**
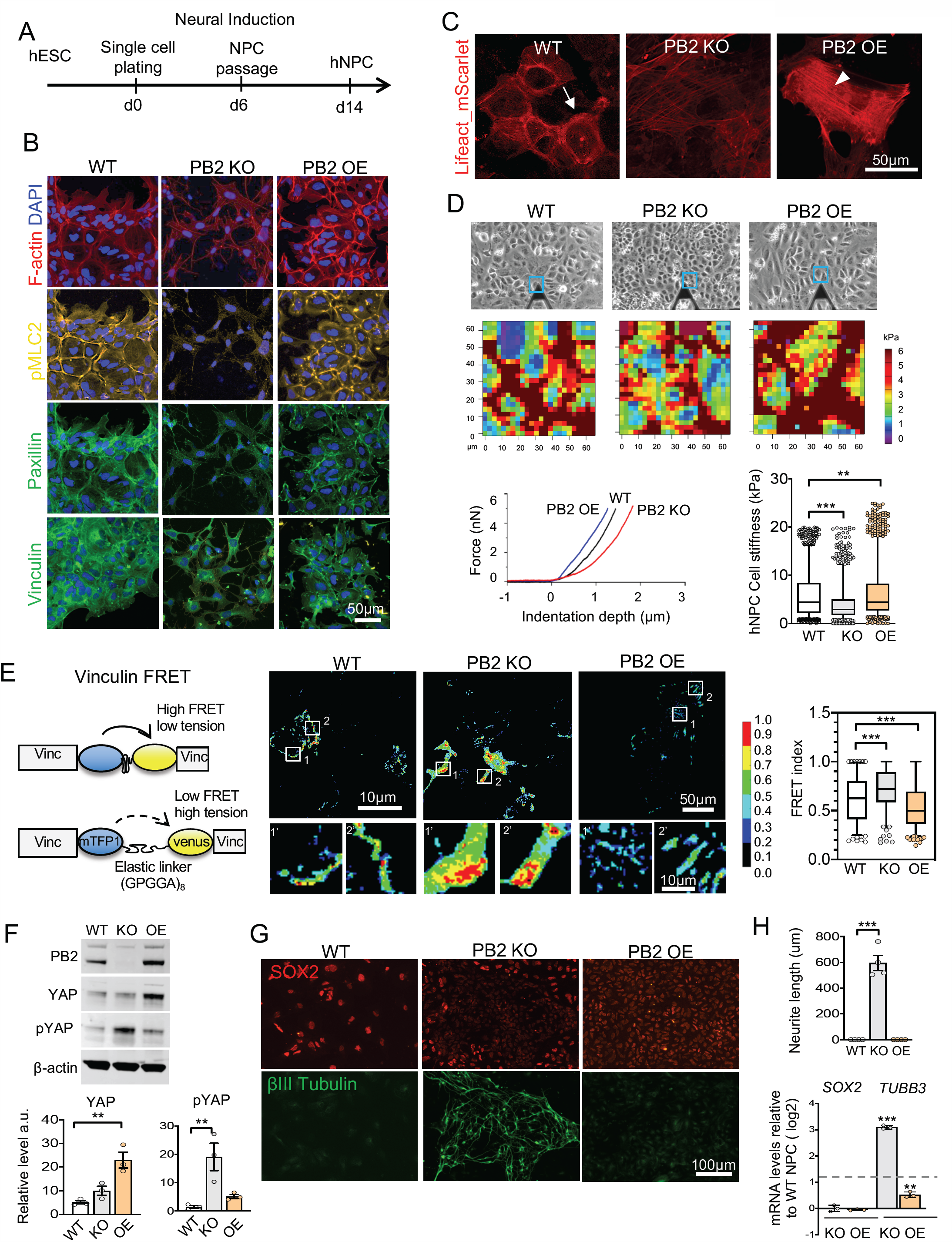
Plexin-B2 regulates cell mechanics and neuronal differentiation of hNPC. (A) Experimental scheme of neural induction in dissociated hESC culture. At day 6, hNPCs were passaged for further maturation. (B) Confocal IF images show differences in F-actin (phalloidin) and pMLC2 network, as well as FA components paxillin and vinculin in hNPCs. (C) Live-cell imaging of hNPCs shows different F-actin patterns in indicated hNPCs as revealed by Lifeact_mScarlet. Arrow points to cortical F-actin in WT hNPCs, which was reduced in *PLXNB2* KO cells. Arrowhead points to stress fibers in *PLXNB2* OE hNPCs. (D) Top panel, phase contrast images show areas of hNPC cultures that were measured by AFM (blue squares). Middle panel, heat maps of stiffness of hNPCs. Bottom, graphs of AFM measurements of hNPCs (left) and quantification of cell stiffness (box plots show median and 25-75% quantiles and top and bottom 5% data points). Kruskal-Wallis test followed by Dunn’s multiple comparisons test versus WT. Data collected from 3 culture plates per group. * * p<0.01, * * * p<0.001. (E) Left, diagram of vinculin-based FRET tension sensor (mTFP1 and venus, linked by an elastic linker) to gauge internal tensile forces in hNPCs. Stretched elastic linker of tension sensor gives rise to lower FRET signal. Middle, representative images of normalized FRET signals. Right, quantification of FRET index as box plots (median, 25-75 quantile, and top or bottom 5% data points). Kruskal-Wallis test followed by Dunn’s multiple comparisons test versus WT. Data collected from 10 fields for WT, 5 fields for *PLXNB2* KO and 12 fields for *PLXNB2* OE. * * * p<0.001. (F) WB shows levels of total YAP and pYAP in indicated hNPCs. One-way ANOVA (for YAP, *F*_2, 6_= 16.58; for pYAP, *F*_2, 6_= 10.59) followed by Dunnett’s multiple comparisons test versus WT. n=3. * * p<0.01. (G) IF images show increased expression of β-III tubulin at the expense of SOX2 in *PLXNB2* KO hNPCs derived from hESCs (clone KO#2), indicating premature neuronal differentiation. (H) Top, quantification shows enhanced neurite length as revealed by β-III tubulin immunostaining in *PLXNB2* KO cells as compared to WT or *PLXNB2* OE cells. Data are presented as mean ± SEM. One-way ANOVA followed by Tukey’s multiple comparisons test versus WT. n=4 fields. *F*_2, 9_= 101.5. * * * p<0.001. Bottom, qRT-PCR results show relative transcript levels of *SOX2* or *TUBB3* in *PLXNB2* KO or OE hNPCs relative to WT. Data are presented as mean ± SEM. One-way ANOVA followed by Tukey’s multiple comparisons test versus WT. n=3. For *SOX2*: *F*_2, 6_= 1.08; for *TUBB3: F*_2, 6_= 492.5, * * p<0.01; * * * p<0.001. Also see Figures S8-S9.

Live-cell imaging further highlighted the striking differences in cell morphology and cortical F-actin network between WT and *PLXNB2* KO hNPCs, whereas *PLXNB2* OE cells displayed angular morphology and numerous stress-fibers, visualized by Lifeact_mScarlet (Figure 5C and Movie S7). AFM measurement revealed higher stiffness for *PLXNB2* OE hNPCs (6.3 kPa), and lower stiffness for *PLXNB2* KO compared to WT hNPCs (4.0 vs. 5.9 kPa) (Figure 5D). As a measurement for tensile forces at focal adhesions in hNPCs, we introduced a vinculin-based FRET tension sensor ^49^ (Figure 5E). Indeed, we observed higher FRET index in *PLXNB2* KO hNPCs than in WT (0.71 ± 0.015 vs. 0.61 ± 0.017), indicative of lower tensile forces at focal adhesions; conversely, *PLXNB2* OE resulted in lower FRET index (0.53 ± 0.014), indicative of higher tensile forces (Figure 5E). Concordantly, the hanging drop cell-cell cohesion assay showed less compaction for *PLXNB2* KO, but more compaction for *PLXNB2* OE relative to WT hNPCs (Figure S8C). WB demonstrated increased YAP levels in *PLXNB2* OE hNPCs, but increased pYAP levels in *PLXNB2* KO hNPCs, while total β-catenin levels were similar in all three groups (Figure 5F and S8D). Taken together, as in hESCs, Plexin-B2 activity in hNPCs regulates cell stiffness and contractile forces, which in turn impacts YAP mechanosensing activity.

### Plexin-B2 activity affects proliferation and differentiation of hNPCs

As in hESCs, *PLXNB2* KO in hNPCs also resulted in reduced proliferation, while *PLXNB2* OE did not further enhance the already high proliferative state of hNPCs (Figure S8E). Previous studies showed sensitivity of mesenchymal stem cells (hMSCs) to tissue elasticity during lineage specification, with soft ECM promoting neuronal whereas stiffer ECM promoting muscle differentiation ^50^. We thus examined the impact of Plexin-B2-mediated cell intrinsic stiffness on lineage specification of hESCs. Upon neural induction, using either single-cell plating or embryoid body neural induction protocols (Figures 5A and S8F-G), *PLXNB2* KO cells showed enhanced upregulation of neuroblast marker doublecortin (DCX), as well as neuronal markers β-III tubulin (TUJ1) and NeuN, but reduced expression of NSC markers PAX6 and SOX2, indicating accelerated neuronal differentiation (Figures 5G and S8F-H). qRT-PCR analyses confirmed upregulation of *TUBB3* and a trend of decrease of *SOX2* transcription in *PLXNB2* KO cells relative to WT; *PLXNB2* OE, on the other hand, did not accelerate neuronal differentiation (Figures 5H).

We next tested whether Plexin-B2-mediated cell mechanics would also impact mesenchymal lineage specification for hESCs. To this end, we subjected hESCs to mesoderm induction followed by cardiomyocyte specification and maturation, which gave rise to rhythmically beating cardiomyocytes (Figure S9A and Movie S8). Remarkably, *PLXNB2* KO or KD cells exhibited massive cell death upon mesodermal induction. However, cells with KD induced after cardiac specification did not affect cell survival or cardiomyocyte specification (Figure S9B). hESCs with *PLXNB2* OE also successfully differentiated into cardiomyocytes, although the rhythmic beating appeared slower than WT (Figure S9B and Movie S8). These data suggested an essential role of Plexin-B2 for mechanical adjustment during both neuronal and mesoderm specification.

### Plexin-B2 regulates mechanomorphogenesis and maintains neuroepithelial integrity during cerebral organoid development

To further probe the role of Plexin-B2 during tissue morphogenesis, we derived from hESCs 3D cerebral organoids with forebrain specification ^51,52^. After 6 weeks of culture, with the last 4 weeks embedded in matrigel under rotational condition (Figure 6A), cortex-like structures emerged in WT organoids, containing ventricular zones (VZ) and rudimental cortical plate (CP) (Figure S10A). Immunohistochemistry (IHC) revealed wide expression of Plexin-B2 in both VZ and CP in cerebral organoids, similar to that in 23 week human fetal cortex (Figures S10A-C) and in mouse embryonic cortex ^53^. Notably, outer radial glia (FAM107A+), a progenitor population predominantly seen in primates and responsible for cortical expansion, were found in both organoid and fetal cortex with high levels of Plexin-B2 (Figures S10D). SEMA4B and 4C were widely expressed in cerebral organoids (Figure S10E).

**Figure 6.**
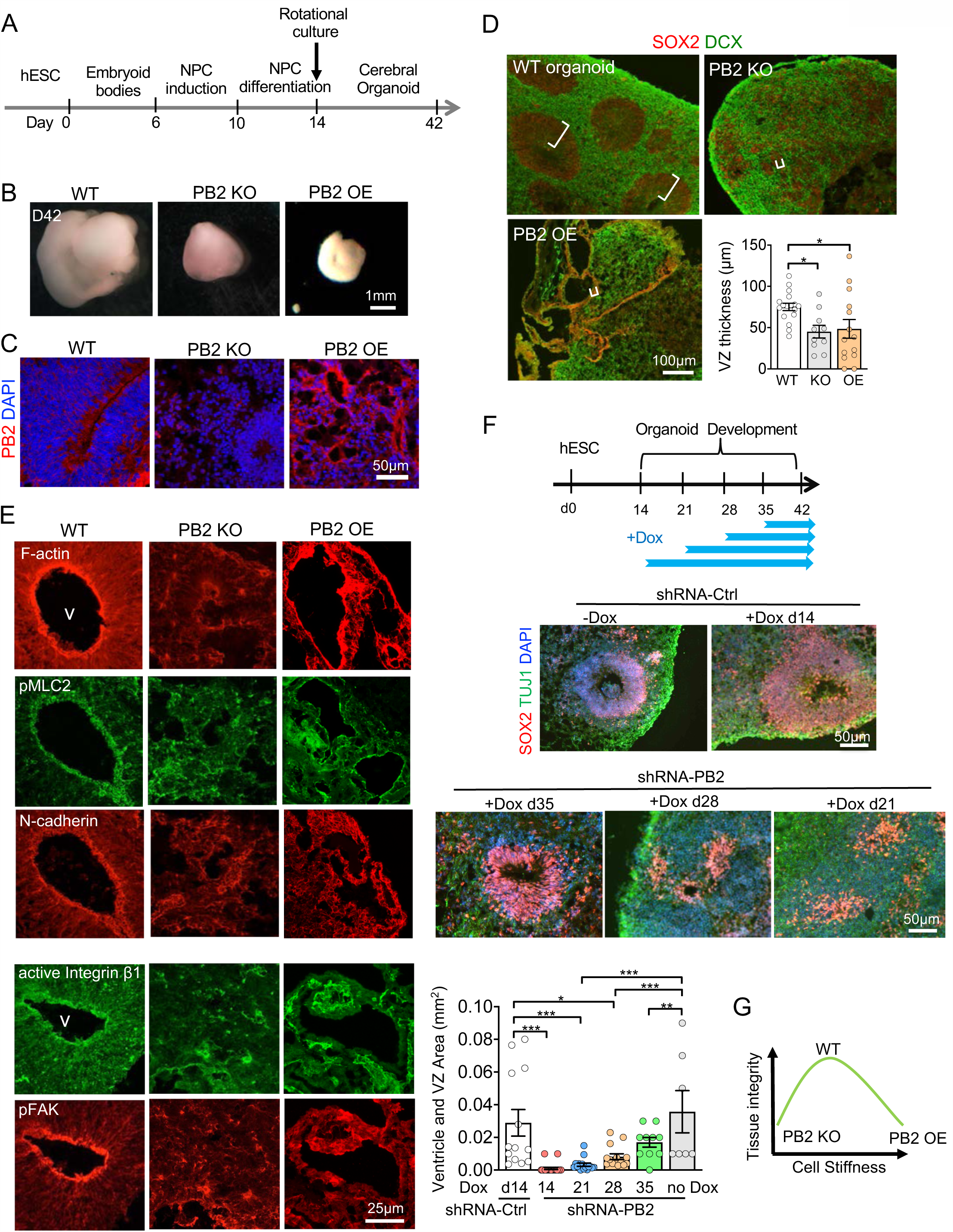
Plexin-B2 regulates neuroepithelium tissue mechanics during cerebral organoid development. (A) Timeline of step-wise cerebral organoid derivation from hESCs. (B) Images show different cerebral organoid sizes with perturbation of Plexin-B2. (C) IHC images show absence of Plexin-B2 in cerebral organoids derived from *PLXNB2* KO hESCs, but increased Plexin-B2 levels in *PLXNB2* OE organoids. Note several developmental anomalies in mutant organoids. (D) IHC images show the presence of SOX2^+^ neuroprogenitors and DCX^+^ neuroblasts in indicated cerebral organoids, which are in disarray in mutants. White brackets denote ventricular zone (VZ) thickness, as quantified in bar graphs showing mean ± SEM. One-way ANOVA followed by Tukey’s multiple comparisons test versus WT. n=10-17, *F*_2, 38_= 4.40. * p<0.05. (E) IF images demonstrate that *PLXNB2* KO or OE organoids (day 42) display disorganized actomyosin network and disarray of cell-cell and cell-matrix junctions indicated by specific markers. Top, schematic diagram of Dox-inducible *PLXNB2* shRNA knockdown at different time points during cerebral organoid development. Middle, IHC images of SOX2^+^ neuroprogenitors and β-III tubulin^+^ immature neurons in cerebral organoids show that earlier KD of *PLXNB2* correlates with more severe phenotypes. Bottom, graphs show mean ± SEM of ventricular area (ventricle plus VZ) in organoids with different timeframe of *PLXNB2* KD. One-way ANOVA followed by Tukey’s multiple comparisons test. n=7-23 fields. *F*_5, 76_= 9.63. * p<0.05; * * p<0.01; * * * p<0.001. (G) Diagram illustrates the importance of balanced Plexin-B2-mediated cell stiffness for tissue integrity during neurodevelopment. Also see Figure S10-S15.

Cerebral organoids derived from *PLXNB2* KO, and more so from *PLXNB2* OE hESCs, were much smaller than matching WT organoids (Figures 6B and S11A-C). Histological analysis by hematoxylin and eosin staining (H&E) revealed severe developmental anomalies in mutant organoids, with multi-layered ventricle-like structures, a defining feature of forebrain specification, completely absent and replaced by numerous simple cysts lacking the VZ; these phenotypes were rescued by CRISPR-resistant *PLXNB2* (Figure S11B). Notably, the cystic structures in *PLXNB2* OE organoids appeared stretched, less spherical with irregular contours (Figure S11B). IHC confirmed Plexin-B2 ablation or overexpression in cerebral organoids and further demonstrated severe disruption of neuroepithelial architecture (Figure 6C).

Remarkably, mutant cerebral organoids contained SOX2^+^ NPCs and DCX^+^ neuroblasts, albeit in disarray (Figure 6D). Thus, *PLXNB2* KO or OE did not completely abolish neural induction per se, but rather affected cell alignment and architectural integrity of neuroepithelium, reflected by marked disorganization of actomyosin networks, N-cadherin at cell junctions, activated integrin β1 and pFAK at FA (Figure 6E).

As structural anomalies in *PLXNB2* KO cerebral organoids may be secondary to altered physiology of hESCs, we switched to Dox-induced *PLXNB2* KD at later stages of organoid development (Figure 6F). IHC confirmed effective *PLXNB2* KD in organoids by Dox (Figure S12A). We found that the earlier the *PLXNB2* KD, the more severe the phenotype, even though the overall sizes of organoids were similar (Figures 6F and S12B). Specifically, when *PLXNB2* KD was initiated at d14, ventricular structures were nearly absent, phenocopying KO; when KD was initiated at d21, only rudimentary cystic structures formed; when KD was initiated at d28 or d35, ventricle-like structures were detected, but they frequently collapsed into smaller cysts. Moreover, in contrast to the stereotypical pattern of FN fibrils in WT organoids, reflective of organized tension forces and cellular alignment, *PLXNB2* KD organoids contained punctate FN patches, whereas *PLXNB2* OE organoids contained more abundant but highly disorganized FN deposits, particularly along the cystic structures (Figure S12C).

### Structural integrity of neuroepithelium affects neuroprogenitor physiology in organoids

The disrupted ventricular cytoarchitecture in *PLXNB2* deficient or OE organoids was associated with decreased proliferation of NPCs, as shown by reduced phospho-Histone3 (M-phase marker), and concordantly, altered localization of YAP/TAZ and β-catenin, even though total level of β-catenin was unchanged (Figures S13A-C). EdU pulse labeled far fewer cells in S phase in *PLXNB2* deficient or OE organoids (Figures S14A-B), while co-labeling with proliferation marker Ki67 revealed increased frequency of cell cycle exit (“quit fraction”), as evidenced by a larger fraction of EdU^+^ Ki67^-^/EdU^+^ cells (Figures S14A-B). Together, these changes resulted in shrinkage of NPC pool, shown by reduced levels of SOX2 in cerebral organoids; however, neuronal lineage progression proceeded in *PLXNB2* KO (albeit in disarray), but severely disrupted in *PLXNB2* OE organoids, which contained fewer β-tubulin III^+^ neurons and CTIP2^+^ lower-layer cortical neurons (Figures S13D-F).

When cerebral organoids were dissociated at day 56 and plated as single cells and cultured for 4 more days without the context of 3D cytoarchitecture, the phenotypes became less pronounced (Figures S15A–B): cells from both WT and *PLXNB2* KO organoids displayed abundant expression of SOX2 and PAX6; and KO cells expressed slightly more β-III tubulin but less DCX than WT, reflecting premature neuronal differentiation as seen in 2D cultures (see Figures 5G-H and S8F-G). Dissociated cells from both WT and *PLXNB2* KO organoids labeled with fluorescent calcium indicator (Fluo-3 AM) or voltage sensitive probe (FluoVolt) displayed calcium activity and neuronal activity with caffeine stimulation (Figure S15C and Movies S9-10). Hence, *PLXNB2* KO or OE does not completely blocks neuronal differentiation, but profoundly affects 3D cytoarchitecture of the developing neuroepithelium (Figure 6G).

### Plexin-B2 integrates mechanochemical interaction through small GTPases and requires its extracellular domain

Plexin-B2 contains an intracellular Ras-GAP domain that functions to deactivate small GTPases such as R-Ras, M-Ras, and Rap1 ^54-56^, and a Rho-binding domain (RBD) that can bind to Rac1 or Rnd to regulate Plexin activity ^54,57^. The C-terminus of Plexin-B2 consists of a PDZ binding motif (amino acids VTDL) that provides a docking site for two Rho-GEFs (guanine nucleotide exchange factors), PDZ-Rho-GEF and LARG, to activate RhoA ^58^. Thus, upon binding to semaphorin, Plexin-B2 signals through either Ras or Rho small GTPases ^59^.

To examine which domains of Plexin-B2 are engaged in mechanoregulation, we carried out structure-function analysis by re-introducing into *PLXNB2* KO hESCs CRISPR-resistant *PLXNB2* signaling mutations in distinct domains: *PLXNB2*-mGAP (RR to AG mutations of critical arginines in the Ras-GAP domain), -mRBD (LSK to GGA mutation in the RBD domain), -ΔVTDL (deletion of PDZ binding motif), and -ΔECTO (deletion of extracellular domain) (Figures 7A-B). WB and ICC confirmed comparable expression levels of rescue Plexin-B2 signaling mutants in hESC lines (Figures S16A-B).

**Figure 7.**
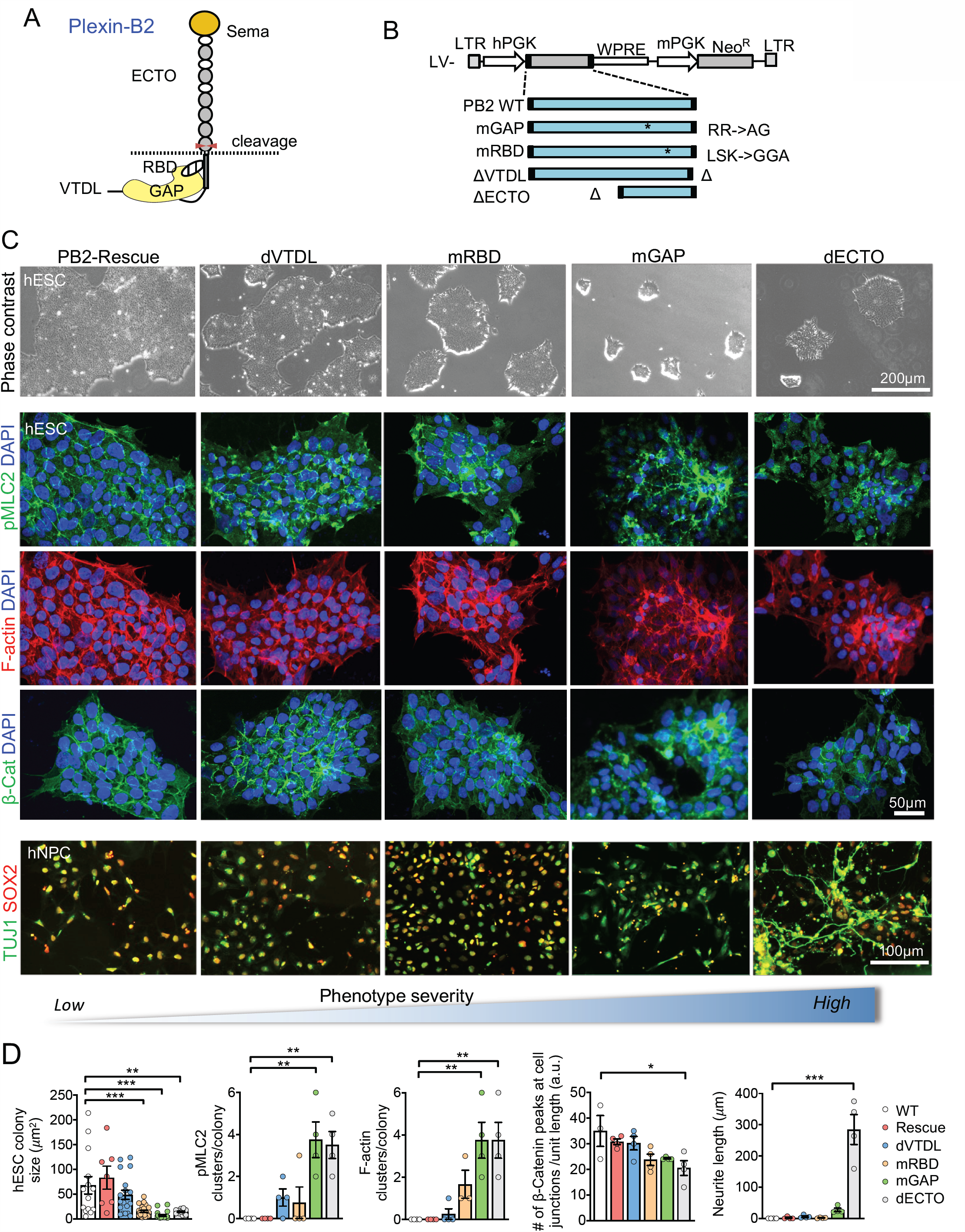
Structure-function analysis of Plexin-B2 signaling domains in mechanochemical integration. (A) Plexin-B2 domain structure. Sema, Sema domain; ECTO, ectodomain; RBD, Rho binding domain; GAP, Ras-GTPase domain; VTDL, four amino acids of the PDZ binding motif at C-terminus. Cleavage site for mature Plexin-B2 protein complex is denoted. (B) Diagram of lentiviral vectors expressing CRISPR-resistant wild type and signaling mutants of Plexin-B2. mGAP, mutation in GAP domain; mRBD, mutation in RBD domain; ΔVTDL, deletion of VTDL; ΔECTO, deletion of ECTO domain. (C) Phase contrast images (top panel row), ICC images for cytoskeletal and neuronal markers (middle panels), and summary diagram of phenotype severity (bottom) of rescue experiments with expression of CRISPR-resistant wild type or signaling mutants of Plexin-B2 in *PLXNB2* KO hESCs or hNPCs. (D) Graphs show hESC colony size, number of F-actin or pMLC2 aggregate foci per colony, distribution of β-catenin at intercellular junctions, and neurite length by β-III tubulin (TUJ1) from the rescue experiments. Data represent mean ± SEM. One-way ANOVA followed by Tukey’s multiple comparisons test versus WT. For hESC colony size: *F*_5, 90_= 10.55, n=4-8 fields. For pMLC2: *F*_5, 18_= 9.2, n= 4 fields; for F-actin: *F*_5, 17_= 10, n= 3-4 fields; for β-catenin: *F*_5, 17_= 3.736, n= 3-4 fields; for neurite length: *F*_5, 18_= 33.29, n= 3-4 fields. * p<0.05; * * p<0.01; * * * p<0.001. Also see Figure S16.

In hESCs, hNPCs and cerebral organoids, introducing CRISPR-resistant wild type *PLXNB2* rescued the with KO phenotypes, including growth kinetics, colony geometry, cell shape, actomyosin network, and junctional localization of β-catenin in hESC colony, neuronal differentiation in hNPC, and ventricular structure in organoids (Figures 7C-D and S16C). These phenotypes were nearly fully rescued with *PLXNB2*-ΔVTDL, partially rescued with -mRBD or -mGAP, but minimally rescued with -ΔECTO mutant. Thus the extracellular and Ras-GAP domains, less so the RBD domain, but not the C-terminal PDZ binding motif, are required for full signaling activity of Plexin-B2 for mechanoregulation.

We conducted additional pharmacological studies on the role of small GTPases in Plexin-B2-mediated mechanoregulation. The Rho inhibitor C3 transferase caused no overt changes in colony expansion of either WT or *PLXNB2* KO hESCs, in agreement with the dispensable nature of C-terminal PDZ binding motif for mechanoregulation by Plexin-B2. In contrast, inhibition of Rap1 (GGTI-298) or Rac1 (NSC-23766) both resulted in compromised colony expansion for WT hESCs, but no additive effect for the already reduced growth kinetics of *PLXNB2* KO hESCs (Figure S16D). Together, these data support engagement of Rap1 and Rac1, but not of Rho small GTPase, for Plexin-B2-mediated mechanoregulation.

## DISCUSSION

Multicellular organization relies on force-mediated processes that involve organization of actomyosin contractile network across cells and adjustment of junctional forces joining cells and cell-matrix in order to keep individual cells in a certain shape while maintaining tissue tension and cohesion ^60^. How cells sense, transmit, and regulate biochemical cues into force-generating processes is little understood. Here, combining a multitude of data from hESC, NPC, and cerebral organoid, spanning from cell morphology, colony geometry, actomyosin network, and junctional complexes, to AFM for measuring cell stiffness and surface topology, FRET tension sensor, and 3D cytoarchitecture, we uncovered a basic function of Plexin-B2 in mediating cell stiffness and tissue mechanics during multicellular organization. This basic function is consistent with plexin’s early evolutionary appearance during the transition from unicellular to multicellular organisms ^24^.

In contrast to studies on individual cells, our data indicate that Plexin-B2 mediates the mechanical state of a cohort of cells by adjusting their cytoskeletal status when interacting with one another and with matrix. Individual hESCs with Plexin-B2 KO or OE in low-density cultures behaved similarly as WT cells, but when cells started to form epithelium-like colonies or 3D aggregates, aberrations in cell morphology, tissue geometry and growth kinetics became evident. The phenotypes become more severe in structurally complex tissues such as neuroepithelium and VZ in cerebral organoids.

From a mechanical perspective, Plexin-B2 orchestrates all three key mechanoelements, i.e. actomyosin contraction, cadherin-mediated intercellular junctions, and integrin β1-based matrix adhesion, in a mechanochemical feedback loop ^33^. Earlier cell mechanics studies demonstrated the interconnectedness of these mechanoelements ^30,61,62^. Consistently, perturbation of Plexin-B2 in hESC and hNPC affected not only actomyosin contractility and cell stiffness, but also junctional recruitment of E-cadherin and ZO-1, surface stabilization of integrin β1, FA assembly, and FN deposition in ECM. Our results also echo the observation that inhibition of Rho, ROCK, myosin, myosin light chain kinase (MLCK), or F-actin polymerization itself, all can convert hESCs into a state of low tensile force, leading to alteration of actin cytoskeleton and stem cell functions ^63-65^. One interesting question is which mechanoelement in the feedback loop plays a more dominant role in dictating cell stiffness as regulated by Plexin-B2. The correlation between the experimental AFM data and mathematical simulation suggests that Plexin-B2 controls cell stiffness primarily through regulating actomyosin contractility, which in turn leads to adjustment of adhesive properties in order to maintain tissue integrity.

From a development point of view, our results support the model that mechanical forces and tissue architecture provide overarching signals to inform cell fate decisions ^66^. Plexin-B2 perturbation affects cell proliferation, differentiation, and migration of hESC and hNPC, which is linked to YAP mechanosensing activity and β-catenin signaling. Strikingly, reduced cell stiffness in hESC from Plexin-B2 deficiency promotes neuronal but not cardiomyoctye differentiation, akin to the impact of matrix stiffness on cell fate choice for hMSC ^50^. As induced neurons from Plexin-B2-deficient hESCs displayed comparable calcium activity and neuronal activation as WT controls, our study may point to a novel way to accelerate neuronal differentiation by manipulating cell mechanics.

In cerebral organoid, perturbation of Plexin-B2 signaling by gene ablation, Dox-inducible shRNA KD, OE, or replacement with signaling mutants all resulted in disrupted ventricular structures, a phenotype reminiscent of the neural tube closure defects seen in Plexin-B2 KO mice ^9^. It also echoes earlier studies that neural tube closure defects arise from deregulated mechanical interactions and tissue tension, which in turn impact cellular alignment and NPC function ^7^. Structural data indicated that class 4 semaphorins form dimers, and bring plexins on a neighboring cell into a dimeric configuration to initiate signaling ^67,68^. We found broad expression SEMA4B and 4C in hESCs and in the developing brain or cerebral organoids. Thus, a cohort of cells in a developing tissue may rely on semaphorin-plexin signaling to provide and receive spatial information for mechanochemical integration. Together, our data provide a force-based mechanism that unifies the multifaceted roles of Plexin-B2 in diverse tissues during development and repair.

From a signaling perspective, the extracellular, Ras-GAP, and, to a lesser extent, RBD domain of Plexin-B2 are essential for mechanoregulation, consistent with their evolutionary conservation ^24^. The main downstream effectors could be Rap1/2 families, which are pleiotropic regulators of cell-cell adhesion and cytoskeletal dynamics ^28^. By contrast, the PDZ-RhoGEF binding domain of Plexin-B2 does not appear to be required for mechanoregulation; in this context, our pharmacological studies confirmed the importance of the Rap1 and Rac1, but not RhoA, for hESC colony expansion. Our epistatic analyses further illustrate the link between Plexin-B2 and YAP: YAP/TAZ responded to Plexin-B2-mediated cell intrinsic stiffness by cytoplasm-nuclear shuttling, which accounted for proliferative state; moreover, YAP1-5SA rescued Plexin-B2 KO phenotypes, whereas YAP KD attenuated Plexin-B2 OE phenotypes. Hence, YAP functions not only as a mechanosensor ^46,47^, capable of sensing both external physical constraints and internal tensile forces, but also as a mechanoregulator downstream of Plexin-B2. In this context, it is noteworthy that YAP mutation resulted in flattening of body in medaka and zebrafish, indicating its requirement for maintaining tissue tension ^40^. Intriguingly, Rap2, a Ras-related GTPase, has been established as a key intracellular signal transducer that relays ECM rigidity signals to mechanosensitive cellular actives through YAP/TAZ ^6^. As Rap2 may also be a direct target of the Ras-GAP domain of Plexin-B2, it raises the tantalizing possibility that Rap may relay mechanical information from Plexin-B2 to YAP.

In summary, our study solidifies a mechanochemical integrative role of Plexin-B2 during multicellular organization. Our data also deepen the appreciation of force-mediated regulatory processes of cell fate specification and tissue morphogenesis, and may have broad implications in understanding neurodevelopmental disorders, tissue repair and regeneration.

## Acknowledgements

We thank all members of the Zou and Friedel laboratories for constructive comments. The work was supported by NINDS awards to H.Z. (NS073596) and R.H.F (NS092735), and NY state DOH award to H.Z. (C32242GG); NIDDK R01s DK106035 to G.L.G and DK118222 to E.U.A.; NIH/NIDCR T32HD075735 to R.J.W. CAPES and CNPq funding to R.A.D and J.P.M. Additional fellowship support was provided by National Council for Scientific and Technological Development (CNPq, Brazil; #200358/2015-4) for C.J.A, and the São Paulo Research Foundation (FAPESP, Brazil; 2016/07541-5) for R.D.

## Author Contributions

C.J.A., R.H.F., and H.Z. designed the study, conducted data analysis and wrote the manuscript. C.J.A., R.D., T.H., and R.T. conducted experiments. N.M.T. provided human fetal brain tissues and expertise in neuropathology analysis. R.W. performed AFM measurements and surface topology study, G.L.G. and E.U.A. assisted with the design, execution and analysis of the AFM and mechanobiological studies. N.D. established the cerebral organoids protocol. R.A.D. and J.P.M. performed mathematical modeling. All authors discussed drafts for preparation of final manuscript.

## Declaration of Interests

All authors declare no competing interests.

## References

1. Mammoto, T. & Ingber, D.E. Mechanical control of tissue and organ development. Development 137, 1407–1420 (2010).

2. DuFort, C.C., Paszek, M.J. & Weaver, V.M. Balancing forces: architectural control of mechanotransduction. Nat Rev Mol Cell Biol 12, 308–319 (2011).

3. Pilot, F. & Lecuit, T. Compartmentalized morphogenesis in epithelia: from cell to tissue shape. Dev Dyn 232, 685–694 (2005).

4. Fernandez-Gonzalez, R., Simoes, S.e.M., Röper, J.C., Eaton, S. & Zallen, J.A. Myosin II dynamics are regulated by tension in intercalating cells. Dev Cell 17, 736–743 (2009).

5. Rosowski, K.A., Mertz, A.F., Norcross, S., Dufresne, E.R. & Horsley, V. Edges of human embryonic stem cell colonies display distinct mechanical properties and differentiation potential. Sci Rep 5, 14218 (2015).

6. Meng, Z., et al. RAP2 mediates mechanoresponses of the Hippo pathway. Nature 560, 655–660 (2018).

7. Nikolopoulou, E., Galea, G.L., Rolo, A., Greene, N.D. & Copp, A.J. Neural tube closure: cellular, molecular and biomechanical mechanisms. Development 144, 552–566 (2017).

8. Tamagnone, L., et al. Plexins are a large family of receptors for transmembrane, secreted, and GPI-anchored semaphorins in vertebrates. Cell 99, 71–80 (1999).

9. Friedel, R.H., et al. Plexin-B2 controls the development of cerebellar granule cells. J Neurosci 27, 3921–3932 (2007).

10. Deng, S., et al. Plexin-B2, but not Plexin–B1, critically modulates neuronal migration and patterning of the developing nervous system in vivo. J Neurosci 27, 6333–6347 (2007).

11. Wansleeben, C., et al. An ENU-mutagenesis screen in the mouse: identification of novel developmental gene functions. PLoS One 6, e19357 (2011).

12. Hirschberg, A., et al. Gene deletion mutants reveal a role for semaphorin receptors of the plexin-B family in mechanisms underlying corticogenesis. Mol Cell Biol 30, 764–780 (2010).

13. Saha, B., Ypsilanti, A.R., Boutin, C., Cremer, H. & Chedotal, A. Plexin-B2 regulates the proliferation and migration of neuroblasts in the postnatal and adult subventricular zone. J Neurosci 32, 16892–16905 (2012).

14. Gurrapu, S. & Tamagnone, L. Transmembrane semaphorins: Multimodal signaling cues in development and cancer. Cell Adh Migr 10, 675–691 (2016).

15. Koropouli, E. & Kolodkin, A.L. Semaphorins and the dynamic regulation of synapse assembly, refinement, and function. Curr Opin Neurobiol 27, 1–7 (2014).

16. Kumanogoh, A. Semaphorins - A Diversity of Emerging Physiological and Pathological Activities, (Springer Japan, Tokyo, 2015).

17. Kumanogoh, A. & Kikutani, H. Immunological functions of the neuropilins and plexins as receptors for semaphorins. Nat Rev Immunol 13, 802–814 (2013).

18. Negishi-Koga, T. & Takayanagi, H. Bone cell communication factors and Semaphorins. Bonekey Rep 1, 183 (2012).

19. Pasterkamp, R.J. Getting neural circuits into shape with semaphorins. Nat Rev Neurosci 13, 605–618 (2012).

20. Sakurai, A., Doçi, C.L., Doci, C. & Gutkind, J.S. Semaphorin signaling in angiogenesis, lymphangiogenesis and cancer. Cell Res 22, 23–32 (2012).

21. Tran, T.S., Kolodkin, A.L. & Bharadwaj, R. Semaphorin regulation of cellular morphology. Annu Rev Cell Dev Biol 23, 263–292 (2007).

22. Worzfeld, T. & Offermanns, S. Semaphorins and plexins as therapeutic targets. Nat Rev Drug Discov 13, 603–621 (2014).

23. Xia, J., et al. Semaphorin-Plexin Signaling Controls Mitotic Spindle Orientation during Epithelial Morphogenesis and Repair. Dev Cell 33, 299–313 (2015).

24. Junqueira Alves, C., Yotoko, K., Zou, H. & Friedel, R.H. Origin and evolution of plexins, semaphorins, and Met receptor tyrosine kinases. Sci Rep 9, 1970 (2019).

25. Kong, Y., et al. Structural Basis for Plexin Activation and Regulation. Neuron 91, 548–560 (2016).

26. Suzuki, K., et al. Structure of the Plexin Ectodomain Bound by Semaphorin-Mimicking Antibodies. PLoS One 11, e0156719 (2016).

27. Pascoe, H.G., Wang, Y. & Zhang, X. Structural mechanisms of plexin signaling. Prog Biophys Mol Biol 118, 161–168 (2015).

28. Kooistra, M.R., Dubé, N. & Bos, J.L. Rap1: a key regulator in cell-cell junction formation. J Cell Sci 120, 17–22 (2007).

29. Bos, J.L. From Ras to Rap and Back, a Journey of 35 Years. Cold Spring Harb Perspect Med 8 (2018).

30. Li, D., et al. Integrated biochemical and mechanical signals regulate multifaceted human embryonic stem cell functions. J Cell Biol 191, 631–644 (2010).

31. Krtolica, A., et al. Disruption of apical-basal polarity of human embryonic stem cells enhances hematoendothelial differentiation. Stem Cells 25, 2215–2223 (2007).

32. Totsukawa, G., et al. Distinct roles of ROCK (Rho-kinase) and MLCK in spatial regulation of MLC phosphorylation for assembly of stress fibers and focal adhesions in 3T3 fibroblasts. J Cell Biol 150, 797–806 (2000).

33. Hannezo, E. & Heisenberg, C.P. Mechanochemical Feedback Loops in Development and Disease. Cell 178, 12–25 (2019).

34. Meltzer, S., et al. Epidermis-Derived Semaphorin Promotes Dendrite Self-Avoidance by Regulating Dendrite-Substrate Adhesion in Drosophila Sensory Neurons. Neuron 89, 741–755 (2016).

35. Azeloglu, E.U. & Costa, K.D. Atomic force microscopy in mechanobiology: measuring microelastic heterogeneity of living cells. Methods Mol Biol 736, 303–329 (2011).

36. Riedl, J., et al. Lifeact: a versatile marker to visualize F-actin. Nat Methods 5, 605–607 (2008).

37. Lecuit, T. & Lenne, P.F. Cell surface mechanics and the control of cell shape, tissue patterns and morphogenesis. Nat Rev Mol Cell Biol 8, 633–644 (2007).

38. Bachmann, M. Thermodynamics and Statistical Mechanics of Macromolecular Systems. (Cambridge University Press, 2014).

39. Binder, K. Monte Carlo and Molecular Dynamics Simulations in Polymer Science. (Oxford University Press, 1995).

40. Porazinski, S., et al. YAP is essential for tissue tension to ensure vertebrate 3D body shape. Nature 521, 217–221 (2015).

41. Singh, P., Carraher, C. & Schwarzbauer, J.E. Assembly of fibronectin extracellular matrix. Annu Rev Cell Dev Biol 26, 397–419 (2010).

42. Rolo, A., Skoglund, P. & Keller, R. Morphogenetic movements driving neural tube closure in Xenopus require myosin IIB. Dev Biol 327, 327–338 (2009).

43. Foty, R. A simple hanging drop cell culture protocol for generation of 3D spheroids. J Vis Exp (2011).

44. Cowin, P. & Burke, B. Cytoskeleton-membrane interactions. Curr Opin Cell Biol 8, 56–65 (1996).

45. Dupont, S., et al. Role of YAP/TAZ in mechanotransduction. Nature 474, 179–183 (2011).

46. Sudol, M., et al. Characterization of the mammalian YAP (Yes-associated protein) gene and its role in defining a novel protein module, the WW domain. J Biol Chem 270, 14733–14741 (1995).

47. Zhao, B., Lei, Q.Y. & Guan, K.L. The Hippo-YAP pathway: new connections between regulation of organ size and cancer. Curr Opin Cell Biol 20, 638–646 (2008).

48. Sun, Y., et al. Hippo/YAP-mediated rigidity-dependent motor neuron differentiation of human pluripotent stem cells. Nat Mater 13, 599–604 (2014).

49. Grashoff, C., et al. Measuring mechanical tension across vinculin reveals regulation of focal adhesion dynamics. Nature 466, 263–266 (2010).

50. Halder, G., Dupont, S. & Piccolo, S. Transduction of mechanical and cytoskeletal cues by YAP and TAZ. Nat Rev Mol Cell Biol 13, 591–600 (2012).

51. Lancaster, M.A., et al. Cerebral organoids model human brain development and microcephaly. Nature 501, 373–379 (2013).

52. Lancaster, M. & Knoblich, J. Generation of cerebral organoids from human pluripotent stem cells. Nat Protoc 9, 2329–2340 (2014).

53. Daviaud, N., Chen, K., Huang, Y., Friedel, R.H. & Zou, H. Impaired cortical neurogenesis in plexin-B1 and -B2 double deletion mutant. Dev Neurobiol 76, 882–899 (2016).

54. Oinuma, I., Ishikawa, Y., Katoh, H. & Negishi, M. The Semaphorin 4D receptor Plexin-B1 is a GTPase activating protein for R-Ras. Science 305, 862–865 (2004).

55. Saito, Y., Oinuma, I., Fujimoto, S. & Negishi, M. Plexin-B1 is a GTPase activating protein for M-Ras, remodelling dendrite morphology. EMBO Rep 10, 614–621 (2009).

56. Wang, Y., et al. Plexins are GTPase-activating proteins for Rap and are activated by induced dimerization. Sci Signal 5, ra6 (2012).

57. Vikis, H.G., Li, W. & Guan, K.L. The plexin-B1/Rac interaction inhibits PAK activation and enhances Sema4D ligand binding. Genes Dev 16, 836–845 (2002).

58. Swiercz, J.M., Kuner, R. & Offermanns, S. Plexin-B1/RhoGEF-mediated RhoA activation involves the receptor tyrosine kinase ErbB-2. J Cell Biol 165, 869–880 (2004).

59. Zhou, Y., Gunput, R.A. & Pasterkamp, R.J. Semaphorin signaling: progress made and promises ahead. Trends Biochem Sci 33, 161–170 (2008).

60. Sasai, Y. Cytosystems dynamics in self-organization of tissue architecture. Nature 493, 318–326 (2013).

61. Parsons, J.T., Horwitz, A.R. & Schwartz, M.A. Cell adhesion: integrating cytoskeletal dynamics and cellular tension. Nat Rev Mol Cell Biol 11, 633–643 (2010).

62. Daley, W.P., Peters, S.B. & Larsen, M. Extracellular matrix dynamics in development and regenerative medicine. J Cell Sci 121, 255–264 (2008).

63. Watanabe, K., et al. A ROCK inhibitor permits survival of dissociated human embryonic stem cells. Nat Biotechnol 25, 681–686 (2007).

64. Walker, A., et al. Non-muscle myosin II regulates survival threshold of pluripotent stem cells. Nat Commun 1, 71 (2010).

65. Chen, G., Hou, Z., Gulbranson, D.R. & Thomson, J.A. Actin-myosin contractility is responsible for the reduced viability of dissociated human embryonic stem cells. Cell Stem Cell 7, 240–248 (2010).

66. Fletcher, D.A. & Mullins, R.D. Cell mechanics and the cytoskeleton. Nature 463, 485–492 (2010).

67. Janssen, B.J., et al. Structural basis of semaphorin-plexin signalling. Nature 467, 1118–1122 (2010).

68. Wang, Y., Pascoe, H.G., Brautigam, C.A., He, H. & Zhang, X. Structural basis for activation and non-canonical catalysis of the Rap GTPase activating protein domain of plexin. Elife 2, e01279 (2013).

