## Supplementary Material and Figures for "Plexin-B2 is a key regulator of cell mechanics during multicellular organization"

**Supplemental Information**

1. **EXPERIMENTAL MODEL AND SUBJECT DETAILS**
2. **METHOD DETAILS**
3. **SUPPLEMENTAL TABLE LEGENDS**
4. **SUPPLEMENTAL MOVIE LEGENDS**
5. **SUPPLEMENTAL REFERENCES**

#### EXPERIMENTAL MODEL AND SUBJECT DETAILS

##### hESC culture

All culture experiments were conducted using WA09 (H9) hESC [1] obtained from University of Wisconsin and validated at Mount Sinai Stem Cell core facility. The Mount Sinai Stem Cell core conducts routine quality controls by verifying karyotype and mycoplasma-free status for all stem cell lines. Briefly, H9 hESC were seeded on matrigel-coated cell culture dishes (1:100, Corning) and grown in mTeSR1 medium (STEMCELL Technologies). Cells were passaged around every five days by dissociation with Accutase (BD Biosciences) and addition of 2.5 μM Thiazovivin (Millipore). The study was approved by the Embryonic Stem Cell Research Oversight Committee (ESCRO) at Icahn School of Medicine at Mount Sinai.

##### Human fetal brain tissues

Human fetal brain tissues were obtained from the Department of Pathology, Icahn School of Medicine at Mount Sinai. All human tissue samples were de-identified and collected in accordance with the policies and regulations of Institutional Review Board, Icahn School of Medicine (IRB#AAAJ9652-Y1M00, HS#14-01007) using appropriate autopsy research consent. Fetal brains were collected from postmortem specimens ranging from 18 to 23 week post-conception, fixed in formalin and embedded in paraffin. Serial histological sections (4 μm thick) of cerebral cortex and germinal matrix were dried at 70°C for 1 hr, deparaffinized in two changes of xylene followed by four serial dilutions of ethanol, and rehydrated in $dH_2O$. Antigen retrieval was performed by antigen unmasking solution (pH 6.0) (Vector Labs) and heating in microwave for 2 min. Slides were washed in $dH_2O$ for 5 min and blocked in 5% Donkey Serum/0.5% Triton in PBS for 30 min before adding the primary antibody.

#### METHOD DETAILS

##### CRISPR genome editing for Plexin-B2 knockout

The CRISPR/Cas9-sgRNA sequence targeting *PLXNB2* exon 3 was embedded inside a CRISPR/Cas9 lentivirus plasmid [2,3] to generate plentiCRISPRv2-sgPLXNB2 (target sequence: GTTCTCGGCGGCGACCGTCA). Lentiviral particles were produced in HEK293T cells by co-transfection of lentiviral plasmid, envelope plasmid pMD2.G, and packaging plasmid psPAX2 (Addgene plasmids #12259 and #12260; deposited by Didier Trono, EPFL Lausanne). Low passage hESCs transduced with the lentivirus were selected with puromycin (1 μg/ml). Successful knockout of Plexin-B2 was confirmed by Western blots, demonstrating absence of mature Plexin-B2 (170 kDa). A lentiviral vector (plentiCRISPRv2-sgEGFP) expressing sgRNA targeting eGFP (target sequence: GGGCGAGGAGCTGTTCACCG) was used as control (wild-type, WT). Plasmids have been deposited at Addgene.org.

##### CRISPR clones sequencing and karyotyping

To isolate clonal lines of hESCs that carry defined CRISPR mutations, cells were transduced with lentiviruses (either targeting Plexin-B2 or EGFP as control) and selected with puromycin (1 μg/ml) for 1 wk. For each condition, three colonies were picked and sequenced for verify mutations at *PLXNB2* locus. Three KO and three WT clones were confirmed. Normal karyotypes of all clonal lines were confirmed by chromosome G-banding at Mount Sinai Genetic Testing Laboratory Sema4 (New York, USA).

##### Lentiviral vectors for Plexin-B2 rescue, overexpression and signaling mutants

For rescue experiments, human *PLXNB2* cDNA was modified to generate a CRISPR-resistant *PLXNB2* cDNA, which was inserted into the lentiviral vector pLenti-PGK (plasmid backbone pLENTI PGK Neo DEST, Addgene #19067; deposited by Eric Campeau & Paul Kaufman) by Gateway LR Clonase II (Thermo Fischer Scientific). The resulting vector pLV-*PLXNB2* was then used to generate lentiviruses to transduce *PLXNB2* KO hESC for rescue experiments. The pLV-*PLXNB2* lentivirus was also used for overexpression experiments by transducing wild type hESC.

The four functional domains of Plexin-B2 containing mutations or deletions were generated in *PLXNB2* cDNA (CRISPR resistant) using the Phusion Site-Directed Mutagenesis Kit (Thermo-Fischer): mGAP (mutation in the GAP domain: R1391A, R1392G), mRBD (mutation in the RBD domain: LSK 1558-1560 -> GGA), ∆VTDL (deletion of four last C-terminal amino acids VTDL of PDZ binding motif) and ∆ECTO (lacking extracellular domain, deletion of amino acids 30-1189). The mutant cDNAs were cloned into pLenti-PGK destination vector as described above. Plasmids have been deposited at Addgene.org.

**Doxycycline-inducible shRNA Plexin-B2 knockdown**

Temporally controlled knockdown of Plexin-B2 at different phases of hESC colony expansion and cerebral organoid maturation was achieved with TET-ON lentiviral vectors (Tet-pLKO-Puro backbone, Addgene #21915, deposited by Dimitri Wiederschain) expressing doxycycline-inducible shRNA targeting *PLXNB2*, along with puromycin resistance marker (pLKO-Tet-On-*PLXNB2*-shRNA1 and –shRNA2). A non-targeting shRNA vector (pLKO-Tet-On-shRNA-Control) served as control. Stable TET-ON cell lines were established by puromycin (1µg/ml) selection. The shRNA-mediated *PLXNB2* knockdown was induced by adding 1µg/ml doxycycline (MP Biomedicals) to culture medium. Plasmids have been deposited at Addgene.org.

**Doxycycline-inducible YAP overexpression and short hairpin constructs**

DNA plasmids for lentiviral vectors with doxycycline-inducible human *YAP1* and *YAP1-5SA* cDNAs were commercially manufactured by VectorBuilder. Plasmids were designed using VectorBuilder website tools. In brief, human *YAP1* plasmid was designed using an mRNA sequence from NCBI transcript NM_001282101, and human *YAP1-5SA* was designed based on [4] to generate a unphosphorylated constitutively active form of *YAP1*. Both WT and mutant *YAP1* plasmids are driven by a TRE3G promoter and have a CMV-puromycin-T2A-mCherry cassette inserted after the gene of interest to facilitate selection of transfected cells. A second plasmid to express TeT3G protein was designed to include a PGK-hygromycin cassette to facilitate cell selection.

YAP short hairpin (sh)RNA lentiviral expression vectors were cloned by ligating oligonucleotides encoding the shRNA hairpins (Integrated DNA Technologies) and into the pLKO-Tet-On vector (Addgene #21915) to have temporal control of YAP1 knockdown. All plasmids were validated using restriction enzymes and Sanger sequencing (Macrogen, New Jersey) using a primer (ggcagggatattcaccattatcgtttcaga) that annealed upstream of the H1/TO promoter.

**hESC colony and 3D aggregation assays**

For adherent hESC culture, cells were plated at a density of $5 \times 10^4$ in Matrigel coated 6-well plates, and individual colonies were imaged at day 2 and 6 after passage and sizes were measured using ImageJ. Spontaneous differentiation in hESC colonies was assessed by alkaline phosphatase live stain (Thermo). Images were captured using Zeiss microscopes (AxioCam) and mean fluorescence intensity was quantified by ImageJ.

For 3D aggregation assay was performed by culturing $4 \times 10^4$ hESCs resuspended in mTeSR1 medium in suspension in low-adhesion 24-well plates with addition of 10 µM Thiazovivin. The size of hESC aggregates was evaluated after 48 hours. For hanging drop assay, $2.5 \times 10^4$ hESCs were resuspended in 10 µl of mTeSR1 medium with 10 µM Thiazovivin for 24 hours. Images were taken using Olympus CKX53 inverted microscope, and the areas of the compacted cells in the center were measured using ImageJ.

**Cerebral organoid derivation from hESCs**

Cerebral organoids were generated as described per [5,6] with slight modifications. Briefly, H9 hESCs (< 40 passages) were dissociated into single cells with Accutase (BD Biosciences) and plated in ultra-low attachment 96-U-well plates (Costar) at a density of 9,000 cells/well (150 µl per well) in mTeSR1 medium (STEMCELL Technologies) containing 10 μM Rho-associated protein kinase (ROCK) inhibitor Thiazovivin (Millipore). After 6 days, embryoid bodies were treated with neural induction medium STEMDiff (STEMCELL Technologies). The neuroepithelial tissue that formed over the next 4-6 days was transferred into 30 μl matrigel droplets (Corning) and grown for 4 days in stationary condition in culture medium composed of DMEM-F12 (Invitrogen), Neurobasal medium (Invitrogen), N2 supplement (Invitrogen), human insulin (Sigma), GlutaMAX supplement (Gibco), MEM-NEAA (Gibco), penicillin-streptomycin (Sigma), 2-mercaptoethanol (Millipore), and B27 supplement minus vitamin A (Invitrogen). This was then followed by culturing in 6-well plates on an orbital shaker with addition of B27 supplement plus vitamin A (Inivitrogen) to the media (rotationary medium) to promote neuronal differentiation. At day 42 after initiation of the protocol, cerebral organoids developed to a size of ~3-5 mm in diameter and were analyzed.

**Neural differentiation**

NPCs were generated using either single-cell plating monolayer cultures or embryoid body protocols with STEMDiff medium (Stem Cell Technologies). For the monolayer culture protocol, hESC were plated at high density (2.5 x $10^5$ cells/$cm^2$) in STEMDiff medium, and at day 6, the cells were re-plated at high confluence, and at day 14, cells were plated on glass coverslips coated with Matrigel and analyzed by immunocytochemistry. For the embryoid body protocol, hESC (9x$10^3$) were cultured in suspension for 4 days in mTeSR1 medium in the presence of 10 μM Thiazovivin. The EBs were plated onto Matrigel-coated glass coverslips and maintained in STEMDiff medium until day 14 for immunocytochemistry.

**Cardiac differentiation**

The cardiac differentiation of hESCs was performed using the PSC Cardiomyocyte Differentiation Kit (Thermo). Briefly, after 3 consecutive passages using enzymatic dissociation (single-cell condition) with Tryple (Thermo) reagent, hESCs were seeded in Matrigel-coated 6-well plate (50-60% confluence) in mTSeR1 media supplemented with 2.5 μM Thiazovivin. When hESCs reached 75-90%

confluence (48 or 72 hours after plating), mTSeR1 media was replaced by Media A (mesoderm induction). Two days later (day 6) the media was replaced by Media B. After day 8 the cells were kept in cardiomyocyte maintenance media and at day 30 (approx. 20 days after the control cells started beating), cells were fixed, stained with phalloidin for F-actin and with anti-cardiac troponin T antibodies for immunocytochemistry. The shRNA *PLXNB2* knockdown was induced by adding 1 μg/ml doxycycline (MP Biomedicals) to culture medium.

**Histology and immunofluorescence staining**

For immunocytochemistry (ICC), cells were fixed in 4% formaldehyde in PBS at room temperature for 10 min, washed 3x with PBS, permeabilized and blocked with 5% donkey serum and 0.3% Triton X-100 in PBS for 1 hr. Primary and secondary antibody incubations as well as image acquisition were performed as described above.

For histological analysis, cerebral organoids were fixed in 4% paraformaldehyde/PBS at 4°C for 15 min, washed with PBS, cryoprotected in 30% sucrose at 4°C overnight, embedded in OCT (Fisher HealthCare), frozen on dry ice and stored at -80°C. The organoids were sectioned using cryostat (Leica) at 12 μm thickness and the cryosections were used for hematoxylin-eosin (H&E) histology and immunohistochemistry (IHC).

For IHC, organoid cryosections were permeabilized and blocked with 5% donkey serum and 0.3% Triton X-100 in PBS for 1 hr. Primary antibodies were incubated at 4°C overnight in PBS with 1% BSA and 0.3% Triton X-100. Secondary antibodies were incubated using the same buffer at room temperature for 1 hour. Primary antibody information is listed in Table S1. Secondary antibodies used were Alexa Fluor 488, 594, or 647-conjugated donkey anti-goat, -rabbit, -rat, or -mouse IgG and goat anti-guinea pig IgG (Jackson ImmunoResearch Laboratories, 1:300).

**pLifeAct-mScarlet and CellMask labeling**

Lentiviral preparations for introducing LifeAct-mScarlet that labels *in vivo* F-actin [7] were transduced into hESCs or hNPCs, selected with G418 (200$\mu$g/mL) and expanded for subsequent experiments. For CellMask green plasma membrane stain (Thermo Fisher Scientific), live hESCs were incubated with the dye for 5 min, and imaged immediately using Zeiss LSM 780 confocal microscope to videography of dynamic changes of plasma membranes and cellular borders.

The circularity index of cells was calculated with the formula: 4π*area/perimeter^2. A value of 1.0 indicates a perfect circle; a value approaching 0.0 indicates an increasingly elongated shape.

**Time lapse video microscopy**

hESCs were plated at low density, and live-cell videography was carried out from day 2 to 5, using Zeiss LSM 710 microscope, with environmental control chamber conditions set at 37°C temperature and 5% $CO_2$, every 120 min over 3 days. Quantification was performed using ImageJ, to measure colony size and contour of the colony geometry in each frame. Migration of colony was measured by tracking a single point in the middle of individual colony in each frame, averaged every 120 min over 3 days using MTrackJ [8]. For LifeAct-mScarlet movies, frames were captured every 20 min for 360 min duration and for CellMask movies, frames were taken every 3 min for 60 min duration.

**EdU Click-iT assay**

To assess the proliferative potential of hESCs, cells were plated at a density of 2 x $10^4$ in Matrigel-coated 8-chamber slides, and after 3 days of culture, the nucleotide analog 5-ethynyl-2'-deoxyuridine (EdU, Life Technologies) was added at 10 μM to the medium for 30 min, after which cells were washed with PBS and fixed in 4% PFA/PBS at room temperature for 10 min. EdU Click-iT assay was performed per manufacturer's instruction (Thermo), followed by DAPI staining. The number of $EdU^+$ cells was quantified using Photoshop CS5 (Adobe).

For EdU pulse-chase assay, cerebral organoids were pulsed with 10 μM EdU for 30 min, the media was then washed off and replenished, and after 24 hour chase, the organoids were fixed in 4% PFA/PBS. The EdU Click-iT assay was performed and the sections were prepared for subsequent immunofluorescence staining with Ki67 antibody. Quantification was carried out using Photoshop CS5 to determine the percentage of cells that underwent active cell division (S-phase) a day earlier ($EdU^{+}$) but were no longer proliferative ($Ki67^{-}$), i.e. the exit or “quit fraction”.

**Subcellular fractionation**

A subcellular protein fractionation kit (Cell Signaling) was used to separate cell lysates into cytoplasmic (Cyto), membrane (Mem) and nuclear (Nuc) fractions based on the use of detergents per manufacturer’s protocol. The fractions were mixed with 3x SDS loading buffer with DTT and analyzed by Western blotting.

**Western blot**

For Western blot, freshly collected samples were lysed in RIPA buffer (Sigma) containing protease and phosphatase inhibitors (Sigma). Protein concentrations were determined using BCA assay (Pierce Thermo Scientific). Proteins were resolved on 4-12% polyacrylamide NuPAGE gels (Invitrogen) using the XCell SureLock system (Invitrogen) and transferred to nitrocellulose membranes (Li-Cor Biosciences). The immunoreactive bands were detected by fluorescent ODYSSEY infrared imaging system (Li-Cor Biosciences). Equal protein loading was controlled by probing the blots with an antibody against β-actin. A complete list of primary antibodies is in Table S1. Infra-red dye conjugated secondary antibodies were: IRDye 800CW and 680RD-conjugated donkey anti-sheep, -rabbit, or -mouse (Li-Cor Biosciences, all used at 1:10,000 dilution).

**qRT-PCR array and validation**

Cells were lysed using RLT buffer (Qiagen) plus 2-mercaptoethanol (Millipore), and lysates were stored at -80°C until RNA extraction. Total RNA was isolated using the RNeasy Mini Kit (Qiagen).

RNA was converted to cDNA using oligo(dT) primer, and quantitative PCR reactions were set up using the PerfeCTa SYBR Green FastMix ROX qPCR kit (QuantaBio). Quantitative RT-PCR array analysis was performed using a hESC-specific PCR array (PAHS-081YA; SABiosciences). Reactions were run on an ABI/Life Technologies 7900HT real-time PCR instrument.

Validation of qRT-PCR array was performed with individual qRT-PCR reactions using the primers for the indicated genes (primers sequences are listed in Table S2). Relative mRNA expression levels were determined by ΔΔCt method relative to *GAPDH* housekeeping control. qRT-PCR data are average from at least 3 technical replicates for each sample.

**Dissociation of cerebral organoids, calcium imaging, and FluoVolt analysis**

For imaging of calcium sparks, cerebral organoids were collected in Cultrex organoid harvesting solution (AMSBIO), dissociated into single cells using Accutase, and plated on Matrigel-coated coverslips. Four days after plating, cells were loaded with 10 μM Fluo-3 AM (Biotium) for 30 min at room temperature, washed, and superfused with Tyrode's solution. Fluo-3 AM was excited at 488 nm, and emitted fluorescence above 505 nm was recorded by confocal microscopy (LSM 5 Exciter) in frame scan mode on a heated stage at 37°C. Cells were stimulated using a puff of 10 mM caffeine (Sigma).

For imaging of action potentials, the dissociated cells were loaded with FluoVolt (Thermo-Fisher). Briefly, cells plated in single-cell conditions on Matrigel-coated coverslips were washed 3 times with Tyrode's solution at room temperature. Cells were incubated in FluoVolt loading solution (Component A, B, and Tyrode's solution) at room temperature for 20 min, followed by 3 washes with Tyrode's solution. FluoVolt was excited at 488 nm, and emission above 505 nm was recorded by confocal microscopy (LSM 5 Exciter) as described above. Cells were stimulated using a puff of 10 mM caffeine (Sigma).

**Cytoskeletal drugs and integrin antibody blocking experiments**

Cytoskeletal drugs and integrin antibody blocking experiments were carried out directly in live hESCs transfected with LifeAct_mScarlet. Drugs or vehicles were added to media, and cells were incubated for 3 hr in a cell incubator. Drugs were used at the following concentrations: Rock Inhibitor Y-27632 (20μM), blebbistatin (10μM) and latrunculin (5μM). Integrin-β1 antibody (P5D2; sc-13590) 10 μg/mL or IgG antibody (used as control) were added to cells in suspension for 15 min before plating in Matrigel-coated coverslips, the mTeSR media was supplemented with 2.5 μM Thiazovivin (Millipore) and cells were incubated for 1 hr before imaging. After incubation hESCs were treated with CellMask dye (Thermo) for 5 min and imaged immediately using Zeiss LSM 780 confocal microscope.

**Pharmacological inhibition of small-GTPases**

hESCs ($10^4$ cells seeded in 24-well plate) were treated during 5 days of culture with small GTPase inhibitors for RhoA (C3, 2.5μg/ml; Cytoskeleton), Rap (GGTI, 1.5μM; Tocris Bioscience) or Rac1 (NSC, 5 μM; Tocris Bioscience). At day 6, hESC colonies were stained with cresyl violet, images were captured using a stereo microscope (Nikon SMZ-745T) and the area of hESC colonies was measured with ImageJ software.

**Atomic force microscopy elastography**

Atomic force microscope (AFM) measurements were conducted using an Asylum MFP 3D-BIO AFM equipped with environmental thermal and vibrational control coupled with an Olympus IX-80 inverted spinning disk confocal microscope as previously described [9]. Briefly, the AFM employs a specialized and finely controllable probe that is capable of coming into direct physical contact with live cells in fluid without injuring them. AFM uses its probe to indent directly into a cell and measures how much force was required, which in turn is used to calculate stiffness. For the experiments, hESCs were plated on 60 mm Matrigel coated plastic dishes at a density of 5 x $10^5$ for WT and *PLXNB2* OE groups and 7.5 x $10^5$ cells for *PLXNB2* KO group (to compensate for slower colony expansion). hNPCs were plated at a density of 4.2 x $10^6$ for WT and *PLXNB2* OE groups and 6.3 x $10^6$ cells for *PLXNB2* KO group on 60 mm dishes. After 3 day culture, the plates were secured to the AFM, and maintained at 37°C for the entire duration of the AFM experiment. A gold-coated silicon nitride probe with triangular shaped body, blunted pyramidal tip, and nominal spring constant of 0.1 N/m (Cat #: TR400PB, Asylum) was used for cell measurements and calibrated using the thermal noise method. The Hertzian model was used to fit our indentation data for its elastic modulus, where this model called for probe tip half angle (9°), probe material (silicon nitride), and sample Poisson's ratio (v = 0.45). To measure the colony elastic modulus a 24 x 24 indentation array was conducted over a 64 $\mu m^2$ region of the colony (avoiding colony edges). The AFM was set to conduct its indentation array at an indentation rate of 0.5 Hz and indent each time until the probe deflected 50 nm (about 6 nN of force) which would trigger its retraction. For hESCs, each condition was measured for 6-7 different colonies across 6-7 different dishes from two independent preparations. For hNPCs, each cell condition was measured for 3 colonies across 3 different dishes. For each cell group, every indentation was aggregated into a pool of measurements which was then averaged to return a mean colony stiffness. This aggregation allowed for proper sampling power regarding biological variability and batch variation. Due to the phenotypic nature of hNPCs, NPC cultures had open/dish exposed areas, which translated into direct dish measurements during AFM indentations, a phenomenon called substrate effect. To avoid these unwanted influences, NPC modulus values larger than 20 kPa were filtered out (expected NPC stiffness ~5 kPa). Mean colony stiffness trends and statistical comparisons were done on both filtered and unfiltered NPC data and confirmed to be equivalent. For statistical analysis a one-way ANOVA with Kruskall-Wallis post hoc correction for multiple pairwise comparisons was conducted in Prism 8 (GraphPad).

**Cell topology imaging via atomic force microscopy**

In addition to elasticity measurements, the atomic force microscope (AFM) can image colony topology via direct contact. We employed the AFM imagining technique called contact mode imaging whereby a feedback loop is set upon the probe as it is told to slide across, and track, the surface of the colony in a raster scanning manner. hESCs were prepared as above except that cells were fixed with formaldehyde for 15 min. This allowed preservation of colony structure and features throughout the duration of AFM imaging, as well as increase colony robustness during extended AFM probe interaction. This step was acceptable for contact mode imaging but would not be appropriate for elasticity measurements as fixing may changes the cells elastic properties. The same type of probe used for AFM elasticity measurements was selected for contact mode imaging (see methods above), its 0.1 N/m spring constant and 42 nm tip radius allowed for gentler contact imaging and detection of finer cell features. A 90 $\mu m^2$ region central to the cell colony was selected for imaging. The probe was prescribed to move at a speed of 200 μm/second and scan its 90 $\mu m^2$ region in 256 segments, these 256 segments were then stitched together for a complete topography map. Scans used a setpoint=0.8 volts. Topology images revealed ruffles across the cell membrane particularly for the *PLXNB2* KO group. To quantify these ruffles a profile was taken through a selected cell, perpendicular to the ruffle direction. That profile was then counted for cell membrane features (peaks) of amplitude greater than 15 nm of probe deflection. Each cell was measured for 4 profiles, resulting in a total (sum) of ruffles per cell. 5 cells were selected per image (2 images per cell type) and then averaged for a mean # of ruffles per cell.

**FRET analysis using vinculin tension biosensor**

Analysis of FRET was performed as described [10]. Human neuroprogenitor cells were transfected with vinculin tension biosensor using lentiviral particles as described above. Forty-eight hours after transduction, cells were sorted by FACS (BD FACSAria IIu). Media was then supplemented with CloneR (StemCell Technologies) and cells were expanded for the imaging experiment. Live-cell FRET imaging was performed using Zeiss LSM 880 confocal microscope equipped with 1.4 NA water objective and with humidity and $CO_2$ control. eCFP was excited with 458 nm emission line using an Argon ion laser and eYFP was excited with 515 nm emission line. eCFP, FRET and eYFP channels were simultaneously imaged using an internal GaAsp detector collecting the emission from a range of wavelengths appropriate for each channel: For eCFP channel: 463 nm – 520 nm, FRET channel:

520 nm – 620 nm and eYFP channel: 520 nm – 620 nm. Intensity based ratiometric FRET indices and heatmaps were obtained on a per-cell and per-focal adhesion basis using custom-written scripts in MATLAB and ImageJ plugin (FRET Analyzer). Stepwise, images were sorted into donor, FRET and acceptor channels. Each set of images from different channels was registered and background corrected. To obtain FA masks, CLAHE and Laplacian of Gaussian (LoG) were performed on the eYFP channel, followed by applying size and intensity filter on the resulting particles. Since vinculin tension biosensor is a single-chain construct, FRET Index was calculated by dividing the FRET channel (donor excitation with acceptor emission) by the CFP channel (donor excitation with donor emission).

**Quantification and statistical analysis**

Data are presented as mean ±SEM. Bar graphs with scatter dots were shown, unless otherwise indicated. Distributions of the raw data were tested for normality of distribution. One-way analysis of variance (ANOVA) analysis of variance and Tukey's or Dunnett's post-hoc test was performed for comparisons of data with more than two groups. Unpaired *t*-test (two sided) was applied when comparing data from two groups. For studies with repeated measures (RM), two-way ANOVA RM and Bonferroni post-hoc test was performed. Statistical analysis were performed using GraphPad Prism 8 statistical software. Statistical significance was considered as $p<0.05$ (*); $p<0.01$ (**); $p<0.001$ (***). Sample sizes and statistical details are indicated in figure legends.

**Mathematical modeling**

The computational model consists of simulating the mechanical rigidity of a cell membrane based on using AFM tip to measure force as a function of its deformation. Molecular Dynamics (MD) models with the approximation of macromolecular systems [11,12] were used to simulate the dynamics of a cell composed of membranes and actin filaments in two dimension (2-D).

**I. Cell Model:**

We developed a membrane model that can reproduce the main features of the mechanical structure of a cell. The cell membrane, nuclear membrane, and actin filaments are modeled as being formed by beads connected by springs. Specifically, the cell membrane is composed of 201 beads connected by springs forming a diameter of 5.25 μm. To describe the nuclear membrane, it was necessary to use 33 beads connected by springs, generating a diameter of 1.5 μm for the nucleus. Finally, the actin filaments, which are radial in the initial position, are separated into two types: type I filament has one end attached to the nuclear membrane and one end free, while type II filament has both ends free. For each type I configuration, we have two type II configurations interspersed, as depicted in Figure 3E. The type I filament contains 10 beads with a total length of 1.5 μm, while type II contains 8 beads and a length of 1.2 μm.

To facilitate the simulation, each bead was labeled with one of three different numerical marks. All beads that make up the cell membrane are marked as 0. The beads forming the actin filament and nuclear membrane were marked as 1. The ends of actin filaments near the cell membrane were labeled as 3. The elastic constants responsible for the bonds between each bead can be separated into 3 types: the elastic constant between the beads that make up the membranes (referred to as $k_{00}$); the elastic constant that forms the actin filaments ($k_{11}$), except the head next to the cell membrane; and the elastic constant between the head of the filament and the nearest membrane bead ($k_{13}$). The relation between the constant modules in the simulation is given by: $k_{11} = k_{00}$ and $k_{13} = 5k_{11}$.

**II. Potential Model:**

To calculate the forces capable of creating the basic structure of a cell, it is necessary to define potential of attraction and repulsion between the beads of type 0, 1 and 3 (see above).

We based our system on a Bead-Spring model of polymers, also known as the coarse-grained model. In this model, we have the following potentials based on corresponding equations:

(1) Lennard-Jones (L-J) potential, where the parameter r is the distance between particles (beads), while σ is the equilibrium distance, and $\epsilon$ the well depth energy.

$$U_{LJ} = \epsilon \left[ \left( \frac{\sigma}{r} \right)^{12} - 2 \left( \frac{\sigma}{r} \right)^{6} \right]$$

(2) Weeks-Chandler-Anderson (WCA), representing the repulsive part of L-J potential:

$$U_{WCA}(r) = \begin{cases} U_{LJ}(r) + \varepsilon, if r < \sigma \\ 0. \qquad\qquad otherwise \end{cases}$$

(3) Finite Extensible Nonlinear Elastic (FENE) potential, where κ is the FENE spring constant, $\Delta r_{max}$ is a maximal extension/compression, and $r_0$ is equilibrium position.

$$U_{FENE}(r) = -\frac{\kappa(\Delta r_{max})^2}{2} \log \left[ 1 - \left( \frac{r - r_0}{\Delta r_{max}} \right)^2 \right]$$

(4) Bend potential, related to torsional potential. It represents the energy associated with bending between three neighboring beads and $\kappa$ is the bend spring constant.

$$U_{BEND}(\theta) = \frac{\kappa}{2} (\cos\theta - \cos\theta)^2$$

In the following graphs, these potentials are shown as a function of the distances between beads for various values of $\kappa_{ij}$ , $\epsilon_{ij}$ , $\sigma_{ij}$ , $\Delta r_{ij}$ and $r_{ij}$, with i, j = 0, 1, 3 (types of beads, see above). All quantities are dimensionless and are represented by the symbol ”*”.

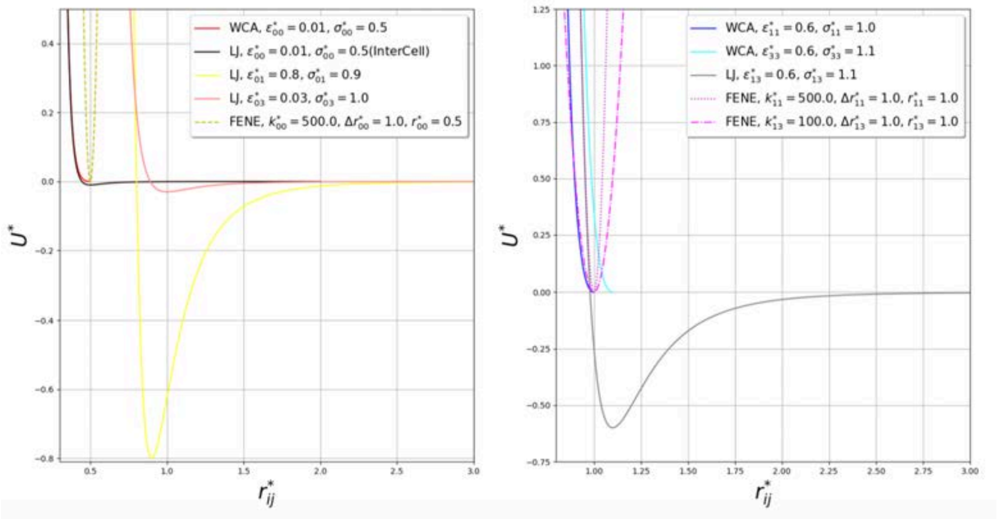


**III. AFM tip Model:**

To model the tip of an AFM, we used two elastic constants, one in x direction ($k_x$) and one in y direction ($k_y$), noting that we are working on a 2-D problem. The spring are clamped into an apparatus and the assembly moves in the direction of the cell membrane with constant velocity V. The force on the membrane is measured as a function of the deformations of the springs. The tip of the AFM is shaped like a bead and marked as 7. See details in Figure 3E.

**IV. Numerical simulation:**

With the definitions given above and appropriately choosing the parameters for σ, it is possible to analyze the stiffness behavior of the cell using the AFM tip. The proposed model allows us to attraction energy between the membranes ($\epsilon 00$) and also the attraction energy that the actin filament head makes in the membrane ($\epsilon 03$). The results of the force made by the AFM tip on the membrane as a function of its distance traveled, for both cases, are shown in graphs in Figure 3E. To facilitate analysis both graphs were plotted on the same scale.

Additional information regarding the nature of the system model is available upon request.

**Supplemental table legends:**

**Table S1. List of antibodies used for immunostaining, immunoblotting and reagents used for functional assays (Related to Figures 1-7).**

This table lists all the antibodies and reagents used for immunohistochemistry and immunocytochemistry (S1a), immunoblotting (S1b) and functional assays (S1c), including the concentration, sources, and species.

**Table S2. List of qRT-PCR primers (Related to Figures 4H and 5H).**

This table lists the primers of the indicated genes for qRT-PCR.

**Table S3. List of short hairpin constructs (Related to Figures 1, 4 and 6).**

This table list all the short hairpin (shRNA) sequences used in this study.

**Supplemental video legends**

**Movie S1. Live-cell imaging of colony formation of hESCs (Related to Figures 2 and S3).**

Lentiviral β-actin-GFP labeled hESCs in adherent culture after fresh passage at low density (mTeSR1 medium containing 2.5 μM of Thiazovivin) imaged every 2 hour over three days. Note that *PLXNB2* OE cells showed low propensity for self-assembly and did not survive after plated at low density.

**Movie S2. Live-cell imaging of F-actin dynamics in the indicated hESC colonies (Related to Figure 3).**

WT, *PLXNB2* KO and OE hESCs labeled with Lifeact_mScarlet were plated at $2 \times 10^4$ cells in Matrigel-coated coverslips and imaged after 3 days of culture at 20 min interval over 6 hr. Arrow points to filopodia-like protrusion from colony edge in the KO colony, and arrowhead points to angular cell morphology with straight junctional borders in the OE colony. Also note the differences in cortical F-actin distribution and cell morphology of mutant vs. WT colonies.

**Movie S3. Live-cell imaging of cell morphology and cellular junctions in the indicated hESC colonies (Related to Figures 3 and S4).**

WT, *PLXNB2* KO and OE hESCs labeled with CellMask were plated at $2 \times 10^4$ cells in Matrigel-coated coverslips and imaged after 3 days of culture at 3 min interval over 1 hr. Arrow points to wavy junction borders and filopodia-like protrusions from colony edge in the KO colony, and arrowhead points to a big and polygonal-shaped cell in the OE colony. Note the rounded cell morphology and cobble-stone-like cellular organization in WT hESC colony.

**Movie S4. Live-cell imaging of cell morphology and cellular junctions in the indicated hESC colonies after Blebbistatin treatment (Related to Figures 3 and S4).**

WT, *PLXNB2* KO and OE hESCs labeled with CellMask were plated at $2 \times 10^4$ cells in Matrigel-coated coverslips, and after 3 days of cell culture, cells were treated with Blebbistatin (10$\mu$M) for 3 hr and immediately imaged every 3 min over 1 hr. Note filopodia-like protrusions from hESCs in the KO colony (arrow) and disruption of epithelial arrangement in all three groups after Blebbistatin treatment, with cells aggregated into small clusters (arrowheads), signifying reduced traction forces in epithelial colonies.

**Movie S5. Mathematical simulation of mechanical rigidity of cell with constant membrane-membrane attraction energy and variable attraction energy of actin filament head and cell membrane (Related to Figure 3).**

The videos show evolution of cellular arrangement of four cells. When the system reaches equilibrium, the AFM tip (green dot on the right) begins to indent cell membrane. The membrane-membrane attraction energy ($\epsilon00$) was fixed at 0.01 and the attraction energy of actin filament head and cell membrane ($\epsilon03$) was variable (0.03, 4.00 and 30.0). Note that the total distance traveled by the AFM tip was around 60 μm, far beyond the tip displacement shown in the graph in Figure 3E, which ranged from 0 to 3 μm, similar to the AFM experiment. The beads represent actin filaments (red), actin filament head (gray), cell membrane (yellow), and AFM tip (green).

**Movie S6. Mathematical simulation of mechanical rigidity of cell with constant attraction energy of actin filament head and cell membrane and variable membrane-membrane attraction energy (Related to Figure 3).**

The videos show evolution of cellular arrangement of four cells. When the system reaches equilibrium, the AFM tip (green dot on the right) begins to indent cell membrane. The attraction energy of actin filament head and cell membrane ($\epsilon03$) was fixed at 0.03 and the membrane-membrane attraction energy ($\epsilon00$) was variable (0.01, 1.0 and 10.0). Note that the total distance traveled by the AFM tip was around 60 μm, far beyond the tip displacement shown in the graph in Figure 3E, which ranged from 0 to 3 μm, similar to the AFM experiment. The beads represent actin filaments (red), actin filament head (gray), cell membrane (yellow), and AFM tip (green).

**Movie S7. Live-cell imaging of F-actin dynamics in the indicated hNPCs (Related to Figure 5).**

Human NPCs derived from WT, *PLXNB2* KO and *PLXNB2* OE hESCs labeled with Lifeact_mScarlet were plated at 6 x $10^4$ cells in Matrigel-coated coverslips and imaged after 3 days of cell culture at every 20 min interval over 6 hr. Note that WT cells displayed dynamic movement and abundant cortical F-actin. *PLXNB2* KO and OE hNPCs exhibited larger sizes and specific F-actin patterns.

**Movie S8. Live-cell imaging of beating cardiomyocytes derived from WT and *PLXNB2* OE hESCs (Related to Figure S9).**

Phase contrast live-cell imaging shows rhythmic beating of cardiomyocytes derived from WT and *PLXNB2* OE hESCs. Images were captured at 25 frames/sec over 8 sec. Note that *PLXNB2* OE cardiomyocytes exhibited slower and a pacemaker-like beating pattern in contrast to a more fluidic and faster beating pattern in WT.

**Movie S9. Live-cell imaging of dissociated neurons from WT and *PLXNB2* KO cerebral organoids labeled with Fluo-3-AM shows neuronal activity (Related to Figures 6 and S14).**

Fluorescent live-cell imaging of dissociated neurons from cerebral organoids labeled with fluorescent calcium indicator Fluo-3-AM, at 10 frames/millisec over 30 sec.

**Movie S10. Live-cell imaging of dissociated neurons from WT and *PLXNB2* KO cerebral organoids labeled with FluoVolt shows neuronal activity (Related to Figures 6 and S14).**

Fluorescent live-cell imaging of dissociated neurons from cerebral organoids labeled with voltage sensitive probe Fluovolt, at 10 frames/millisec over 30 sec.

#### References


1. Thomson, J.A., *et al.* Embryonic stem cell lines derived from human blastocysts. *Science* **282**, 1145-1147 (1998).

2. Ran, F.A., *et al.* Genome engineering using the CRISPR-Cas9 system. *Nat Protoc* **8**, 2281-2308 (2013).

3. Sanjana, N.E., Shalem, O. & Zhang, F. Improved vectors and genome-wide libraries for CRISPR screening. *Nat Methods* **11**, 783-784 (2014).

4. Zhao, B., *et al.* TEAD mediates YAP-dependent gene induction and growth control. *Genes Dev* **22**, 1962-1971 (2008).

5. Lancaster, M.A., *et al.* Cerebral organoids model human brain development and microcephaly. *Nature* **501**, 373-379 (2013).

6. Lancaster, M. & Knoblich, J. Generation of cerebral organoids from human pluripotent stem cells. *Nat Protoc* **9**, 2329-2340 (2014).

7. Riedl, J., *et al.* Lifeact: a versatile marker to visualize F-actin. *Nat Methods* **5**, 605-607 (2008).

8. Meijering, E., Dzyubachyk, O. & Smal, I. Methods for Cell and Particle Tracking. *Methods Enzymol* **504**, 183-200 (2012).

9. Azeloglu, E.U. & Costa, K.D. Atomic force microscopy in mechanobiology: measuring microelastic heterogeneity of living cells. *Methods Mol Biol* **736**, 303-329 (2011).

10. Grashoff, C., *et al.* Measuring mechanical tension across vinculin reveals regulation of focal adhesion dynamics. *Nature* **466**, 263-266 (2010).

11. Bachmann, M. Thermodynamics and Statistical Mechanics of Macromolecular Systems. (Cambridge University Press, 2014).

12. Binder, K. Monte Carlo and Molecular Dynamics Simulations in Polymer Science. (Oxford University Press, 1995).

Fig. S1

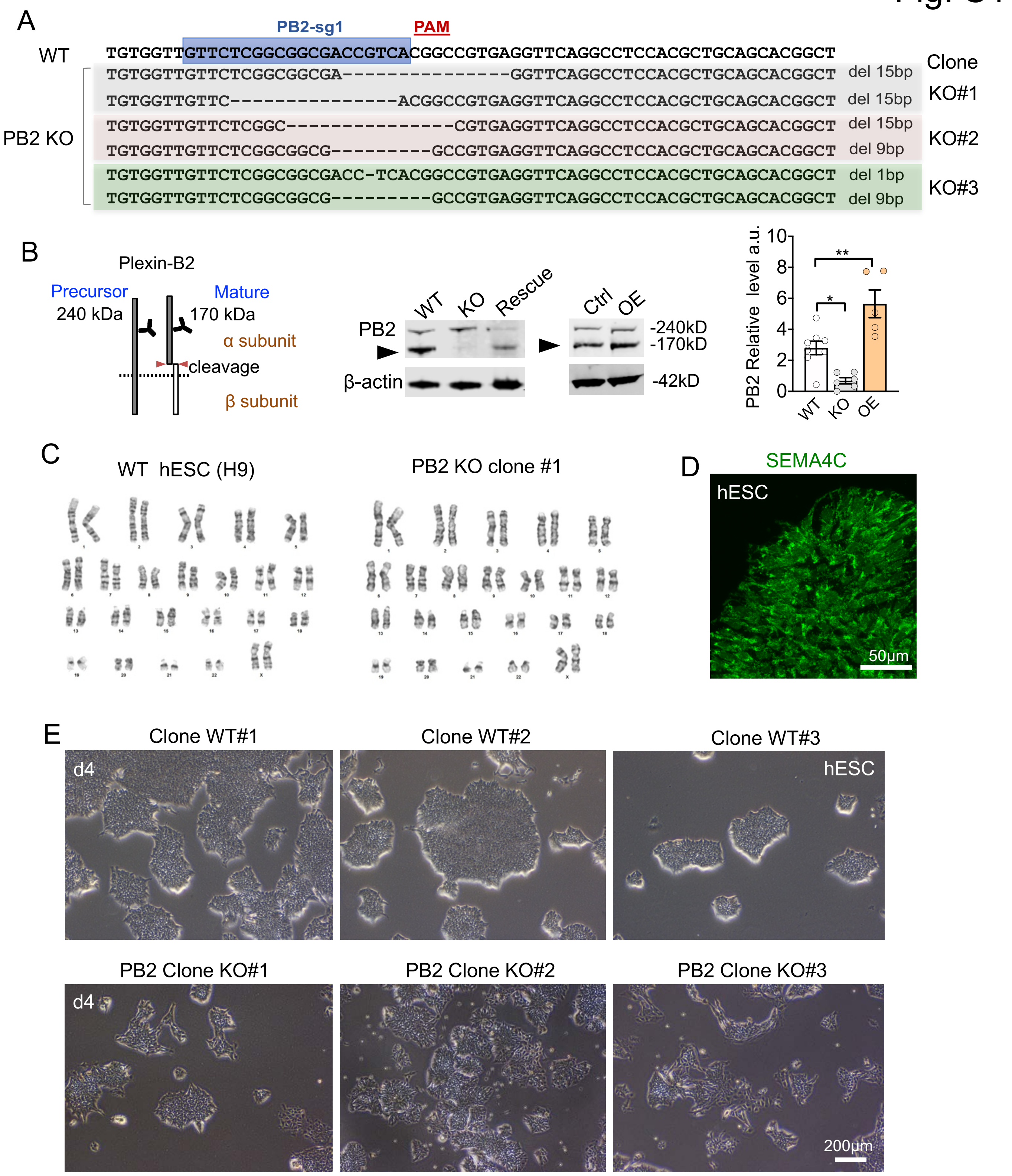


**Figure S1. CRISPR-mediated Plexin-B2 knockout results in smaller hESC colony, related to Fig. 1**

(A) Sequencing results show bi-allelic frameshift or in-frame deletion mutations in each of the three clones of *PLXNB2* KO hESC. PB2-sg, small guide RNA targeting *PLXNB2.* PAM: protospacer adjacent motif.

(B) Left, diagram of precursor (240 kD) and mature form (170 kD, α subunit after cleavage) of Plexin-B2. Middle panels, WB shows loss of mature Plexin-B2 (black arrowhead) with CRISPR-mediated knockout (PB2 KO), note the presence of mutant Plexin-B2 precursor. CRISPR-resistant *PLXNB2* rescued the expression of mature form. WB show Plexin-B2 overexpression (OE) with lentiviral vector. β-actin as loading control. Right, quantification of relative expression level of Plexin-B2. One-way ANOVA followed by Dunnett's multiple comparisons test versus WT. $F_{2,16}$= 19.45. n=5-8. *$p<0.05$, **$p<0.01$.

(C) Normal karyotypes for both wild type (WT) and *PLXNB2* KO hESC (clone #1).

(D) Immunostaining of Sema4C shows its wide expression in hESCs.

(E) Phase contrast images show smaller colony size and altered colony geometry for all three clones of *PLXNB2* KO as compared to WT clones.

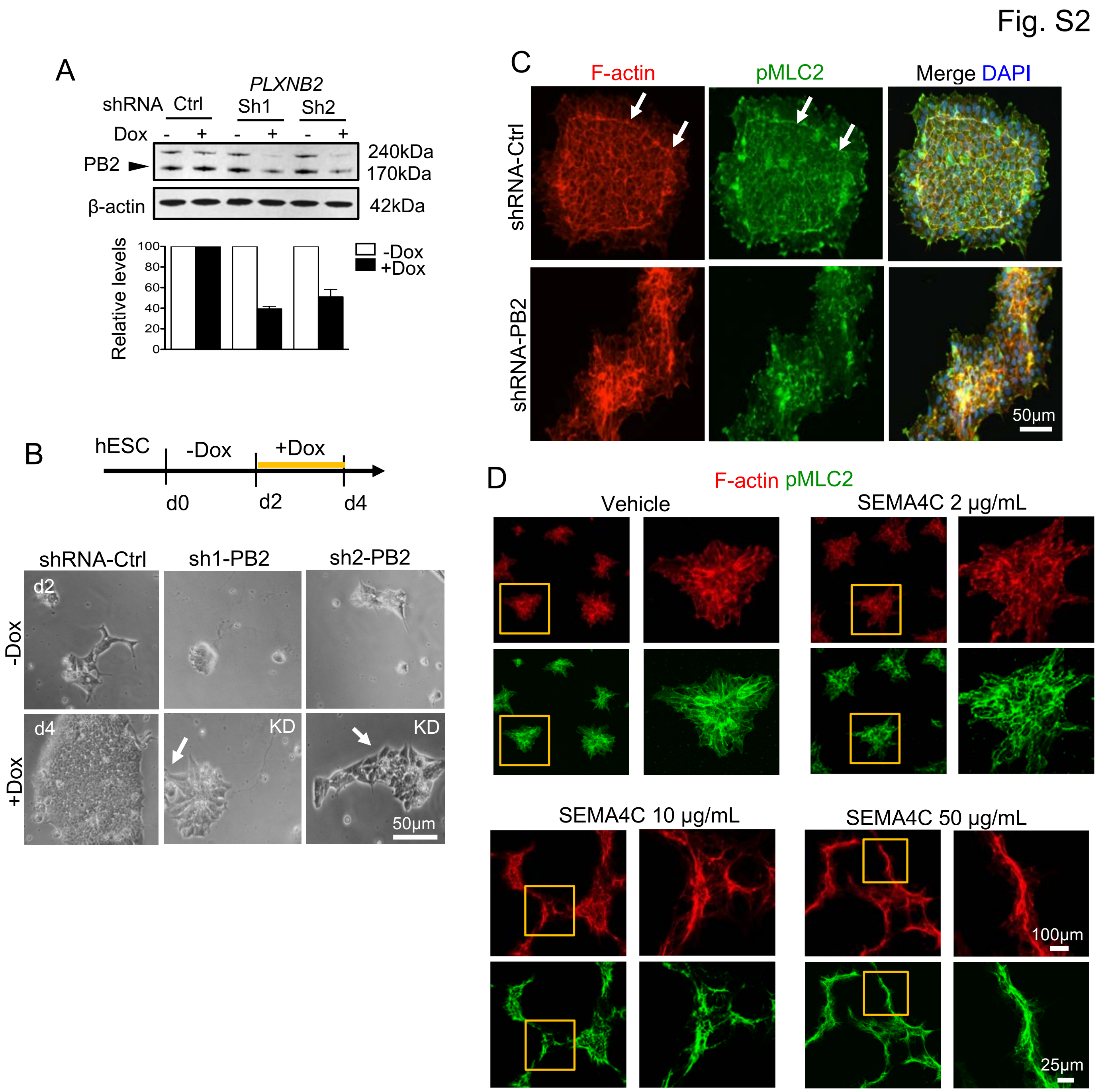


**Figure S2. Plexin-B2 regulates actomyosin network and colony geometry of hESC, Related to Figure 1**

(A) WB and quantification show Plexin-B2 knockdown with two independent Doxycycline (Dox)-inducible shRNAs. β-actin served as loading control. Black arrowhead points to mature Plexin-B2.

(B) Top, experimental scheme of Dox-induced *PLXNB2* shRNA KD in hESCs. Bottom, phase contrast images show comparable initial formation of small colonies 2 days after fresh passage of hESCs with control shRNA (Ctrl) or two Plexin-B2 shRNAs. At day 4, two days after Dox, control hESC colonies had expanded, and cells at colony edge exhibited round shape; *PLXNB2* KD resulted in slower colony expansion and altered colony geometry, with cells at colony edge exhibiting elongated morphology (arrows).

(C) Fluorescence images show actomyosin band (arrows) at periphery of WT hESC colony. *PLXNB2* KD resulted in cellular rearrangement and dissipation of actomyosin band at colony periphery.

(D) ICCs show dose-dependent effect of SEMA4C on colony geometry, F-actin (phalloidin) and pMLC2 network in hESC colonies. After fresh passage, hESCs were cultured on matrigel coating mixed with SEMA4C and analyzed four days later. Enlarged images of boxed area are shown on the right.

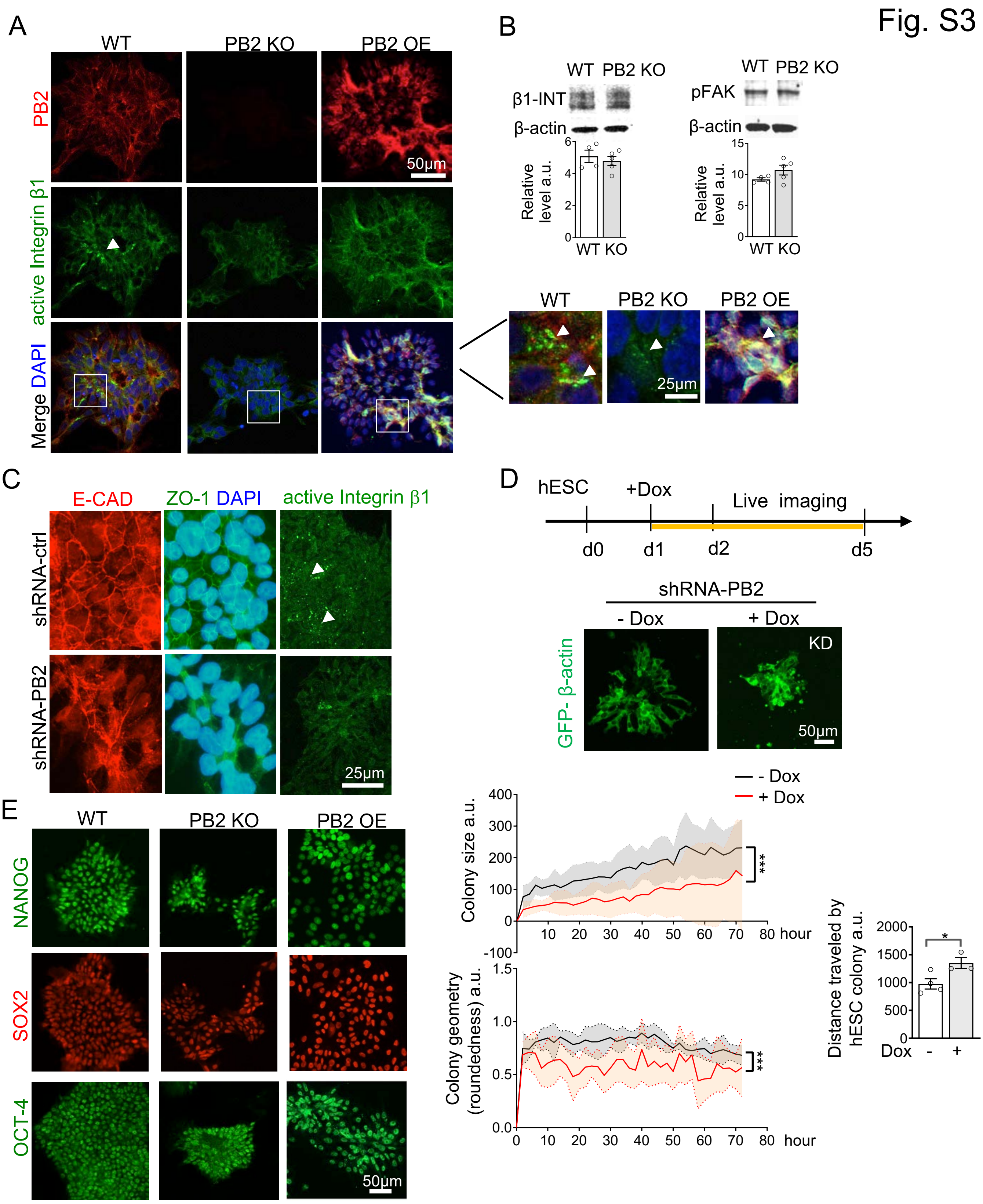


**Figure S3. Plexin-B2 stabilizes surface expression of integrin β1, Related to Figure 2**

(A) Immunostaining show juxtaposition of Plexin-B2 and activated integrin β1 (HUTS-4 antibody) in WT hESCs, both were markedly reduced or enhanced in *PLXNB2* KO or OE hESCs, respectively. Enlarged images of boxed areas are shown on the right. White arrowheads point to the dotty appearance of activated integrin β1.

(B) WB shows similar levels of total integrin β1 and pFAK in WT and *PLXNB2* KO hESCs. Graphs show mean ± SEM, a.u., arbitrary unit. Two-tailed unpaired *t*-test, n= 3-4. not significant.

(C) Confocal ICC images show junctional recruitment of E-Cadherin (E-Cad) and ZO-1 in WT hESC colony with even distribution; both were markedly reduced in *PLXNB2* KD hESC colonies, so was activated integrin β1 (HUTS-4 antibody, dotty appearance denoted by white arrowheads).

(D) Top, experimental scheme for videography of hESC colony formation under control (no Dox) or *PLXNB2* KD (+Dox) conditions, visualized by GFP-tagged β-actin. Middle: still images of hESC colonies captured from videography. Bottom, quantification of colony size, geometry, and migration. Graphs indicate mean ± SEM (shadowed area). Two-way ANOVA followed by Bonferroni's post hoc correction. For colony size (Treatment: $F_{1,216}$= 21.09, ***$P$<0.001; Time: $F_{35,216}$= 0.53; Interaction: $F_{35,216}$= 0.04), n=4 colonies per group. For colony geometry (Treatment: $F_{1,216}$= 29.39, ***$P$<0.001; Time: $F_{35,216}$= 0.20; Interaction: $F_{35,216}$= 0.12), n=4 colonies per group. Two-tailed unpaired *t*-test for colony migration, n=3-4 colonies per group. *p<0.05.

(E) ICC reveal comparable expression of stem cell markers in the indicated hESCs.

Fig. S4

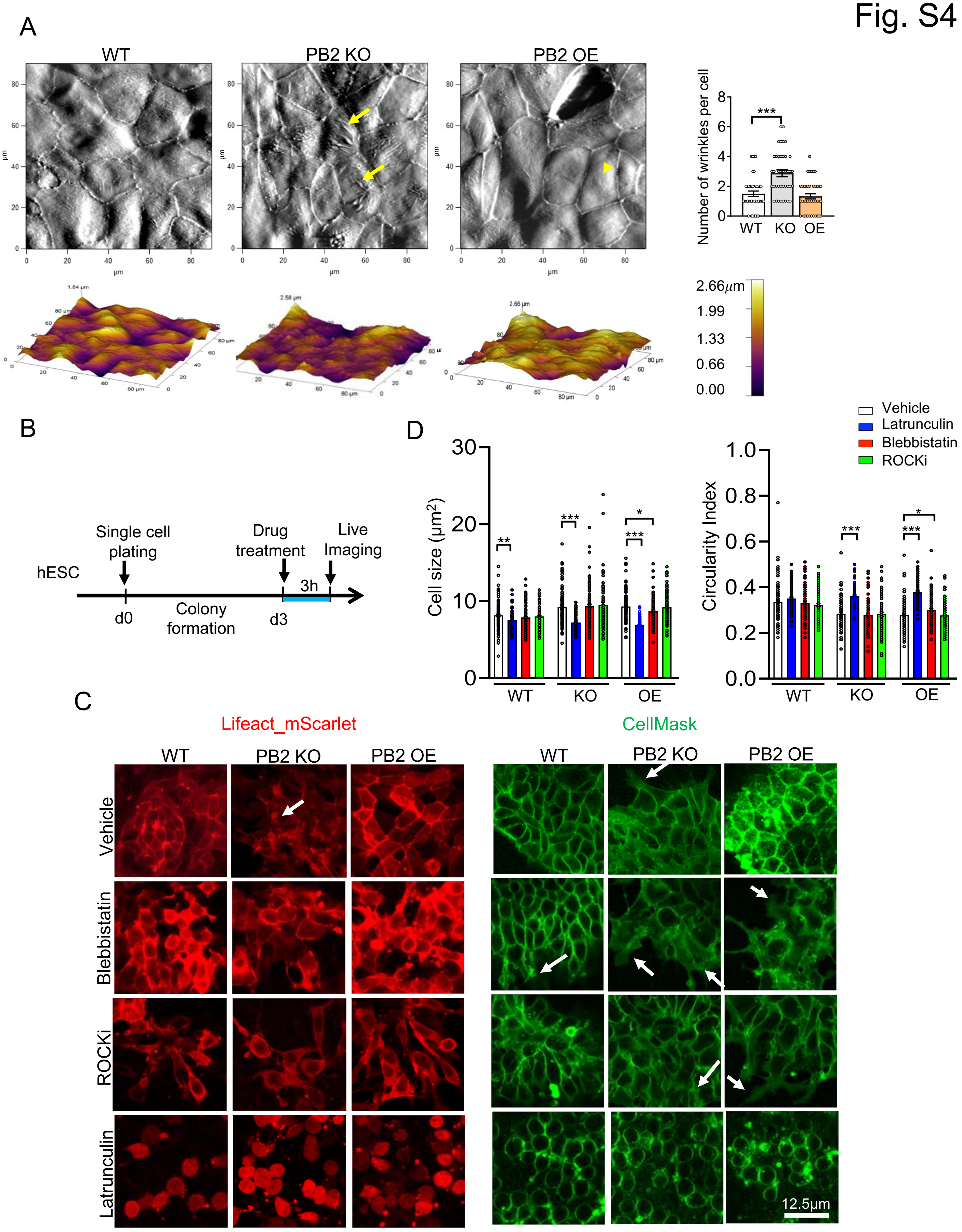
A
WT
PB2 KO
PB2 OE
Number of wrinkles per cell
***
WT KO OE
2.66μm
1.99
1.33
0.66
0.00
B
hESC
Single cell plating
Drug treatment
Live Imaging
3h
d0
Colony formation
d3
D
Cell size (μm²)
Circularity Index
Vehicle
Latrunculin
Blebbistatin
ROCKi
WT KO OE
C
Lifeact_mScarlet
CellMask
WT PB2 KO PB2 OE
Vehicle
Blebbistatin
ROCKi
Latrunculin
12.5μm

**Figure S4. Plexin-B2 regulates cell morphology and surface topography of hESCs in epithelial colony, Related to Figure 3**

(A) Top, AFM topography images show increased surface wrinkles for *PLXNB2* KO cells (arrows) in hESC colony as compared to WT or *PLXNB2* OE. Note the stretched surface of *PLXNB2 OE* cells (arrowhead). Bottom, height heatmaps show the elevation of cells relative to Z-plan. Note that *PLXNB2* KO cells are close the bottom, in contrast *PLXNB2 OE* cells are elevated, reflecting high cell stiffness. Right, graph shows mean ± SEM for number of surface wrinkles per cell. One-way ANOVA ($F_{2,117}$= 18.62) followed by Dunnett's multiple comparisons test versus WT. n=2-3 fields per group, ***p<0.001

(B) Schematics of experimental design for live-cell imaging of hESC colonies 3 hrs after drug treatment.

(C) Live-cell imaging of the indicated hESC colonies treated with vehicle or different inhibitors. F-actin network was visualized by LifeAct_mScarlet (left panels) and cell morphology and junctional borders visualized by CellMask (right panels). White arrows point to cellular protrusions from colony edge.

(D) Graphs show mean ± SEM for cell size and cell morphology. One-way ANOVA within the groups ((for cell size: WT, $F_{3,396}$= 3.54; *PLXNB2* KO, $F_{3,371}$= 23.13; *PLXNB2* OE, $F_{3,396}$= 44.26) (for cell morphology: WT, $F_{3,396}$= 3.50; *PLXNB2* KO, $F_{3,371}$= 35.19; *PLXNB2* OE, $F_{3,393}$= 63.50)) followed by Dunnett's multiple comparisons test. n=3 fields per group, *p<0.05; **p<0.01; ***p<0.001.

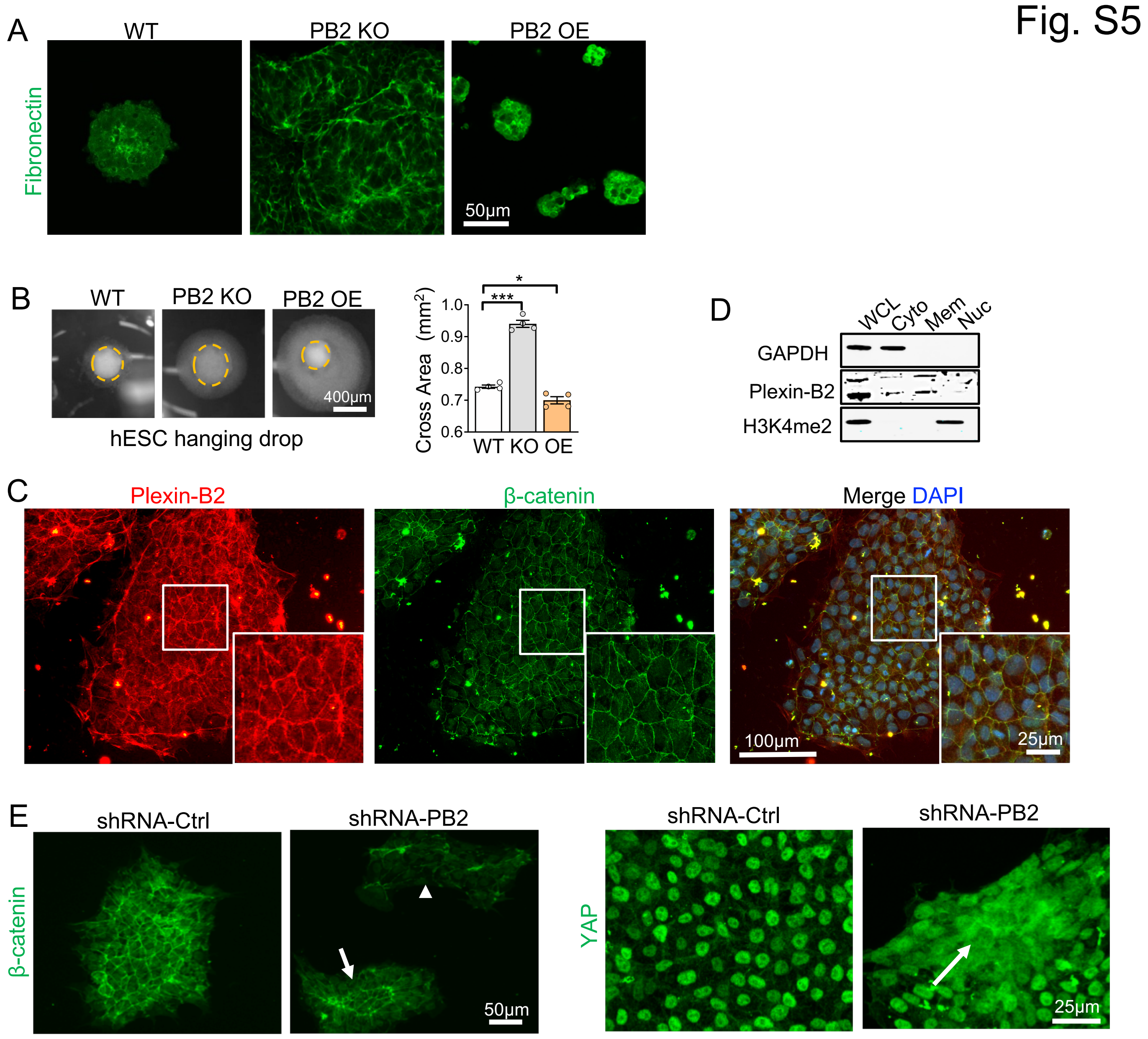


**Figure S5. Plexin-B2 regulates 3D aggregation of hESCs and influences β-catenin and YAP signaling, Related to Figures 3 and 4**

(A) Fluorescence images of 3D hESC aggregates show that *PLXNB2* KO resulted in cellular disarray and disorganized fibronectin deposits in ECM, whereas *PLXNB2* OE led to more compact fibronectin network.

(B) Images of hESC aggregates in hanging drop show reduced or increased compaction for *PLXNB2* KO or OE compared to WT, respectively. Dotted orange lines outline cell compaction. Graphs represent mean ± SEM. One-way ANOVA followed by Tukey's post-hoc test. n=4. $F_{2,9}$= 174.0. *p<0.05; *** p<0.001.

(C) ICC images of hESC colony show cell surface expression of Plexin-B2 and β-catenin in juxtaposition. Enlarged images of boxed area are shown in insets.

(D) WB verifies cell fractionation: GAPDH for cytoplasm (Cyto), Plexin-B2 for cell membrane (Mem), and Histone 3 with lysine 4 dimethylation (H3K4me2) for nucleus (Nuc). WCL, whole cell lysate.

(E) ICC images of hESC colonies show that *PLXNB2* KD resulted in lower levels of membrane β -catenin (arrowhead) and redistribution to center of cell cluster (arrow). *PLXNB2* KD also resulted in a nuclear-to-cytoplasm shift of YAP (arrow).

Fig. S6

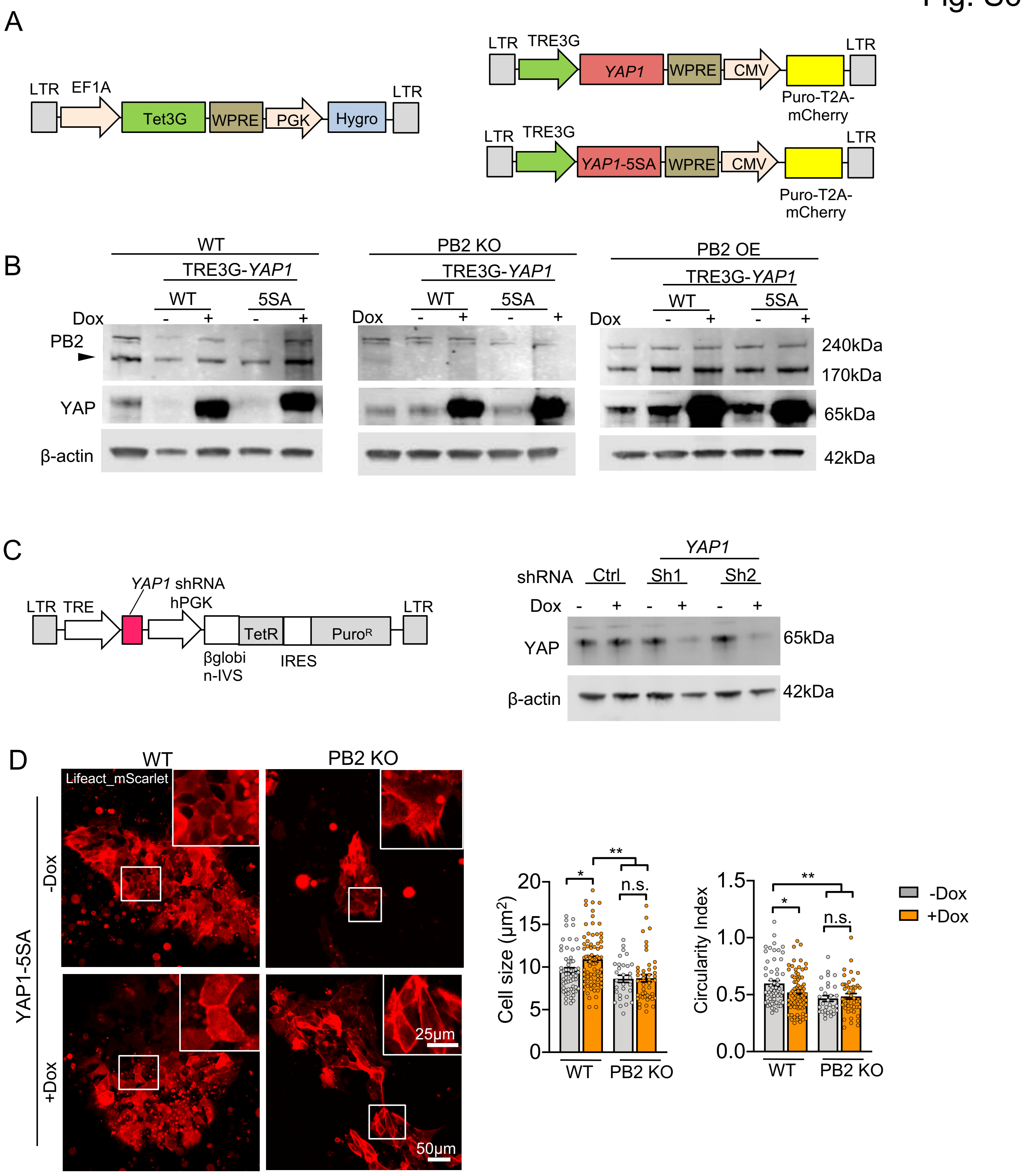
A
LTR
EF1A
Tet3G
WPRE
PGK
Hygro
LTR
LTR
TRE3G
YAP1
WPRE
CMV
Puro-T2A-mCherry
LTR
YAP1-5SA
B
WT
PB2 KO
PB2 OE
TRE3G-YAP1
WT
5SA
Dox
- + - +
PB2
YAP
β-actin
240kDa
170kDa
65kDa
42kDa
C
YAP1 shRNA
hPGK
TRE
TetR
PuroR
βglobin-IVS
IRES
YAP1
shRNA
Ctrl
Sh1
Sh2
Dox
YAP
β-actin
65kDa
42kDa
D
WT
PB2 KO
Lifeact_mScarlet
YAP1-5SA
-Dox
+Dox
25μm
50μm
Cell size (μm²)
Circularity Index
n.s.
*
**
WT
PB2 KO
-Dox
+Dox

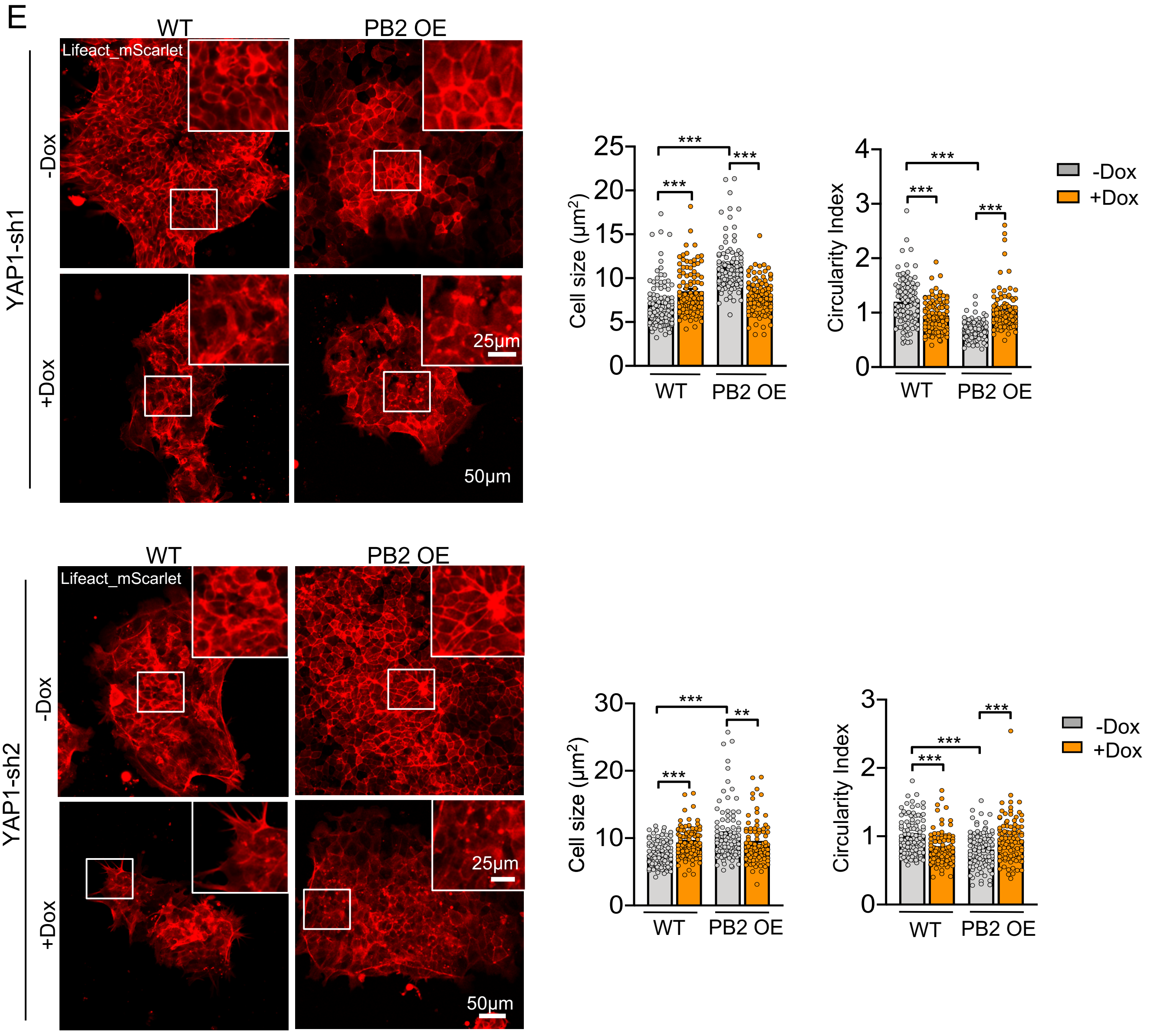


**Figure S6. YAP functions as a downstream effector of Plexin-B2 for mechanoregulation in hESC, Related to Figure 4**

(A) Schematic diagram of lentiviral constructs for Dox-inducible expression of WT (*YAP1*) or mutant YAP (*YAP1-5SA*) using tetracycline-controlled Tet-On gene expression systems Tet3G: Tet-On 3G transactivator protein. TRE3G: Tetracycline response element.

(B) WB on lysates from the indicated hESCs for Dox induced expression of WT (*YAP1*) or mutant YAP (*YAP1-5SA*). Black arrow points to mature Plexin-B2. β-actin as loading control.

(C) Left, schematic diagram of lentiviral construct for Dox-inducible expression of shRNA against *YAP1*. Right, WB shows YAP knockdown by two different shRNA.

(D-E) IF images of F-actin patterns (visualized by Lifeact_mScarlet) show different cortical F-actin, cell morphology, cell borders, and colony geometry of the indicated hESCs with manipulation of Plexin-B2 and YAP. Right, scatter dot plots of cell size and cell morphology with mean ± SEM. One-way ANOVA with Tukey's post hoc test. n=at least 3 fields per groups. For YAP1-5SA: cell size, $F_{3,207}$= 8.09; circularity index, $F_{3,207}$= 5.70. For YAP1-sh1: cell size, $F_{3,396}$= 53.13; circularity index, $F_{3,396}$= 41.98. For YAP1-sh2: cell size, $F_{3,396}$= 16.25; circularity index, $F_{3,396}$= 13.57. *p<0.05; **p<0.01; ***p<0.001.

Fig. S7

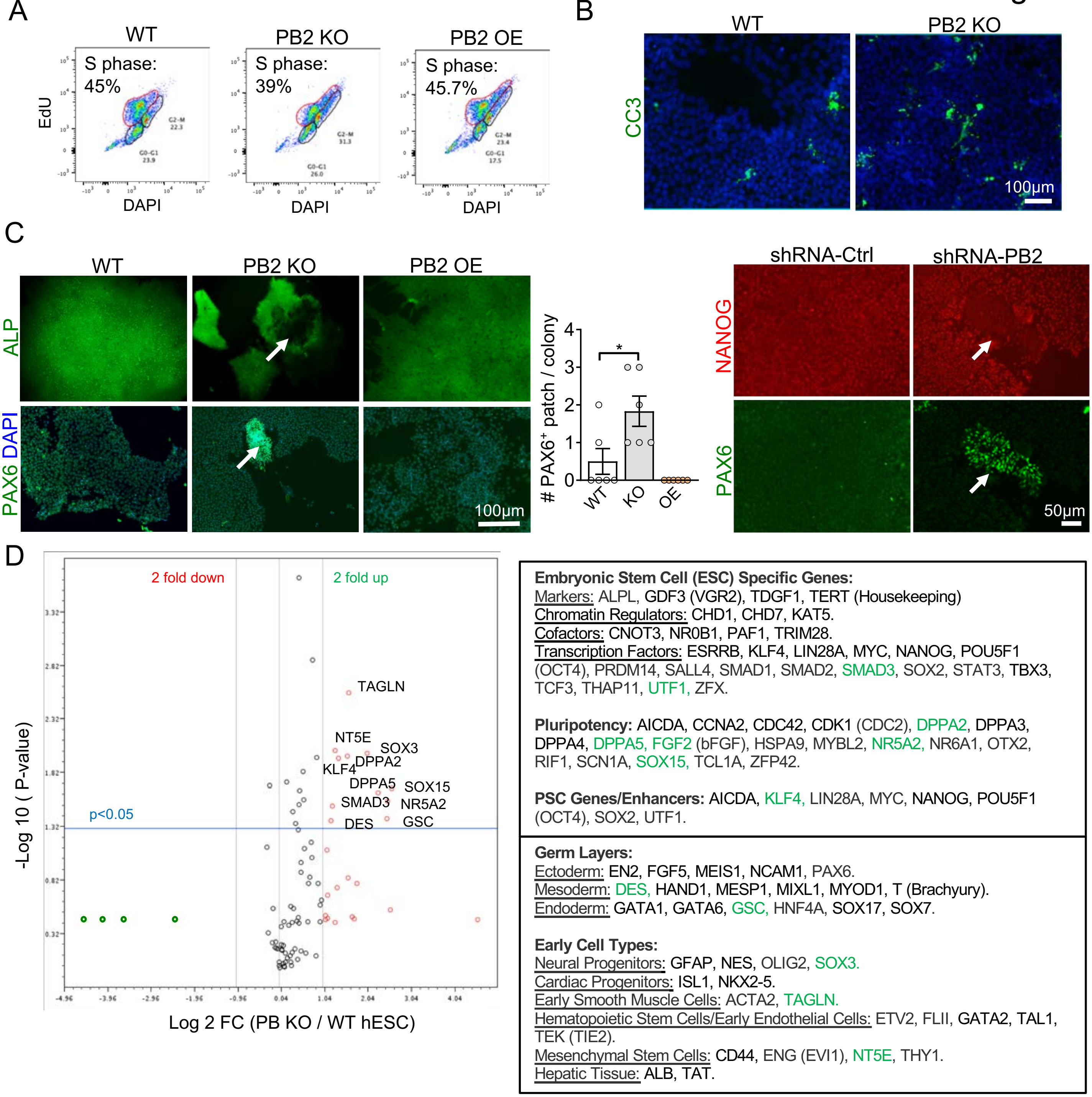


**Figure S7. Plexin-B2 activity influences hESC proliferation and differentiation, Related to Figure 4**

(A) FACS shows reduced fraction of hESCs in S phase for *PLXNB2* KO compared to WT or *PLXNB2* OE.

(B) Comparable apoptosis (cleaved caspase 3, CC3) in WT vs. *PLXNB2* KO hESCs.

(C) ICC images and quantification reveal that small patch of cells in hESC colony with *PLXNB2* KO (left) or KD (right) exhibited loss of alkaline phosphatase (ALP) signal or NANOG expression, but gain of PAX6 expression (white arrows). Graphs indicate mean ± SEM. One-way ANOVA followed by Tukey's post hoc test. n=6-8 randomly selected fields per group. $F_{2,15}$= 9.70. *p<0.05.

(D) Left, volcano plot of qRT-PCR array data displays differentially expressed genes (DEGs, >=2 fold change, p-value < 0.05). X axis, $\log_2$ fold change (FC). Y axis, p-value in -log10. Right, list of marker genes tested in qRT-PCR array. DEGs are labeled in green, denoting increased transcription.

Fig. S8

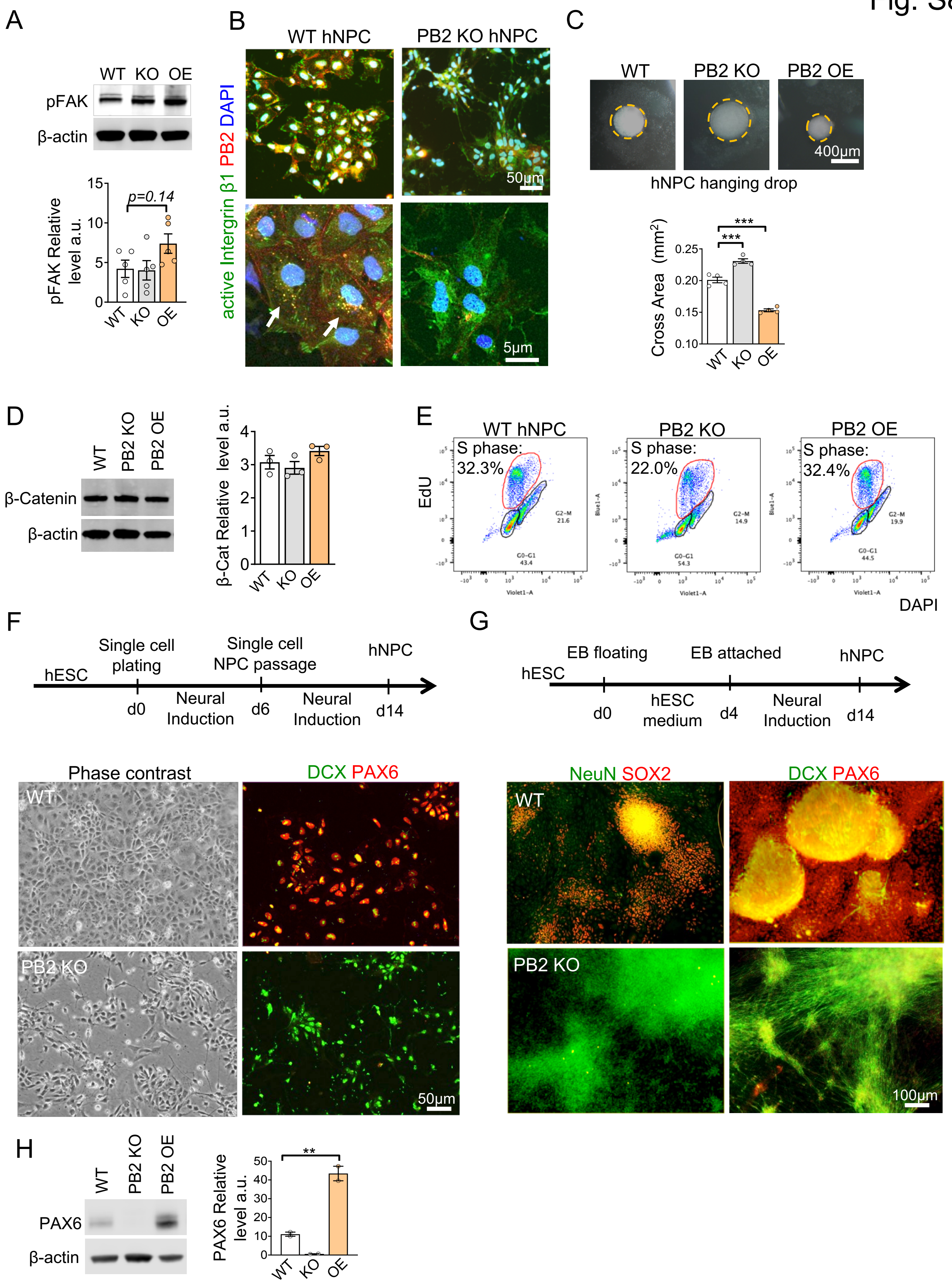
A
WT KO OE
pFAK
β-actin
pFAK Relative level a.u.
p=0.14
B
WT hNPC
PB2 KO hNPC
active Intergrin β1 PB2 DAPI
50μm
5μm
C
WT
PB2 KO
PB2 OE
400μm
hNPC hanging drop
Cross Area (mm2)
***
***
D
WT
PB2 KO
PB2 OE
β-Catenin
β-actin
β-Cat Relative level a.u.
E
WT hNPC
PB2 KO
PB2 OE
S phase: 32.3%
S phase: 22.0%
S phase: 32.4%
EdU
DAPI
F
hESC
Single cell plating
Single cell NPC passage
hNPC
d0
Neural Induction
d6
Neural Induction
d14
Phase contrast
DCX PAX6
WT
PB2 KO
50μm
G
hESC
EB floating
EB attached
hNPC
d0
hESC medium
d4
Neural Induction
d14
NeuN SOX2
DCX PAX6
WT
PB2 KO
100μm
H
WT
PB2 KO
PB2 OE
PAX6
β-actin
PAX6 Relative level a.u.
**

**Figure S8. Plexin-B2 influences hNPC proliferation and differentiation, Related to Figure 5**

(A) WB shows levels of pFAK in the indicated hNPCs. Quantification shown at bottom. One-way ANOVA followed by Dunnett's multiple comparisons test. n=5. $F_{2,12}$= 2.56. not significant.

(B) Confocal ICC images of hNPCs derived from hESCs (low magnification at top, and high magnification at bottom). *PLXNB2* KO led to altered cell morphology, size, and reduced levels of active integrin β1 (HUTS-4 antibody, dotty appearance denote by white arrows) compared to WT. Note juxtaposition of immunosignals for Plexin-B2 and active integrin β1 in WT hNPCs.

(C) Images and quantification of hNPC aggregates in hanging drop show reduced or increased compaction with *PLXNB2* KO or OE compared to WT, respectively. Dotted orange lines outline cell compaction. Graphs represent mean ± SEM. One-way ANOVA followed by Tukey's post-hoc test. n=4. $F_{2,9}$= 121.8. ***p<0.001.

(D) WB show similar levels of β-catenin in the indicated hNPCs. Quantification shown on the right. One-way ANOVA followed by Tukey's post-hoc test. n=3. $F_{2,6}$= 2.100. not significant.

(E) FACS data shows reduced fraction of cells in S phase for *PLXNB2* KO compared to WT or *PLXNB2* OE hNPCs.

(F) Top, experimental scheme of neural induction in dissociated hESC cultures. Bottom, phase contrast and ICC images show induced DCX but reduced PAX6 expression in *PLXNB2* KO vs. WT cells at day 8 of differentiation.

(G) Top, experimental scheme of neural induction of hESCs through embryoid body (EB). ICC images show premature differentiation for *PLXNB2* KO cells, demonstrated by increased NeuN and DCX expression at the expense of stem cell markers SOX2 and PAX6.

(H) WB shows different levels of PAX6 in the indicated hNPCs. Quantification shown on the right. One-way ANOVA followed by Tukey's post-hoc test. n=2. $F_{2,3}$= 92.06. **p<0.01.

Fig. S9

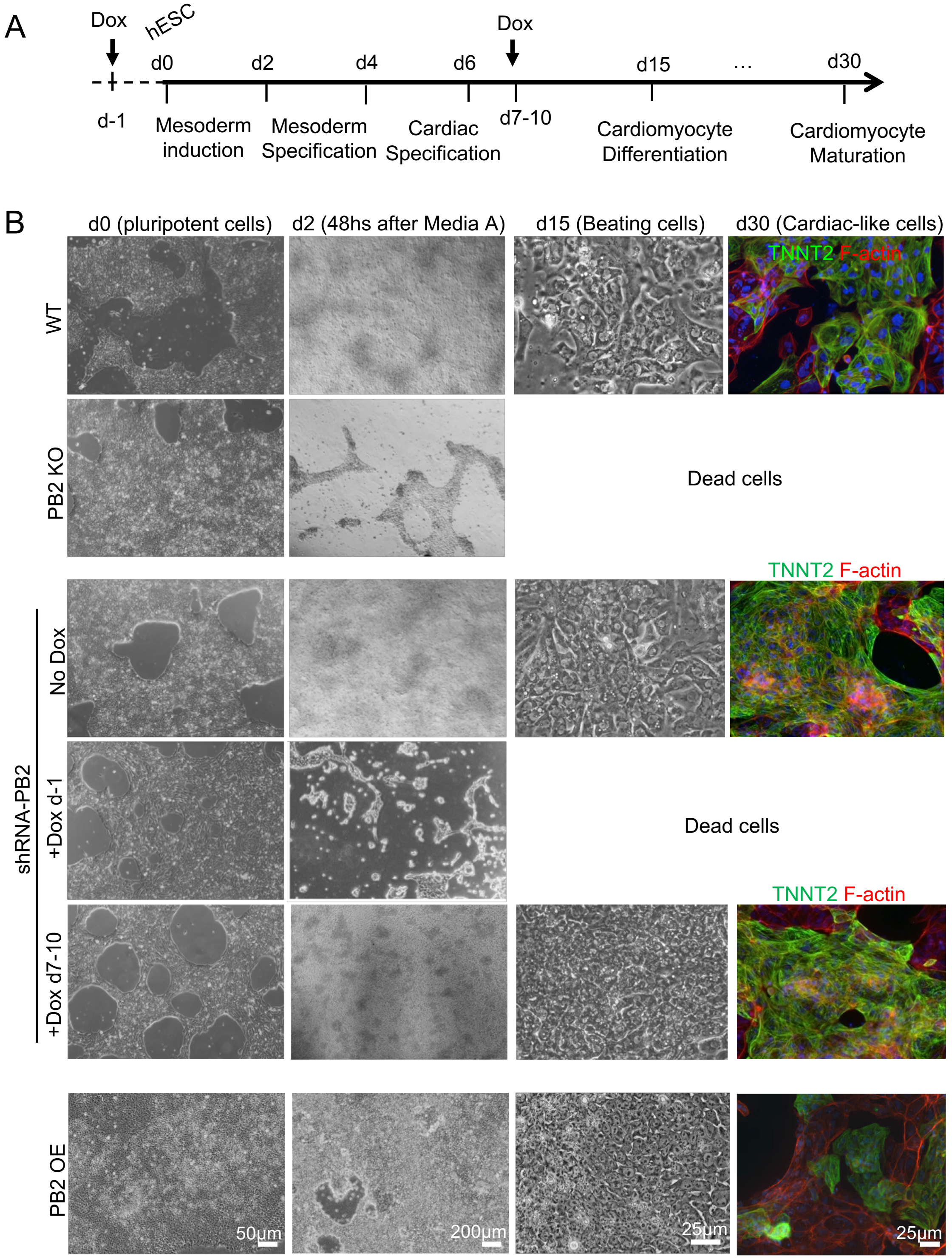


**Figure S9. Plexin-B2 activity in hESC impacts cell fate choice during mesodermal lineage specification, Related to Figure 5**

(A) Experimental scheme of mesoderm induction and cardiomyocyte differentiation from hESCs.

(B) Phase contrast and fluorescence images show mesodermal induction and cardiomyocyte differentiation from hESCs. Cardiac troponin T (TNNT2) was used as cardiomyocyte marker, and phalloidin for F-actin. hESCs with *PLXNB2* KO or early KD (Dox added at day -1) underwent cell death under identical differentiation condition. *PLXNB2* KD after mesoderm specification (Dox added at day 7-10) allowed successful cardiomyocyte differentiation. *PLXNB2* OE did not affect cardiomyocyte differentiation.

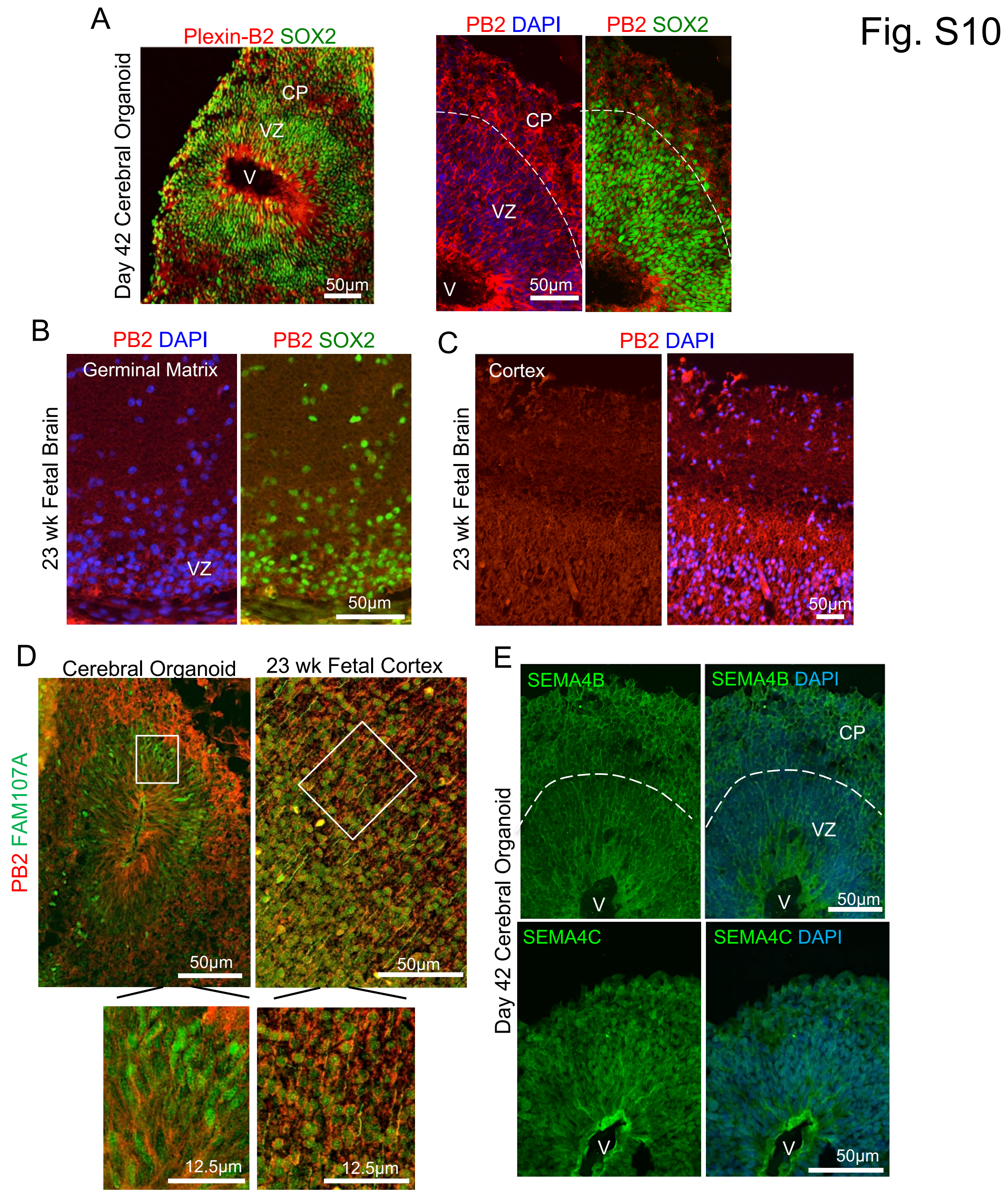


**Figure S10. Expression of Plexin-B2 and SEMA4s in the developing brains, Related to Figure 6**

(A) Left, IHC images of D42 cerebral organoid show wide expression of Plexin-B2 (PB2) in both ventricular zone (VZ) containing SOX2⁺ neuroprogenitors and cortical plate (CP). V, ventricle-like structure. Right, higher magnification images.

(B) IHC images of germinal matrix in 23-week fetal brain show co-expression of Plexin-B2 and SOX2.

(C) IHC images of developing cortex in 23-week human fetal brain show wide expression of Plexin-B2.

(D) IHC images show co-localization of Plexin-B2 with FAM107A, an outer radial glia marker, in both day 42 cerebral organoids and 23-week human fetal cortex. Enlarged images of boxed areas are shown in bottom.

(E) Expression of SEMA4B and 4C in day 42 cerebral organoids. DAPI for nuclear counterstaining.

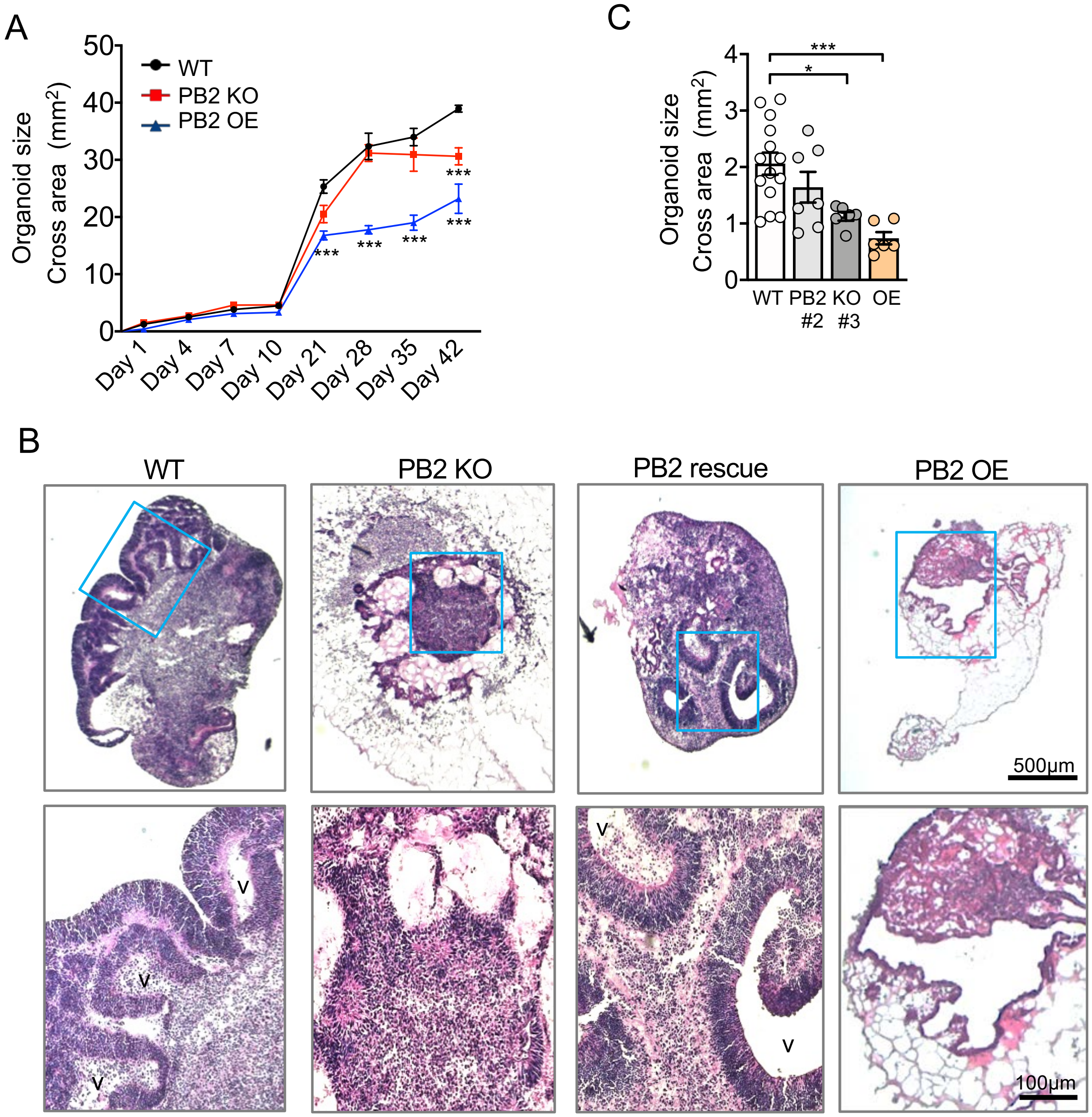


**Figure S11. Plexin-B2 activity is critical for cerebral organoid development, Related to Figure 6**

(A) Quantifications of expansion rate of cerebral organoids over the course of 42 days of differentiation. Two-way ANOVA with Bonferroni's post hoc correction. Genotype: $F_{2,95}$= 106.7, ****P*<0.001; Time: $F_{7,95}$= 477.0; Interaction: $F_{14,95}$= 15.03). n=4-8 organoids per group. ***p<0.001.

(B) H&E histology images demonstrate developmental anomaly and ventricular malformation in day 42 cerebral organoids with *PLXNB2* KO or OE. Plexin-B2 rescue construct reversed the KO phenotypes. V: ventricle-like structures. Note malformed cysts in mutant organoids. DAPI for nuclear counterstaining. Enlarged images of boxed areas are shown in bottom panels.

(C) Graphs show mean ± SEM. One-way ANOVA followed by Tukey's post-hoc test. n=6-14 organoids per group. $F_{3,29}$= 7.96. *p<0.05; ***p<0.001.

Fig. S12

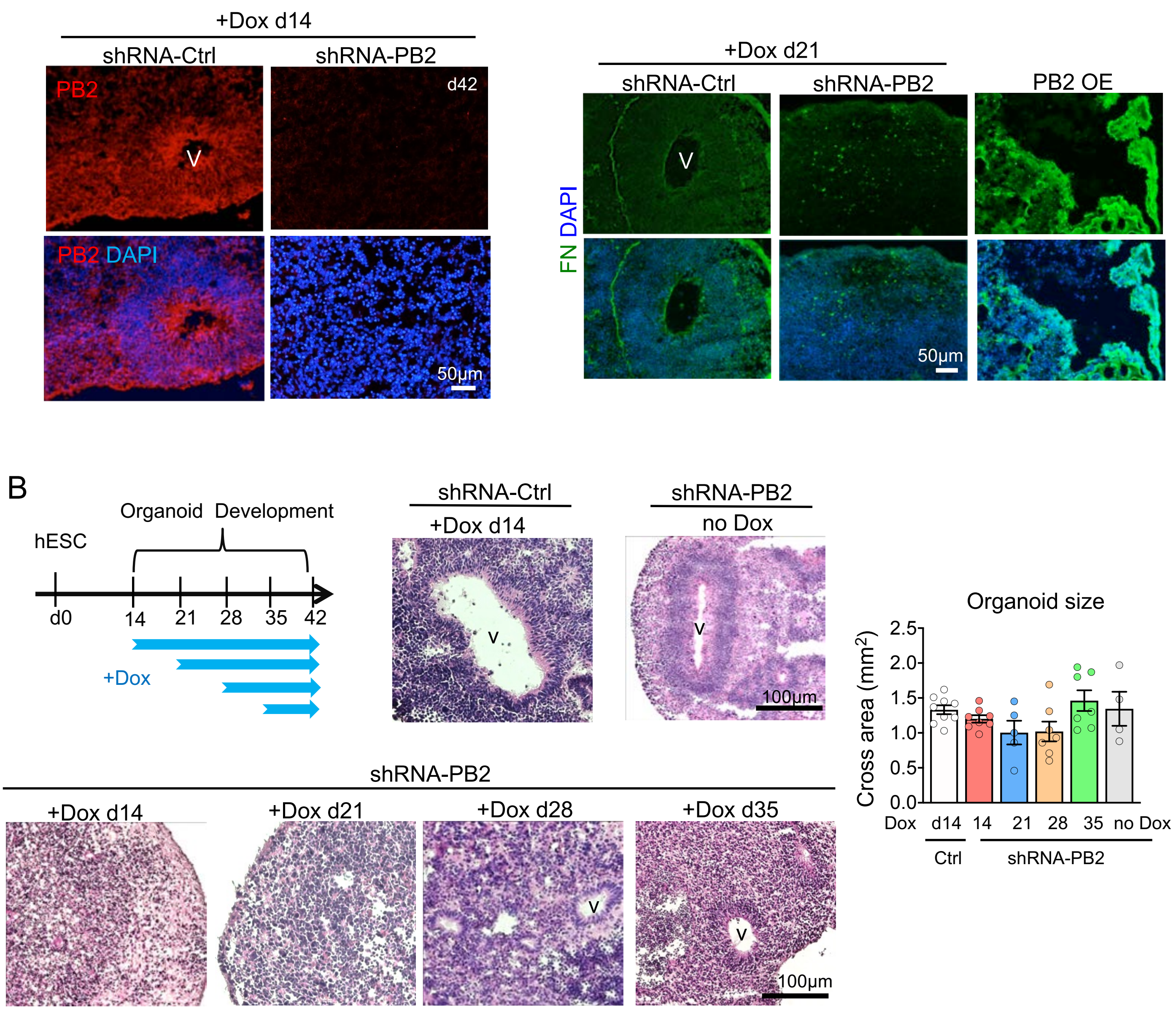


**Figure S12. Plexin-B2 deficiency or OE compromises ventricular formation and neuroepithelium cytoarchitecture in cerebral organoids, Related to Figure 6**

(A) IHC images of day 42 cerebral organoids show knockdown of Plexin-B2 with Dox added from day 14 of differentiation.

(B) Experimental scheme with *PLXNB2* KD (+Dox) at different timepoints during cerebral organoid development. H&E histology images of day 42 organoids show that the earlier the Plexin-B2 KD, the more severe the phenotypes. Cerebral organoids with control-shRNA (+Dox) or with Plexin-B2 shRNA but no Dox displayed no ventricular defects (V). Right, scatter dot plots demonstrate similar overall organoid sizes. One-way ANOVA followed by Tukey's post-hoc test. n=4-9 organoids per group. $F_{3,29}$= 7.96. not significant.

(C) IHC images demonstrate that Plexin-B2 KO or OE organoids (day 42) displayed disorganized fibronectin (FN) network along with ventricular anomaly.

### Fig. S13

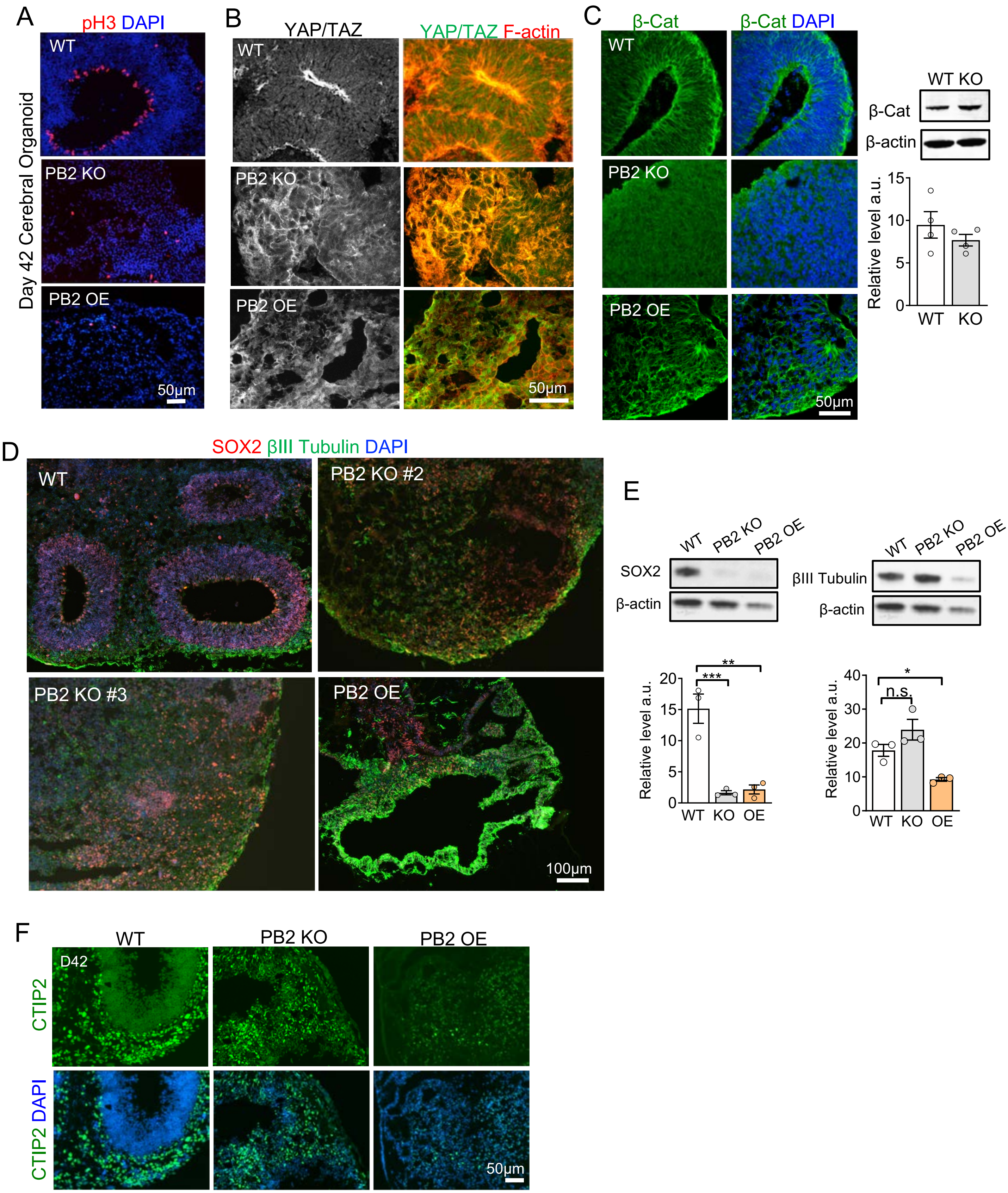

A
pH3 DAPI
Day 42 Cerebral Organoid
WT
PB2 KO
PB2 OE
50μm
B
YAP/TAZ
YAP/TAZ F-actin
WT
PB2 KO
PB2 OE
50μm
C
β-Cat
β-Cat DAPI
WT
PB2 KO
PB2 OE
50μm
WT KO
β-Cat
β-actin
Relative level a.u.
15
10
5
0
WT
KO
D
SOX2 βIII Tubulin DAPI
WT
PB2 KO #2
PB2 KO #3
PB2 OE
100μm
E
WT
PB2 KO
PB2 OE
SOX2
β-actin
βIII Tubulin
β-actin
Relative level a.u.
20
15
10
5
0
**
***
WT
KO
OE
40
30
20
10
0
*
n.s.
F
WT
PB2 KO
PB2 OE
D42
CTIP2
CTIP2 DAPI
50μm

**Figure S13. Plexin-B2 is critical for cerebral organoid development, Related to Figure 6**

(A) IHC images show reduced number and highly disorganized proliferative cells in M phase (phosphorylated Histone 3, pH3) in *PLXNB2* KO or OE cerebral organoids. In contrast, $pH3^+$ cells align apical surface of VZ in WT organoid. DAPI for nuclear counterstaining.

(B) Confocal IHC images show nuclear-to-cytoplasm shift of YAP in *PLXNB2* KO cerebral organoids as compared to WT. F-actin patterns (Phalloidin) highlight ventricular malformation in mutant organoids.

(C) Left, confocal IHC images show highly organized β-catenin (β-cat) pattern in ventricle-like structure in WT, but not in *PLXNB2* KO or OE organoids. DAPI for nuclear counterstaining. Right, WB and quantification show similar expression levels of total β-catenin in WT and *PLXNB2* KO organoids. β-actin served as loading control. a.u., arbitrary unit. Two-tailed unpaired *t*-test. n=4 organoids per group. not significant.

(D) IHC images show stereotypical layout of VZ occupied by $SOX2^+$ neuroprogenitors and outer layers containing βIII $Tubulin^+$ immature neurons in day 42 WT cerebral organoids. Organoids derived from two clones of *PLXNB2* KO hESCs (clone #2 and clone #3) or from *PLXNB2* OE hESCs contain both $SOX2^+$ and βIII $Tubulin^+$ populations, albeit in disarray. Note absence of ventricular-like structures in *PLXNB2* KO organoids, and stretched non-spherical cysts on *PLXNB2* OE organoid.

(E) WB and quantification show levels of SOX2 and βIII Tubulin in indicated cerebral organoids at day 42. a.u., arbitrary unit. One-way ANOVA with Dunnett's multiple comparisons test versus the WT group. n=3 organoids per group. For SOX2: $F_{2,6}$= 28.54. For βIII Tubulin: $F_{2,6}$= 12.91. *p<0.05; **p<0.01; ***p<0.001, n.s., not significant.

(F) IHC images show the presence of $CTIP2^+$ lower layer cortical neurons in *PLXNB2* KO cerebral organoids at day 42, as in WT organoids, albeit in disarray. *PLXNB2* OE cerebral organoids contained fewer $CTIP2^+$ neurons. DAPI for nuclear counterstaining.

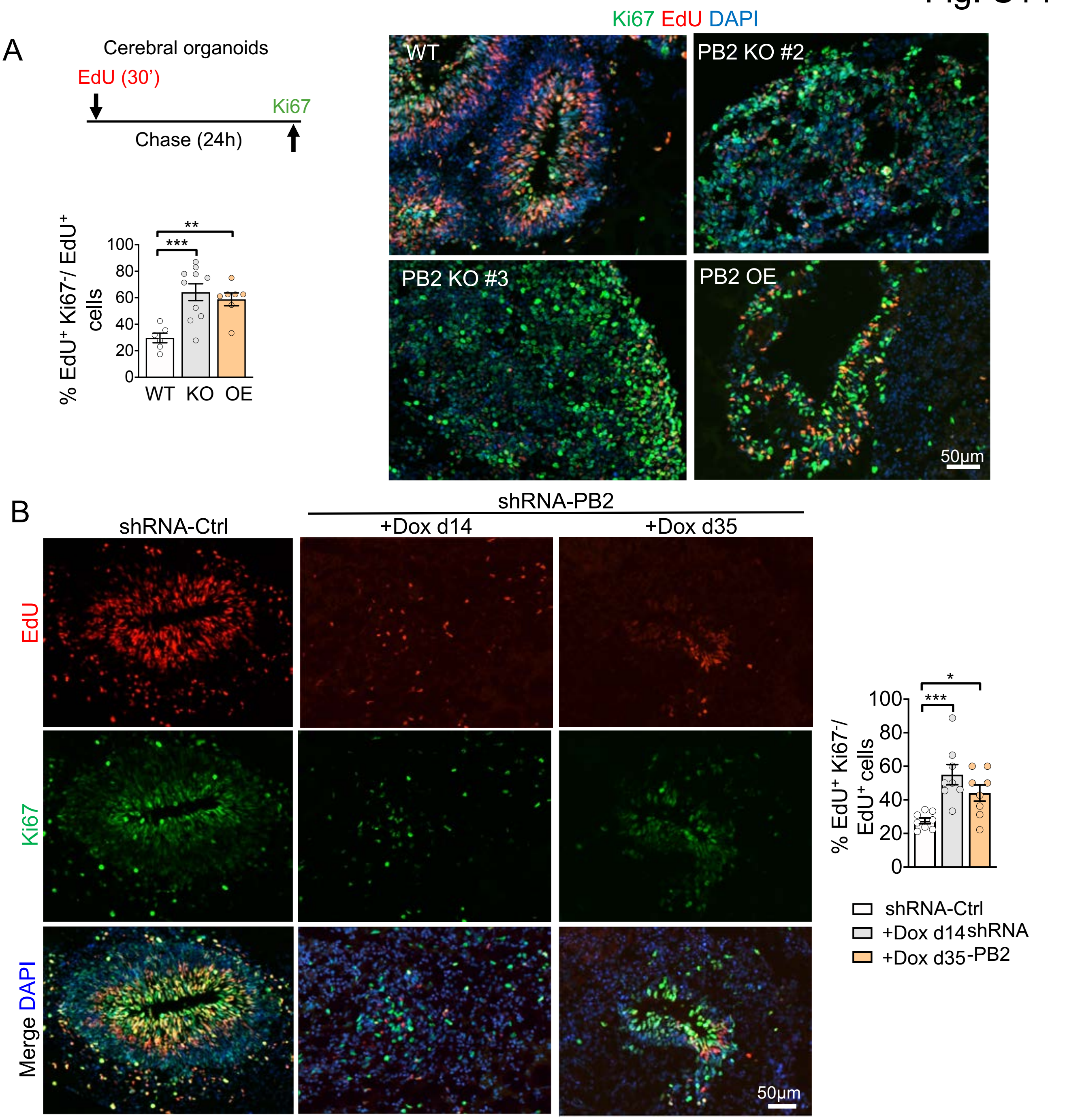


**Figure S14. Plexin-B2 regulates proliferation and cell cycle exit of NPCs in cerebral organoids, Related to Figure 6.**

(A) Experimental scheme of EdU pulse (30 min) and chase (24 hour) study. IHC images show NPCs labeled with EdU after 24 hour chase and proliferation marker Ki67 in day 42 cerebral organoids. Quantification of the percentage of $EdU^+$ $Ki67^-$/$EdU^+$ cells indicate increased cell cycle exit with *PLXNB2* KO or OE relative to WT. Graphs represent mean ± SEM. One-way ANOVA with Dunnett's multiple comparisons test versus the WT group. n=4 organoids per group (2 randomly chosen areas per organoid). $F_{2,20}$= 9.40. **p<0.01; ***p<0.001.

(B) IHC images show NPCs labeled with EdU after 24 hour chase and proliferation marker Ki67 in day 42 cerebral organoids. Dox-induced *PLXNB2* KD at the indicated time periods resulted in malformed ventricles and reduced proliferation. Quantification shows increased cell cycle exit with *PLXNB2* KD. Graphs represent mean ± SEM. One-way ANOVA with Dunnett's multiple comparisons test versus the WT group. n=4 organoids per group (2 randomly chosen areas per organoid). $F_{2,21}$= 9.33. *p<0.05; ***p<0.001.

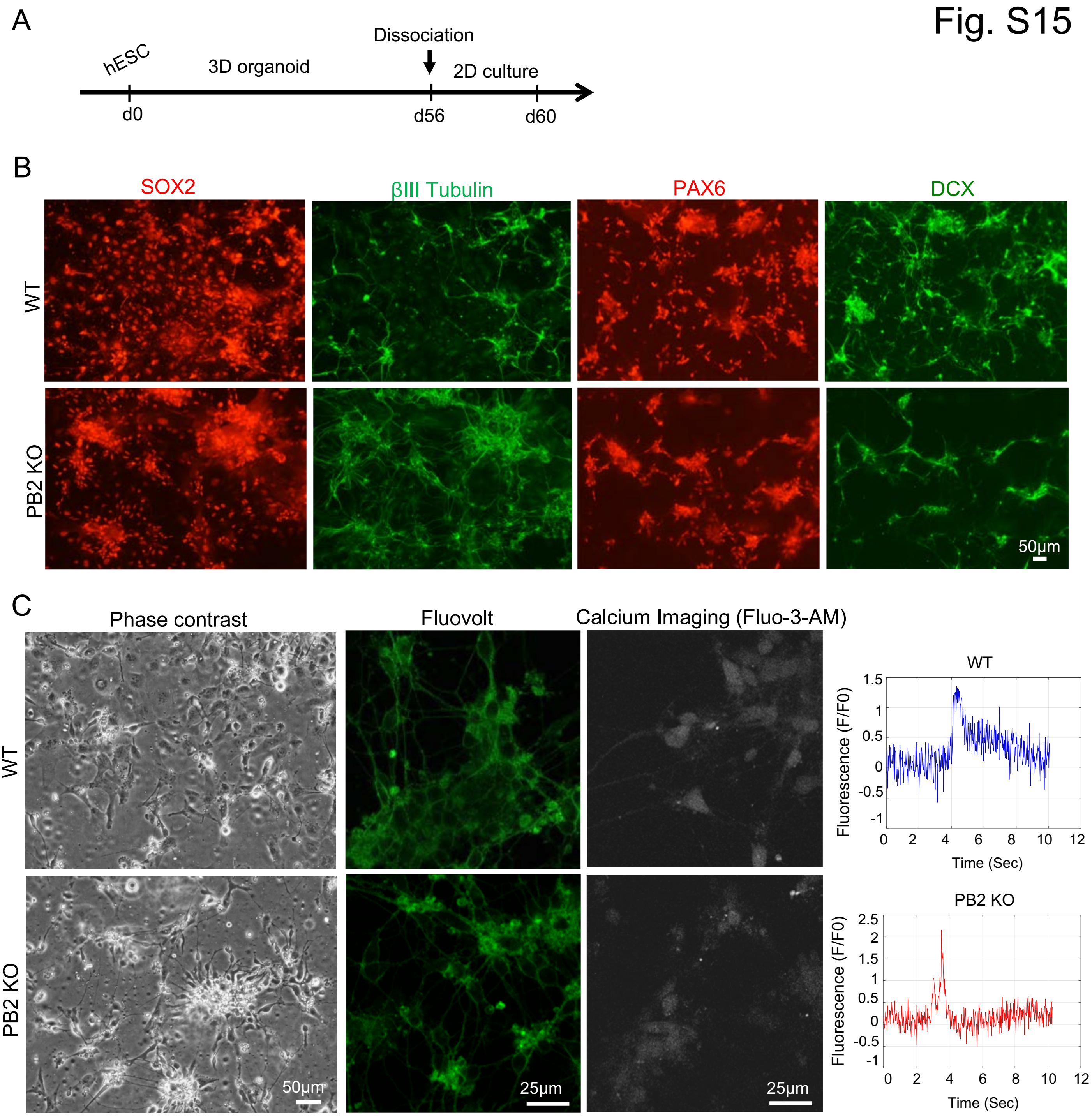


**Figure S15. Plexin-B2 deficiency accelerates neuronal differentiation and does not impair neuronal functionality, Related to Figure 6**

(A) Experimental scheme. Day 56 cerebral organoids were dissociated into single cells and cultured under adherent conditions in the same media for 4 days and analyzed at day 60.

(B) ICC images show expression of indicated markers in dissociated cells at day 60. Note increased βIII Tubulin expression in *PLXNB2* KO condition.

(C) Phase contrast images of dissociated cells from cerebral organoids (left) and still images captured from videography of action potential sparks in cells loaded with Fluovolt (right) show active neuronal activity in both conditions.

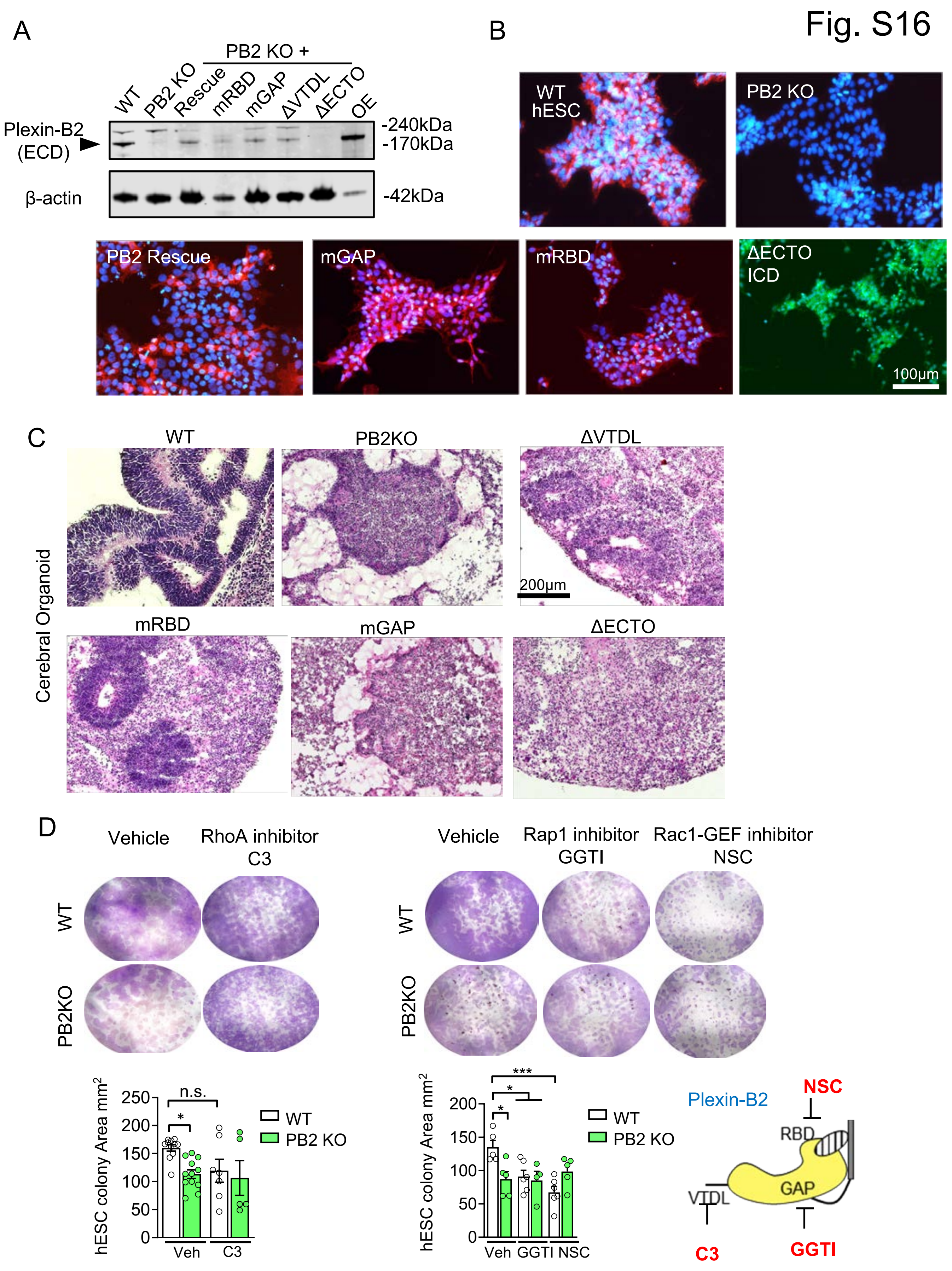
Fig. S16
A
PB2 KO +
WT
PB2 KO
Rescue
mRBD
mGAP
ΔVTDL
ΔECTO
OE
Plexin-B2
(ECD)
-240kDa
-170kDa
β-actin
-42kDa
B
WT
hESC
PB2 KO
PB2 Rescue
mGAP
mRBD
ΔECTO
ICD
100μm
C
Cerebral Organoid
WT
PB2KO
ΔVTDL
200μm
mRBD
mGAP
ΔECTO
D
Vehicle
RhoA inhibitor
C3
WT
PB2KO
Vehicle
Rap1 inhibitor
GGTI
Rac1-GEF inhibitor
NSC
hESC colony Area mm2
250
200
150
100
50
0
n.s.
*
Veh
C3
WT
PB2 KO
***
*
*
Veh GGTI NSC
Plexin-B2
NSC
RBD
GAP
VTDL
C3
GGTI

**Figure S16. Dissection of Plexin-B2 signaling domains necessary for mechanoregulation, Related to Figure 7**

(A) WB demonstrates comparable expression levels of Plexin-B2 signaling mutants. Black arrowhead points to mature Plexin-B2. Note the absence of immunosignal for Plexin-B2-ΔECTO mutant because the Plexin-B2 antibody detects epitopes in extracellular domain (ECD). Rescue denotes lentiviral expression of CRISPR-resistant Plexin-B2 in *PLXNB2* KO hESCs, while overexpression (OE) denotes lentiviral expression of WT Plexin-B2 in WT hESCs. β-actin as loading control.

(B) ICC images show expression of Plexin-B2 signaling mutants in hESCs. Note that for –ΔECTO mutant, a different Plexin-B2 antibody was used to detect the intracellular domain (ICD).

(C) H&E histology images of cerebral organoids expressing different Plexin-B2 signaling mutants in the background of *PLXNB2* KO reveal different degrees of phenotypic rescue, with Plexin-B2-ΔVTDL resulting in the best rescue, Plexin-B2-ΔECTO the least rescue, while other mutants in between.

(D) Top, images of cresyl violet staining show colony expansion of *PLXNB2* KO vs. WT hESCs. Cultures were treated with the indicated inhibitors of small G-proteins or vehicle. Graphs show mean ± SEM. One-way ANOVA followed by Tukey's post hoc test. n=7-12 wells per group. For RhoA inhibitor: $F_{3,31}$= 3.56. For Rap1 and Rac1-GEF inhibitors: $F_{5,25}$= 4.81. *p<0.05; **p<0.01; ***p<0.001; n.s., not significant. Bottom right, diagram of Plexin-B2 domains affected by the indicated inhibitors
